# Novel Class of Psychedelic Iboga Alkaloids Disrupts Opioid Use

**DOI:** 10.1101/2021.07.22.453441

**Authors:** Václav Havel, Andrew C. Kruegel, Benjamin Bechand, Scot McIntosh, Leia Stallings, Alana Hodges, Madalee G. Wulf, Mel Nelson, Amanda Hunkele, Michael Ansonoff, John E. Pintar, Christopher Hwu, Najah Abi-Gerges, Saheem A. Zaidi, Vsevolod Katritch, Mu Yang, Jonathan A. Javitch, Susruta Majumdar, Scott E. Hemby, Dalibor Sames

**Affiliations:** Department of Chemistry, Columbia University, New York, NY, USA; Department of Basic Pharmaceutical Sciences, Fred Wilson School of Pharmacy, High Point University, High Point, NC, USA; Department of Psychiatry, and Department of Molecular Pharmacology and Therapeutics, Columbia University, New York, NY, USA; Division of Molecular Therapeutics, New York State Psychiatric Institute, New York, NY, USA; Center for Clinical Pharmacology, University of Health Sciences & Pharmacy at St Louis and Washington University School of Medicine, St Louis, MO 63110, USA; Department of Neurology and Molecular Pharmacology, Memorial Sloan Kettering Cancer Center, New York, NY 10021, USA; Department of Neuroscience and Cell Biology, Rutgers University, New Jersey, NJ 08854, USA; AnaBios Corporation, 3030 Bunker Hill St., Suite 312, San Diego, CA 92109, USA; Department of Quantitative and Computational Biology, University of Southern California, Los Angeles, CA 90089, USA; Department of Chemistry, Bridge Institute, Michelson Center for Convergent Sciences, University of Southern California, Los Angeles, CA 90089, USA; Institute for Genomic Medicine, Columbia University Irving Medical Center, New York, NY, 10032, USA; Mouse Neurobehavioral Core facility, Columbia University Irving Medical Center, New York, NY, 10032, USA; The Zuckerman Mind Brain Behavior Institute at Columbia University, New York, NY, USA

## Abstract

Substance use and related mental health epidemics are causing increasing suffering and death in diverse communities.^1,2^ Despite extensive efforts focused on developing pharmacotherapies for treating substance use disorders, there is an urgent need for radically different therapeutic approaches.^3,4^ Ibogaine provides an important drug prototype in this direction, as a psychoactive iboga alkaloid suggested to have the ability to interrupt opioid use in drug-dependent humans.^5^ However, ibogaine and its major metabolite noribogaine present considerable safety risk associated with cardiac arrhythmias.^6^ We introduce a new class of iboga alkaloids - “oxa-iboga” - defined as benzofuran-containing iboga analogs and created via structural editing of the iboga skeleton. The oxa-iboga compounds act as potent kappa opioid receptor agonists *in vitro* and *in vivo*, but exhibit atypical behavioral features compared to standard kappa psychedelics. We show that oxa-noribogaine has greater therapeutic efficacy in rat models of opioid use, and no cardiac pro-arrhythmic potential, in contrast to noribogaine. Oxa-noribogaine induces long-lasting suppression of morphine and fentanyl intake after a single dose, persistent reduction of morphine intake and reinforcing efficacy after a short treatment regimen, and suppression of morphine and fentanyl drug seeking in relapse models. Oxa-noribogaine also induces a lasting elevation of neurotrophin proteins in the ventral tegmental area and medial prefrontal cortex, consistent with targeted neuroplasticity induction and alteration of addiction-like states. As such, oxa-iboga compounds represent candidates for a novel type of pharmacotherapy for treatment of opioid use disorder.

---

Ibogaine is the major psychoactive alkaloid found in the iboga plant (*Tabernanthe iboga*), a shrub native to West Central Africa.^5^ While the root bark has been harvested as a ceremonial and healing commodity in Africa for centuries, the use of iboga plant or pure ibogaine has recently become a world-wide movement with growing numbers of ibogaine healers, providers, and clinics, largely driven by the growing crises of drug addiction, trauma, despair, and spiritual starvation.^2,7,8^ Ibogaine induces profound psychedelic effects that typically include dream-like states (oneiric effects), panoramic and interactive memory recall, experiences of death and rebirth, confrontation with personal trauma, and loosening of maladaptive habits.^9^ Ibogaine is unique among other psychedelics for its ability to rapidly interrupt opioid drug dependence, as measured by dramatic reduction in opioid withdrawal symptoms.^10^ Although rigorous demonstration of clinical efficacy via controlled clinical trials is pending, the profound anti-addiction effects of ibogaine have been amply documented in anecdotal reports and open label clinical trials, including rapid and long-lasting relief of drug cravings, increased duration of abstinence, as well as long term reduction of anxious and depressive symptoms.^11–13^ The clinical claims of ibogaine’s anti-addictive properties have been recapitulated in numerous rodent models of substance use disorders (SUDs) and depression.^14–17^

Ibogaine has a complex chemical structure, where the tryptamine motif is intricately embedded in the isoquinuclidine ring, leading to a polycyclic tryptamine system that defines the iboga alkaloids (Fig. 1a, d).^18^ The pharmacology of ibogaine is equally complex, featuring a polypharmacological profile, induced by ibogaine and its main metabolite noribogaine - a dominant circulating species. The known molecular targets for both ibogaine and noribogaine include *N*-methyl-D-aspartate receptor (NMDAR, channel blocker), α3β4 nicotinic receptor (antagonist), serotonin transporter (SERT, inhibitor), and kappa opioid receptor (KOR, agonist).^19^ Mechanistically, ibogaine thus appears to bridge several different classes of psychoactive substances, including the “anesthetic psychedelics” (NMDAR blockers such as phencyclidine (PCP) or ketamine), “kappa psychedelics” (KOR agonists such as salvinorin A or U50,488), and monoamine reuptake inhibitors (such as imipramine), but shows no direct interaction with the 5-hydroxytryptamine type 2 receptors (5HT2R), setting ibogaine apart from the classical psychedelics (Fig. 1a, b). We have developed new synthetic methods for *de novo* synthesis of the iboga alkaloid scaffold, which unlocks a broad exploration of its structure and pharmacology (Fig. 1c).^20,21^ Specifically, we developed and optimized both nickel-catalyzed and palladium-catalyzed processes for 7-membered ring formation, that enables modular preparation of the benzofuran iboga analogs (Extended Data Fig. 2). These novel analogs show greatly potentiated KOR activity on the iboga pharmacological background, constituting a new class of iboga alkaloids (Fig. 1).

**Fig. 1.**
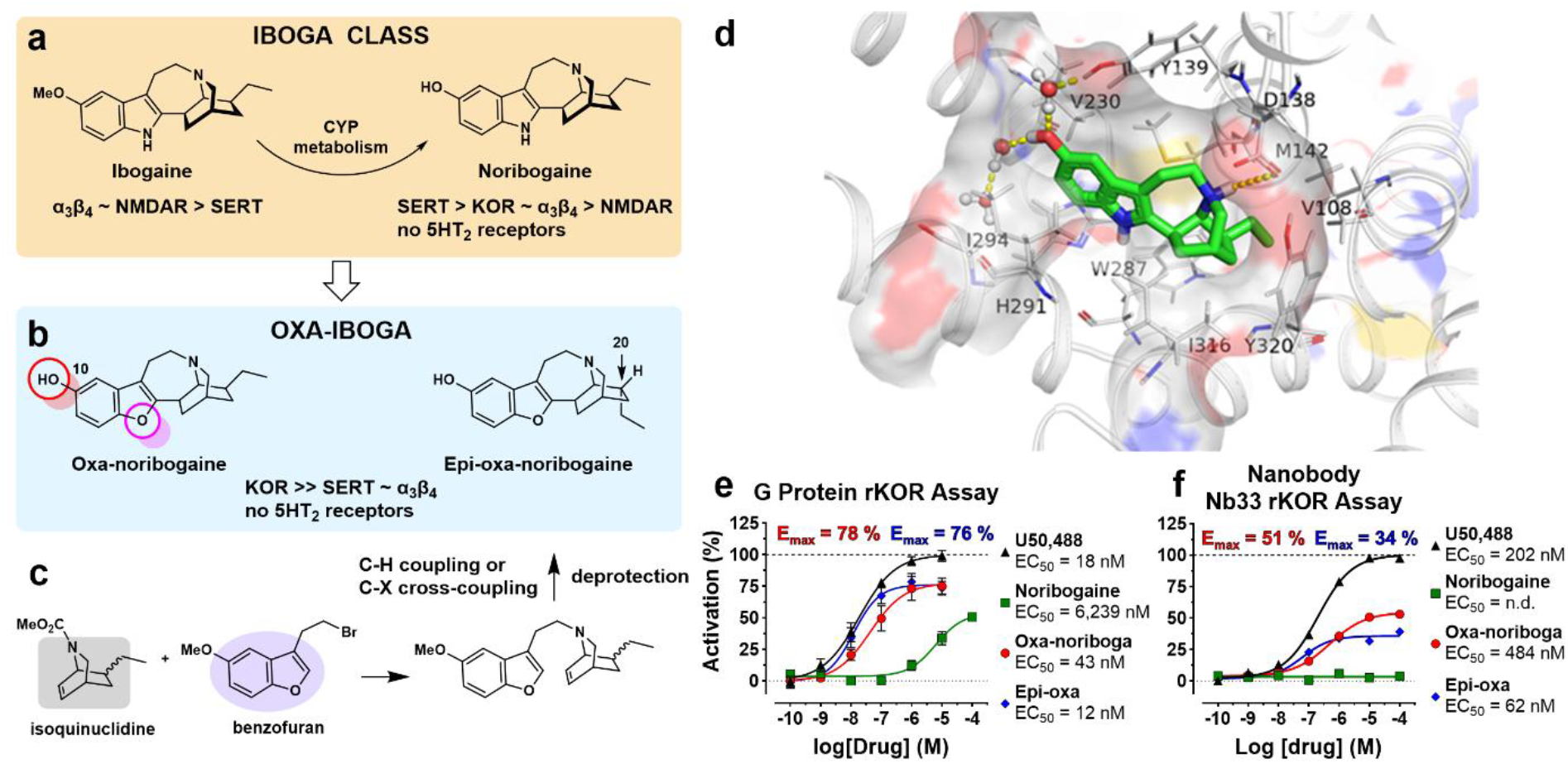
Oxa-iboga is a novel class of iboga alkaloids discovered by structural editing of ibogaine, enabled by efficient *de novo* chemical synthesis. **a**, Ibogaine is a unique probe drug with profound clinical effects and complex pharmacology, that is distinct from classical psychedelic tryptamines. Relative potencies at known molecular targets are shown. **b**, Oxa-iboga analogs are defined by the replacement of indole with benzofuran, resulting in accentuation of the KOR activity on the iboga pharmacological background. **c**, *De novo* synthesis of iboga molecular framework rests on the catalytic union of the two main structural components of oxa-iboga skeleton, the isoquinuclidine and benzofuran ring systems. **d**, Docking pose of noribogaine (carbon frame in green) in sticks representation inside KOR structure (active receptor state). Hydrogen bonding near the C10 phenol is highlighted by yellow dashed lines. **e**, Oxa-noribogaine analogs are agonists of rat KOR *in vitro*, as demonstrated by a G protein activation BRET assay. **f**, In the nanobody Nb33 sensor recruitment assay, which approximates true intrinsic signaling efficacy, oxa-noribogaine analogs are partial agonists. KOR (kappa opioid receptor), α3β4 (α3β4 nicotinic acetylcholine receptors), SERT (serotonin transporter), 5HT (serotonin), NMDAR (*N*-methyl-D-aspartate receptor). Data are presented as mean ± SEM (N = 3).

## Oxa-iboga analogs are potent KOR agonists *in vitro*

Noribogaine, the starting and comparison point for our studies, acts as a KOR partial agonist in a bioluminescence resonance energy transfer (BRET) assay for G protein activation (rat KOR, EC_50_ = 6.2 μM, E_max_ = 54%, Fig. 1e, Table S3), consistent with previous reports.^22^ We found that substitution of the indole NH group in noribogaine with oxygen dramatically accentuates KOR activity: oxa-noribogaine (oxa-noriboga, Fig. 1b) binds to the KOR (mKOR, K_i_ = 41 nM) and activates G protein in the [^35^S]GTP*γ*S assay (mKOR, EC_50_ = 49 nM, E_max_ = 92%) and BRET assay (rKOR, EC_50_ = 43 nM, E_max_ = 78%), representing a large increase in potency versus noribogaine (Fig. 1e, Table S3). Oxa-noribogaine is more than ten-fold selective for KOR versus MOR and about seven-fold over DOR in the [^35^S]GTP*γ*S binding assay using mouse opioid receptors (Extended Data Fig. 3). To further probe KOR signaling initiated by oxa-noribogaine, we used a recruitment assay based on the Nb33 nanobody that senses an active conformation of the KOR, thus limiting the signal amplification of G protein activation assays and approximating true intrinsic signaling efficacy of tested compounds (BRET assay utilizing a luciferase-tagged KOR and Venus-tagged Nb33, Extended Data Fig. 4c). It has previously been shown using the Nb33 sensor that the standard KOR agonist, U50,488, has comparable signaling efficacy to the endogenous KOR ligand, dynorphin A.^23^ In direct comparison to U50,488 as the standard, oxa-iboga compounds exhibit markedly reduced signaling efficacy in this assay (E_max(oxa-noriboga)_ = 51%, E_max(epi-oxa)_ = 34%, Fig. 1f), differentiating themselves from classical KOR agonists such as U50,488.

The phenol group in the 10-position is the essential component of both the iboga and oxa-iboga KOR pharmacophores, as agonist activity is lost if the phenol is masked or removed (Table S7). Noribogaine’s binding pose obtained by docking to the active KOR state identified water mediated hydrogen-bonding interactions between the C10 phenolic hydroxyl and tyrosine 139^3.33^ residue, as well as an ionic interaction between the isoquinuclidine amine and acidic aspartate 138^3.32^ residue (Fig. 1d). The KOR potency and selectivity can be further modulated by the inversion of the geometry of the C20 center, such as in the endo-epimer, epi-oxa-noribogaine (epi-oxa, Fig. 1b), although the effects are relatively modest (Fig. 1e).

For the rest of iboga pharmacology, SERT is arguably the most rigorously validated molecular target of noribogaine.^24,25^ The oxa-iboga compounds largely maintain 5HT reuptake inhibitory activity with a modest loss of potency compared to noribogaine (IC_50(oxa-noriboga)_ = 711 nM versus IC_50(noriboga)_= 286 nM, Extended Data Fig. 4d). Similarly, the inhibitory activity of oxa-noribogaine at the α3β4 nicotinic receptors is comparable to that of noribogaine, with a marginal increase in potency (IC_50(oxa-noriboga)_ = 2.9 μM vs IC_50(noriboga)_ = 5.0 μM, Extended Data Fig. 4e). A broad receptor screen showed a favorable pharmacological profile (Table S11) with more than 100-fold separation in binding potency between KOR and non-opioid human targets (with the exception of SERT and α3β4 molecular targets).

## Oxa-noribogaine shows potent analgesia, no aversion, and no pro-depressive-like effects

Oxa-noribogaine induces potent antinociceptive effects in male mice in the tail-flick assay (thermal nociception), with comparable potency to the standard KOR agonist, U50,488 (Fig. 2a, ED_50(oxa-noriboga)_ = 3.0 mg/kg vs ED_50(U50)_ = 2.2 mg/kg; s.c. administration). In KOR knock-out (KOR-KO) male mice, the analgesic effect is substantially attenuated, with a one-log-unit right shift in the dose response curve (ED_50(WT)_ = 2.3 mg/kg to ED_50(KOR-KO)_ = 17.7 mg/kg, Fig. 2b). Using MOR knock-out (MOR-KO) male mice, a small rightward shift in the dose-response curve was observed (Fig. 2b, ED_50(MOR-KO)_ = 7.0 mg/kg). Epi-oxa-noribogaine also induces a potent antinociceptive effect, which is abolished in KOR-KO mice, while a marginal rightward shift was observed in MOR-KO mice (Fig. 2c, ED_50(WT)_ = 1.3 mg/kg, ED_50(KOR-KO)_ = n.d., ED_50(MOR-KO)_ = 2.0 mg/kg). In wild type female mice, epi-oxa exhibits a rightward shift in potency in tail flick compared to males (ED_50(male)_ = 1.9 mg/kg vs ED_50female_) = 9.7 mg/kg, Fig. 2a,d), as does U50,488 (ED_50(male)_ = 2.2 mg/kg to ED_50female_) = 6.3 mg/kg, Fig. 2a,d), consistent with previous reports for this and related standard KOR agonists.^26,27^. In contrast, oxa-noribogaine shows a small rightward shift in the tail-flick assay in one cohort (ED_50(male)_ = 3.0 mg/kg to ED_50(female)_ = 4.9 mg/kg, Fig. 2a,d), with practically no difference in other cohorts used as controls for KOR-KO experiments (ED_50(male)_ = 2.3 mg/kg vs ED_50(female)_ = 2.1 mg/kg, Fig. 2b,e). In female KOR knock-out mice, the analgesic effect of oxa-noribogaine is greatly diminished (ED_50(WT)_ = 2.1 mg/kg vs ED_50(KOR-KO)_ = 34.3 mg/kg, Fig. 2e). Thus, the analgesic potency and efficacy of oxa-noribogaine is comparable between male and female mice and primarily driven by KOR activation in both sexes. Pharmacological studies with opioid receptor antagonists support this interpretation and show that the delta opioid receptors (DOR) do not contribute to the analgesic effect (Extended Data Fig. 5). The antinociceptive test not only indicates a desirable therapeutic-like effect, but also provides a useful physiological readout for functional KOR engagement *in vivo*, which enables dosing calibration for behavioral studies (*vide infra*).

**Fig. 2.**
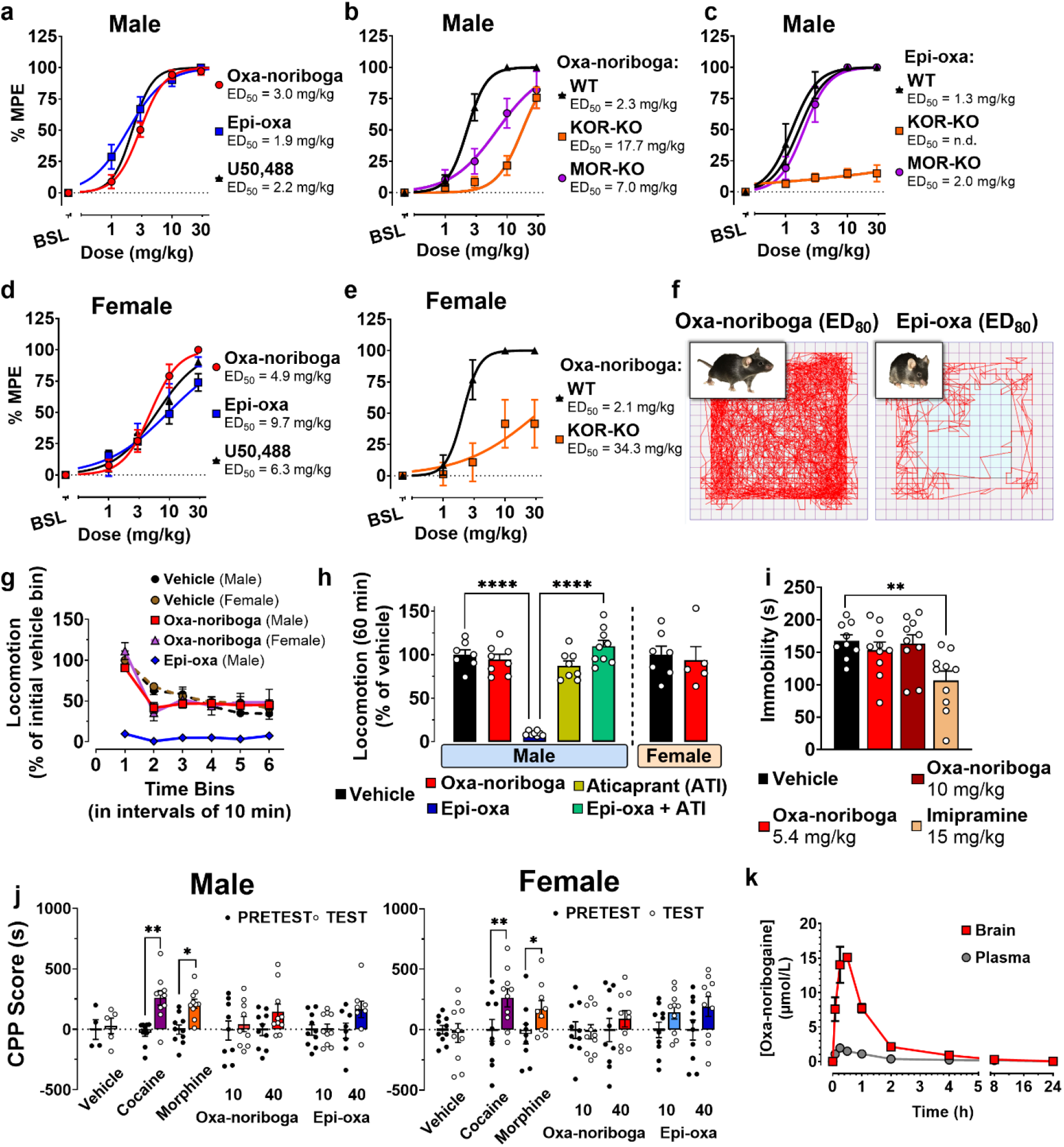
Oxa-noribogaine is an atypical KOR agonist *in vivo* with potent analgesic effect, no aversion, and no psychedelic-like effects at efficacious analgesic doses. **a**, Oxa-iboga analogs induce potent analgesia in the mouse tail-flick test (male mice), comparable in potency and efficacy to the standard kappa psychedelic, U50,488. Analgesia of oxa-noribogaine **b**, and epi-oxa-noribogaine **c**, is KOR dependent as demonstrated in KOR knock-out mice (KOR-KO), compared to mu receptor knock-out (MOR-KO) and wild type (WT) mice (KOR-KO vs WT, P < 0.0001). **d**, Oxa-noribogaine induces potent analgesia in female mice, **e**, driven by KOR as demonstrated in female KOR-KO mice. **f**, Traces visualizing ambulatory distance traveled by WT mice in open field test (OF) after drug administration show that oxa-noribogaine causes no sedation at a high analgesic dose (analgesic ED_80_, 5.4 mg/kg), in contrast to epi-oxa-noribogaine (ED_80_ = 5.2 mg/kg) which is profoundly sedative. **g**, Quantification of OF test (ED_80_ doses) for oxa-noribogaine and epi-oxa-noribogaine, data are normalized to initial locomotion of vehicle group. **h**, Sedation of epi-oxa-noribogaine (P < 0.0001) is KOR-driven as demonstrated by pre-treatment (P < 0.0001) by the selective KOR antagonist aticaprant (ATI). Total locomotion over 60 min period is normalized to vehicle. **i**, No pro-depressive-like effects were detected using the forced swim test after oxa-noribogaine administration (30 min post administration). **j**, Male and female mice do not develop a conditioned place preference or aversion (CPP/CPA) after administration of oxa-noribogaine (P=0.4531, P=0.5994, respectively). The two doses of epi-oxa-noribogaine do not induce a significant CPP or CPA in male mice (P = 0.122). The place conditioning varied by dose in females (P=0.0238), but post hoc analysis shows that neither dose produces a conditioned preference or aversion (10 mg/kg: P=0.6484; 40 mg/kg: P=0.3315), showing a trend towards CPP. Significant place preferences were observed in male and female mice for cocaine (10 mg/kg: P = 0.0029 and 0.0013) and morphine (20 mg/kg: P = 0.001 and 0.0153). **k**, Pharmacokinetic distribution of oxa-noribogaine in mice plasma and brain tissue reveals high brain penetration (10 mg/kg, s.c.). The maximal brain concentration is reached ~30 min after injection. % MPE (percentage of maximum potential effect). Data are presented as mean ± SEM, number of replicates, specific statistical tests, information on reproducibility, and P values are reported in Methods and in Supplementary Statistics Table, \**P* < 0.05, \*\**P* < 0.01, \*\*\**P* < 0.001, \*\*\*\**P* < 0.0001.

It is well established that KOR agonists induce dose-dependent hallucinosis in humans, accompanied by sedation and mood worsening in healthy subjects.^28^ In rodents and other species, kappa psychedelics also induce a sedation-like phenotype characterized by reduced locomotor activity without a complete loss of ambulatory functions (animals respond to gentle physical stimulation, but appear sedated when left undisturbed). At nearly maximal analgesic doses (e.g., ED_80_, 5.4 mg/kg), oxa-noribogaine did not induce sedation in mice in either males or females, indicated by locomotor activity comparable to that of the vehicle group in the open field (OF) test (Fig. 2f, g, h). In stark contrast, epi-oxa induced a strong sedative effect at an equianalgesic dose, an effect completely reversed by the KOR antagonist aticaprant (Fig. 2h). Thus, these two diastereomers, while producing analgesia with similar potency, show differences in sedative-like effects in mice. However, at supra-analgesic doses of oxa-noribogaine (> 10 mg/kg, s.c.), sedation in mice is observed (Extended Data Fig. 5b,d). Thus oxa-noribogaine provides a differential dosing window between the analgesic and sedation effects, in contrast to epi-oxa and typical kappa psychedelics, where the dose-dependence of sedation approximately follows that of analgesia (Extended Data Fig. 5c).

We next assessed the rewarding or aversive effects of oxa-noribogaine using the place conditioning assay. Typical kappa psychedelics show aversive effects, or conditioned place aversion (CPA), while the majority of drugs of abuse produce conditioned place preferences (CPP) in this test.^29^ Oxa-noribogaine showed no CPA or CPP at supra-analgesic doses in male mice, indicating no aversion or reward compared to significant conditioned place preferences for both cocaine (P=0.0029) and morphine (P=0.001, Fig. 2j). As sex differences have been reported for KOR modulators in reward-related behaviors in rodents,^27^ we tested also female mice to show no aversive effects induced by oxa-noribogaine or its epimer (Fig. 2j). Thus, oxa-noribogaine does not function as a rewarding or aversive stimulus in either female or male mice. Typical kappa psychedelics also induce worsening of mood in healthy humans and pro-depressive-like effects in rodents in the forced swim test.^28,30^ Oxa-noribogaine demonstrated no acute depressive-like effects in male mice at highly analgesic doses (Fig. 2i). These results indicate that even at high functional engagement of KOR, pro-depressive or aversive behavioral effects are not induced by this compound.

A pharmacokinetic profile of systemically administered oxa-noribogaine was investigated in mice (Fig. 2k) and rats (Extended Data Fig. 6). Oxa-noribogaine is highly brain penetrant and the estimated free drug concentrations in the brain after analgesic doses match well with the *in vivo* physiological readouts and *in vitro* pharmacological parameters (Extended Data Fig. 6 and Supplementary Information).

In summary, the *in vitro* and *in vivo* pharmacology indicate that, oxa-noribogaine is an atypical KOR agonist that exerts potent analgesia in the absence of common side effects of kappa psychedelics, namely, acute aversion and pro-depressive behaviors. Oxa-noribogaine also provides a pharmacological window for induction of efficacious antinociceptive effects with no or limited sedation.

## Oxa-iboga compounds do not show pro-arrhythmia risks in adult primary human heart cells

The use of ibogaine has been associated with severe cardiac side effects, most notably, cardiac arrhythmias and sudden death in humans. It has been suggested that these adverse effects are linked to the inhibition of human ether-a-go-go-related gene (hERG) potassium channels by both ibogaine and noribogaine.^6^ hERG inhibition can result in retardation of cardiomyocyte action potential repolarization and prolongation of the QT interval in the electrocardiogram, thus increasing the risk of arrhythmias. As most of the reported adverse effects occurred > 24 hours post ibogaine ingestion, noribogaine with its long circulation and large exposure appears to be the culprit of cardiac risks.^6^ This hypothesis is supported by a recent dose escalation clinical study, which reported a linear relationship between the cardiac QT interval prolongation and the plasma noribogaine concentration.^31^

Preclinical assessment of cardiotoxicity of novel compounds is complicated by species differences in cardiac ion channel expression and pharmacology. Further, inhibition of the hERG channel alone is not sufficient to predict delayed ventricular repolarization and cardiac pro-arrhythmia risk, as modulation of other ion channels involved in different phases of the cardiac action potential may mitigate or exacerbate the QT prolongation/pro-arrhythmia risk. This is particularly relevant for compounds like iboga alkaloids with complex pharmacology and multi-ion-channel activities.^32^ For example, ibogaine shows no effect on repolarization in guinea pig cardiomyocytes, likely due to the compensatory effects induced by the enhanced inhibition of L-type calcium channels in this species versus human.^33^ As a result, preclinical *in vivo* tests in rodents or other non-human species may be misleading.

We therefore used adult human primary cardiomyocytes in a state-of-the-art assay with high predictive validity of clinical cardiac effects.^32^ Adult human primary ventricular myocytes are isolated from ethically consented donor hearts, and field stimulated to induce stable contractility transients (Fig. 3). The assay detects pro-arrhythmic events such as after-contractions and contraction failures, and has been validated with clinically characterized drugs including pro-arrhythmic drugs and non-pro-arrhythmic drugs. The assay has high sensitivity and specificity, and outperforms many other assays due to the phenotypic stability of adult primary cardiomyocytes compared to human stem cell-derived cardiomyocytes.^32,34^

**Fig. 3.**
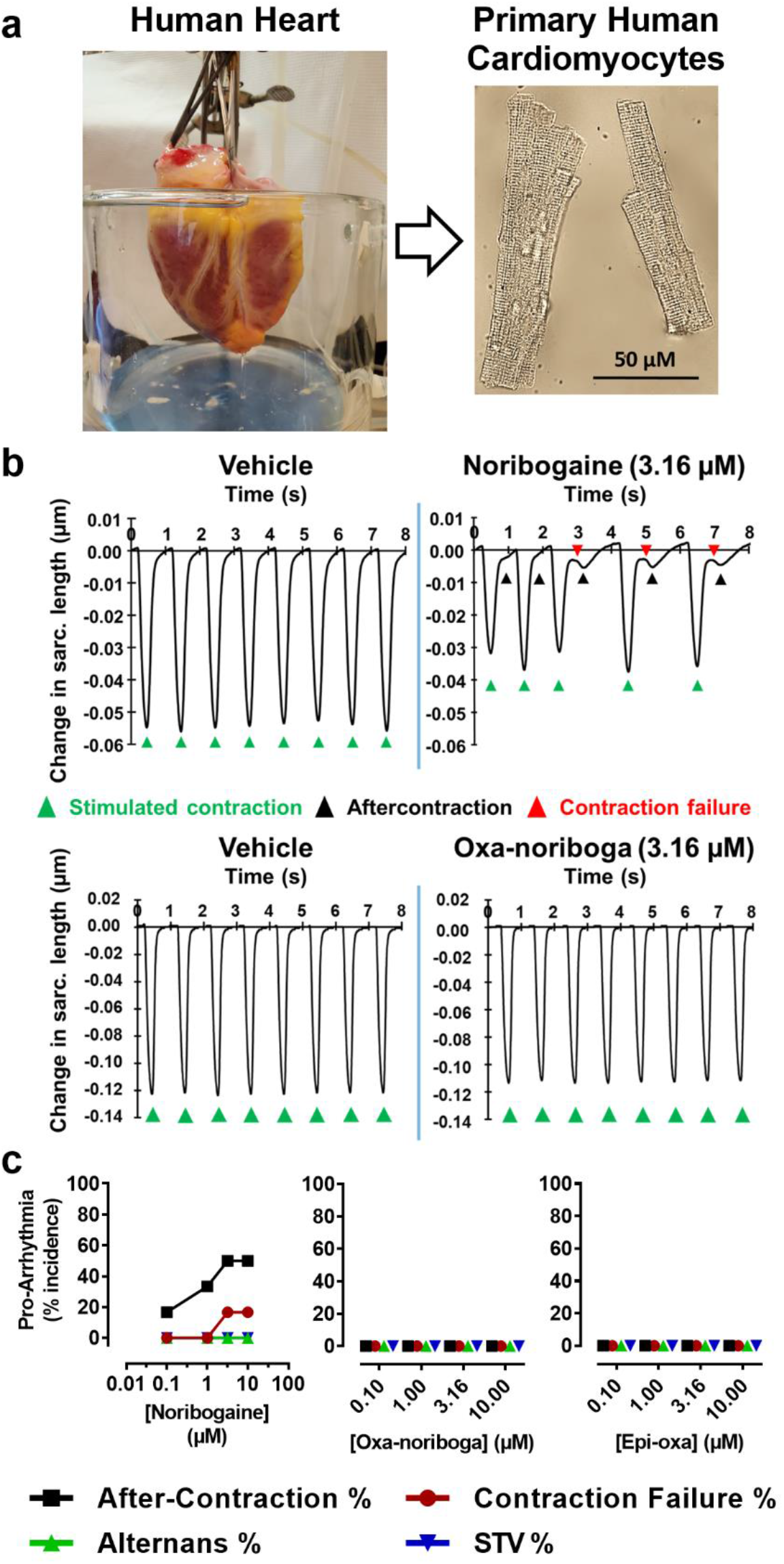
Oxa-iboga analogs do not show pro-arrhythmia risk in adult human primary cardiomyocytes. **a**, Human heart is digested to isolate single cardiomyocytes, which are field stimulated to produce a regular pattern of contractility transients. These cells are phenotypically stable and provide a robust preclinical assay with high translational validity. **b**, Representative traces capturing contraction-induced change in sarcomere length after administration of vehicle, noribogaine and oxa-noribogaine solutions. **c**, Noribogaine demonstrates pro-arrhythmic potential by causing after-contractions and contraction failures in cardiomyocytes as quantified in the plot, providing validation of this assay for the iboga compounds. No pro-arrhythmic potential was detected for oxa- or epi-oxa-noribogaine (for information on replicates see Extended Data Fig. 7). Alternans % (percentage of repetitive alternating short and long contractility amplitude transients), STV % (Short-Term Variability in percentage).

Based on available noribogaine clinical data, we hypothesized that plasma concentrations above 300-400 nM would be associated with a considerable pro-arrhythmia risk in this assay.^31^ As such, we tested it in the clinically relevant concentration range (0.1-10 μM, for more information see Extended Data Fig. 6, 7 and the Methods section). As expected, noribogaine shows a concentration-dependent pro-arrhythmia risk in the human cardiomyocyte assay, eliciting an increased frequency of aftercontractions and contraction failures (Fig. 3b, c). Greater than 20% incidence of any of the arrhythmia-associated events indicates a considerable pro-arrhythmia risk, which was observed at noribogaine concentrations of 1 μM or higher. In contrast, oxa-noribogaine and epi-oxa-noribogaine showed no pro-arrhythmic potential at any of the concentrations tested (up to 10 μM). We hypothesize that the observed differences in pro-arrhythmia risks between the oxa-iboga and iboga compounds are related to the activity at multiple ion channels of these compounds. However, to elucidate the precise mechanisms will require detailed follow-up studies, which are beyond the scope of this report. We note that only male human hearts were used in this study. As gender differences in KOR-mediated physiological processes have been reported, future studies will also need to address this important point.^27^

## Oxa-noribogaine suppresses morphine and fentanyl self-administration and cue-induced reinstatement of morphine and fentanyl responding

To assess the therapeutic potential of oxa-noribogaine we adopted a widely accepted model of opioid use, the rat intravenous self-administration (SA) paradigm.^35^ In this behavioral assay, animals are trained to respond on an operant schedule to receive intravenous infusions of a drug while discrete stimuli (tone and light) are paired with the drug delivery (Fig. 4a). For the purpose of direct comparison to the iboga alkaloids, we chose noribogaine as the standard on the basis of these points: 1) it is the dominant and long circulating molecular species after ibogaine’s administration in both humans and rats,^17,24^ 2) noribogaine shows nearly identical effects to ibogaine in the rodent opioid self-administration paradigm,^14^ and 3) noribogaine is structurally a close analog of oxa-noribogaine, differing by a single structural change. The specific experimental design (Fig. 4, Methods section) was selected to replicate previous results^14^ and validate noribogaine as the comparison drug in our experiments. The selected dose of noribogaine (40 mg/kg, i.p.) has previously been shown to fully suppress morphine SA acutely,^14^ and corresponds to the lower end of ibogaine “psychedelic reset doses” or “flood doses” used in ibogaine clinics (estimate based on simple allometric scaling).

**Fig. 4.**
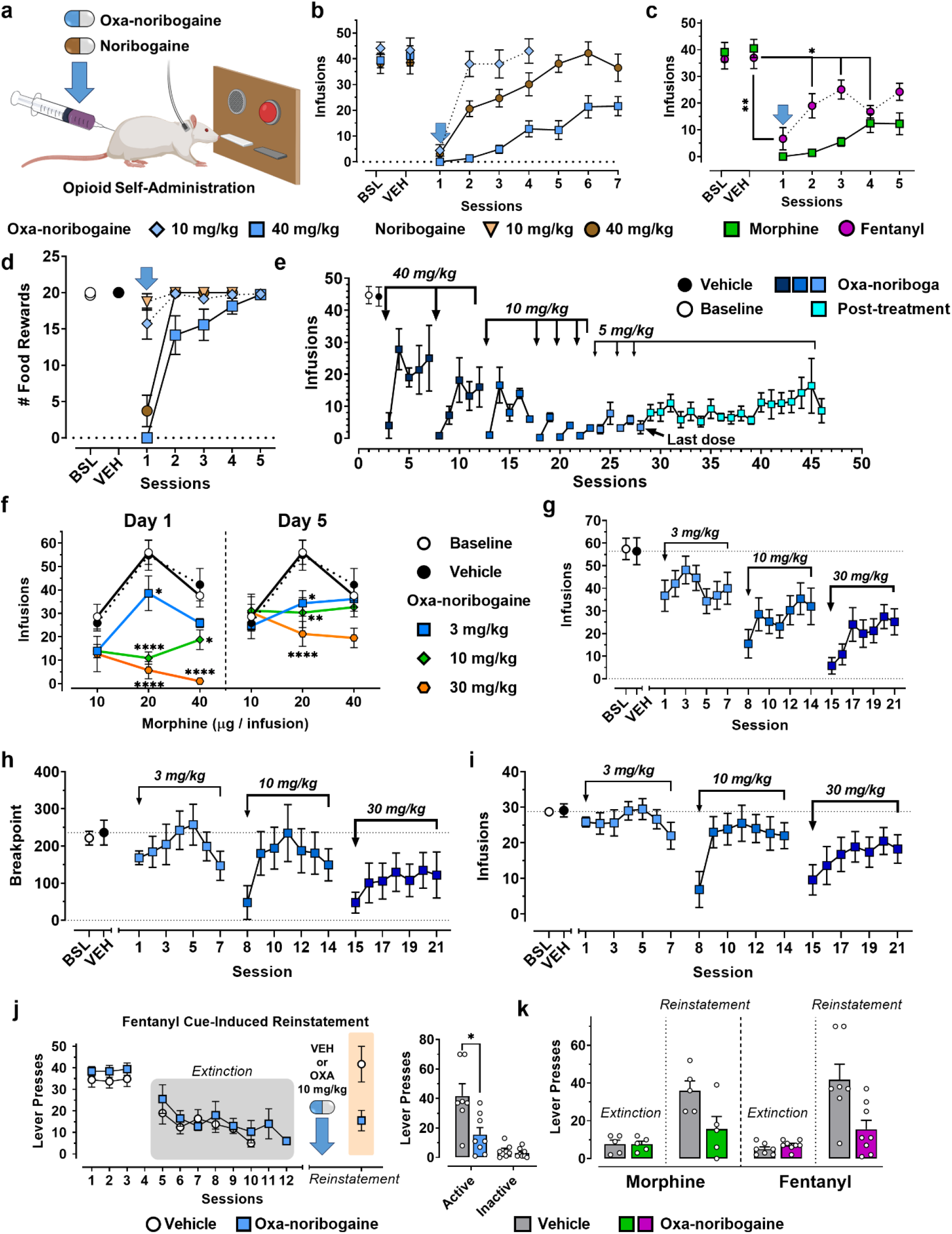
Oxa-noribogaine induces acute and long-lasting suppression of morphine and fentanyl self-administration, and relapse to morphine and fentanyl responding. **a**, A schematic depiction of the experimental treatment paradigm of opioid use disorder (OUD) animal model (male rats used in this study). **b**, Oxa-noribogaine (40 mg/kg) is more efficacious than noribogaine (40 mg/kg) in suppressing morphine self-administration (P = 0.0001, seven sessions). One injection of oxa-noribogaine (40 mg/kg) results in statistically significant suppression of morphine intake for 7 days, with suppression trends observable for at least 2 weeks (see Extended Data Fig. 8a). The effect on morphine self-administration was dose dependent. Both doses induce strong acute suppression, 40 mg/kg of oxa-noribogaine is more efficacious than 10 mg/kg in suppressing morphine self-administration (P < 0.0001, four sessions) with significant differences observed on sessions 2-4. **c**, One injection of oxa-noribogaine (40 mg/kg) results in statistically significant suppression of fentanyl intake for 4 days, with suppression trends observable for at least 5 days. Comparison with morphine data (panel 4b). **d**, Food operant intake (natural reward) is reduced following administration of 40 mg/kg but not 10 mg/kg (P < 0.0001), with significant reductions observed after 40 mg/kg on sessions 1 (P < 0.0001). The moderate dose (10 mg/kg) has a marginal effect on food intake indicating drug-selective suppression of responding at this dose. **e**, The repeated dosing regimen significantly reduced morphine SA across sessions (P = 0.0054). Arrows indicate sessions where oxa-noribogaine was administered. Repeated dosing increases oxa-noribogaine’s efficacy and leads to persistent morphine intake suppression (experiment terminated 18 days after the last dose; for individual responses see Extended Data Fig. 8). **f**, Visualization of acute (Day 1) and post-acute (Day 5) dose-dependent effects of oxa-noribogaine (3, 10 and 30 mg/kg; i.p.) on morphine dose-effect function in morphine self-administration (10, 20 and 40 μg/inf). On Day 1, oxa-noribogaine dose dependently decreases self-administration of morphine 20 μg/inf (oxa-noribogaine doses, 3 mg/kg: *P* < 0.05, 10 mg/kg: *P* < 0.0001, and 30 mg/kg: *P* < 0.0001) and 40 μg/inf (10 mg/kg: *P* < 0.0001, and 30 mg/kg: *P* < 0.0001), and intake continued to be significantly decreased on Day 5 for 20 μg/inf (Oxa-noribogaine doses, 3 mg/kg: *P* < 0.05, 10 mg/kg: *P* = 0.0044, and 30 mg/kg: *P* < 0.0001). **g**, A temporal profile across all sessions of the dose dependent effects of oxa-noribogaine in the rat cohort self-administering 20 μg/inf of morphine. **h**, Oxa-noribogaine (3, 10 and 30 mg/kg) decreases the reinforcing efficacy of morphine using progressive ratio (PR) schedule of reinforcement. Temporal profile of breakpoints and **i**, infusions obtained. **j**, A single dose of oxa-noribogaine (10 mg/kg) reduces cue-induced fentanyl seeking (P = 0.0141). **k**, Bar graph visualization of morphine (P = 0.042) and fentanyl cue-induced reinstatement following extinction period (extinction bars represent average responding of subjects in their respective last extinction session, when responding decreased to < 20% of baseline). Data are presented as mean ± SEM, specific statistical tests, information on reproducibility, and P values are reported in Methods and in Supplementary Statistics Table, \**P* < 0.05, \*\**P* < 0.01, \*\*\**P* < 0.001, \*\*\*\**P* < 0.0001.

We found that noribogaine suppressed morphine self-administration in the session immediately following its administration (day 1) and led to a partial but statistically significant suppression of morphine intake on days 2 and 3, returning to pre-treatment level of morphine intake on days 4 and 5 (Fig. 4b, male rats). Using a non-drug reinforcer, we found responding maintained by food was also suppressed on day 1 (by > 80%) but returned to baseline responding on Day 2 (Fig. 4d). These results replicate the previous studies in terms of the efficacy (extent of morphine intake suppression), selectivity (drug vs food), and duration of the effect.^14^ We next examined oxa-noribogaine under the same conditions, which induced a more profound and longer-lasting suppression of morphine intake (Fig. 4b, P = 0.0001), an effect that was statistically significant for 7 days and showed a clear suppression for at least 15 days (Extended Data Fig. 8a). One subject from this cohort showed a > 80% decrease in morphine intake at the end of the second week (Days 12-15, Extended data Fig. 8b). Following oxa-noribogaine administration, food-maintained responding was suppressed on day 1, but largely returned on day 2, and did not differentiate statistically from vehicle from day 3 onward (Fig. 4d). Thus, the suppression effect of a reset dose of oxa-noribogaine is morphine-specific from day 3 onward. We next examined a 10 mg/kg dose of oxa-noribogaine (fully analgesic dose in mice, see above), which induced a strong acute suppression of morphine intake (> 85% reduction), an effect that is morphine selective as there was only a small, non-significant effect on acute food responding (Fig. 4b, d). Morphine self-administration largely returned to baseline on subsequent days. Thus, a single 10 mg/kg dose enabled selective acute suppression of morphine self-administration without affecting behavior motivated by natural rewards.

The remarkable long-term effects induced by a single administration of oxa-noribogaine prompted examination of the effect of repeated dosing, the possibility of dose tapering, and long-term post-treatment effects. The experimental design was in part guided by the clinical experience with ibogaine. According to anecdotal reports, for most opioid dependent subjects attainment of lasting abstinence requires repeated ibogaine reset sessions and/or frequent administration of maintenance doses of ibogaine.^36^ We therefore scheduled 2 reset doses (2 × 40 mg/kg) separated by 4 days (guided by the effect of a single reset dose, Fig. 4b), followed by a maintenance dose (10 mg/kg), and a series of intermittent day-on/day-off administrations of 3 × 10 and 3 × 5 mg/kg doses, to explore the effect of more frequent administration of lower doses on both acute and day-after efficacy while tapering the dose (Fig. 4e). Overall, this treatment regimen (9 oxa-noribogaine administrations over 26 days) significantly reduced morphine SA across all sessions (P = 0.0054) and led to a progressive decrease of morphine intake despite the dose tapering. Notably, repeated, intermittent 10 mg/kg doses resulted in complete suppression of morphine intake on days with oxa-noribogaine and increasing efficacy on days without treatment (Fig. 4e). Tapering to intermittent 5 mg/kg doses led to maintenance of low, residual amounts of morphine intake, and remarkably, sustained suppression was observed for an extended period of time following the last administration of oxa-noribogaine (< 30% of pretreatment morphine intake over 2.5 weeks). Examining the responses of individual subjects, some showed high and lasting response to a single reset dose of oxa-noribogaine (as seen above), while others required multiple doses to suppress morphine intake with lasting effects (Extended Data Fig. 8f). Importantly, the long-lasting effects are morphine selective, as no such suppression effects were observed in food responding (Extended Data Fig. 8g).

Illicit fentanyl use has become a major issue exacerbating the opioid epidemic and driving the alarming numbers of drug overdose deaths.^37^ Fentanyl’s high *in vivo* potency, brain penetration, intrinsic efficacy at MOR, and rapid onset of effects likely contribute to its abuse potential and overdose rates. Although rigorous clinical comparative data is not available, for the relative reinforcing efficacy of fentanyl compared to other opioids, anecdotal reports suggest a treatment-resistant phenotype in fentanyl-dependent users.^38^ We therefore examined oxa-noribogaine in fentanyl SA under the SA paradigm used for morphine. Remarkably, a single reset dose of oxa-noribogaine reduced the intake of fentanyl for at least five days (Fig. 4c), indicating the efficacy of this compound against fentanyl and thus, potentially other opioid drugs.

We next turned to examine in more detail the potential behavioral mechanisms underlying the opioid intake suppression by oxa-noribogaine. To this end, we evaluated the dose-dependent effects of oxa-noribogaine on a portion of the morphine dose-effect curve that encompassed the ascending and descending portions of the function (peak of infusions at 20 μg/infusion, Fig. 4f). Specifically, we examined three oxa-noribogaine doses (in approximate half-log steps, 3, 10, and 30 mg/kg) across all morphine infusion doses, to find that oxa-noribogaine decreased the intake of morphine at all doses, essentially flattening the morphine curve (Day 1, Fig. 4f, left panel). In addition to the acute effects, we also examined the effect of each oxa-noribogaine doses six days following administration. At the morphine dose maintaining peak intake, we observed a clear, dose-dependent morphine SA attenuation lasting over at least seven daily operant sessions after each oxa-noribogaine administration (Fig. 4g). Once again, we saw a lasting suppression of morphine intake that results in effective flattening of the morphine dose-response function, even days after oxa-noribogaine’s administration (e.g. Day 5, Fig. 4f). These results indicate that oxa-noribogaine reduces morphine responding both acutely (indirect-antagonist-like effect, “drug-on” effect) and long after oxa-noribogaine is cleared from the subjects (“drug-off” effect).

Next, we directly assessed the effect of oxa-noribogaine on the reinforcing efficacy of morphine using a progressive ratio (PR) schedule of reinforcement. Oxa-noribogaine decreased the breakpoint ratio with a clear acute effect at all three doses; in addition, the top dose induced a lasting depression of the breakpoint (Fig 4h,i). These results demonstrate that oxa-noribogaine administration reduces morphine’s reinforcing efficacy acutely and long term.

Finally, in terms of behavioral studies, we tested oxa-noribogaine’s effect in a model of a critical aspect of substance use disorders (including OUD) - relapse. Cue-induced reinstatement of responding is a well-established rat model of relapse, consisting of a period of self-administration followed by extinction of responding in the absence of drug and associated stimuli. For the test session, responding on the previously drug associated lever is recorded in the presence of stimuli previously associated with drug delivery. For fentanyl, robust baseline responding was extinguished over 8 days, followed by cue-induced reinstatement where responses were comparable to the pre-extinction level in the vehicle cohort (Fig. 4j). In contrast, oxa-noribogaine reduced reinstatement responding by approximately 60%, whereas there was no effect on inactive levers (10 mg/kg, Fig. 4j). Similarly, in another cohort, morphine reinstatement was also strongly reduced by oxa-noribogaine, at a dose that has minimal effect on food responding (10 mg/kg, Fig. 4k).

We hypothesized that the mechanisms underlying the long-term effects of oxa-noribogaine involve induction of restorative molecular programs mediated by neurotrophic factors, most notably, glial cell line derived neurotrophic factor (GDNF) and brain-derived neurotrophic factor (BDNF). Although these neurotrophic factors play complex roles in the neurobiology of substance use disorders,^39^ targeted elevation of GDNF protein levels in specific brain areas enables attenuation or reversal of various aspects of addiction-related effects (e.g. drug intake, reward expression, and dopamine neuron firing patterns). For example, GDNF injection in the ventral tegmental area (VTA) has been shown to reverse molecular, electrophysiological, and behavioral markers of addiction-like phenotypes induced by chronic morphine, cocaine, and alcohol exposure in rats.^40^ VTA dopaminergic neurons project to the mPFC, where deep layer pyramidal cells project back to the VTA, which in turn provides dense dopaminergic innervation in the nucleus accumbens (NAc, Fig. 5a); all three brain areas and the corresponding circuits play prominent roles in addiction biology. We focused directly on alternations of neurotrophin protein levels in these brain regions as 1) GDNF gene expression loci do not always match those of GDNF receptors, and 2) GDNF can be transported over long distances. Specifically, we investigated long-term changes in protein levels of GDNF in the VTA, NAc, and mPFC up to five days after a single reset dose of oxa-noribogaine (Fig. 5b). Remarkably, we observed 200% increase in GDNF levels in the VTA and 100% in the mPFC on day 5 after treatment, while there was no change in the NAc. Noribogaine as a comparator showed no significant change in any of the tested brain regions. We also examined mature BDNF protein levels in the mPFC, where an increase was observed at both one and five days after the oxa-noribogaine treatment (day 1 is statistically significant, day 5 shows a trend, Fig. 5c). Noribogaine also showed elevated BDNF at both time points. These measurements support our hypothesis and suggest targeted induction of neuroplasticity and restoration programs mediated by GDNF and BDNF. To test the association of GDNF levels in the VTA and mPFC with KOR, the key molecular target of oxa-noribogaine, we treated rats with aticaprant, a selective KOR antagonist, prior to oxa-noribogaine administration. Under these conditions, the induction of GDNF was blocked in both mPFC and VTA, revealing protein levels comparable to those of control/vehicle rats (Fig. 5d). These results suggest that KOR activation plays a role in the long-term induction of GDNF by oxa-noribogaine in addiction-relevant brain circuits.

**Fig. 5.**
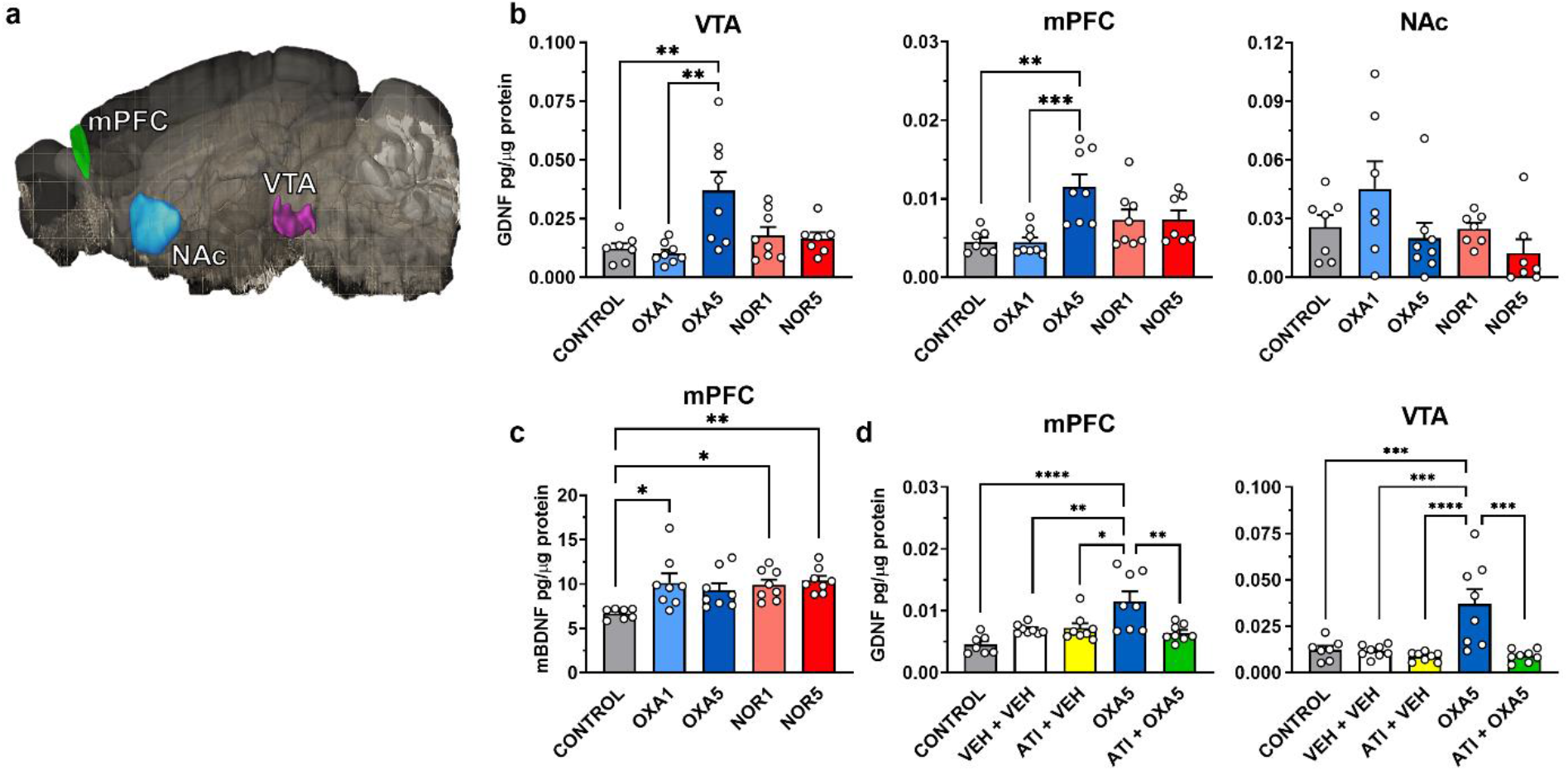
Oxa-noribogaine in rats induces long-term elevation of GDNF and BDNF in critical brain circuits mediated by KOR. **a**, Representative illustration of a left hemisphere (sagittal slice) of a rodent brain (mouse, Allen Institute)^41^, the location and shape of medial prefrontal cortex (mPFC), nucleus accumbens (NAc) and ventral tegmental area (VTA) are highlighted. **b**, Administration of oxa-noribogaine (40 mg/kg; i.p.) but not noribogaine (40 mg/kg; i.p.) significantly increased GDNF protein levels in the VTA and mPFC after 5 days (OXA5). No significant changes in GDNF expression were observed in NAc after administration of either noribogaine or oxa-noribogaine. **c**, Increased levels of mature BDNF protein in the mPFC after a single dose of either oxa-noribogaine (40 mg/kg; i.p.) or noribogaine (40 mg/kg; i.p.) were detected after 24 h (OXA1 and NOR1) and remained elevated for up to 5 days (OXA5 and NOR5). **d**, Pre-treatment of rats with KOR selective antagonist aticaprant (ATI) before oxa-noribogaine administration prevents the increase in GDNF expression. Data are presented as mean ± SEM, specific statistical tests, information on reproducibility, and P values are reported in Methods and in Supplementary Statistics Table, *\*P* < 0.05, \*\**P* < 0.01, \*\*\**P* < 0.001, \*\*\*\**P* < 0.0001.

## Discussion

Currently approved medications for treatment of SUDs do not reverse or weaken the persistent negative alterations in neurocircuitry that underlie addiction-related states in animals, and presumably in humans.^3,42^ Although considerable progress has been made in the development of animal models of various aspects of addiction phenotypes (e.g. drug intake and cue-induced reinstatement of drug seeking),^4,43,44^ the SUD preclinical literature has largely focused on the acute effects of pharmacological interventions, where typically the desirable effect wanes along with the drug’s clearance. In contrast, ibogaine serves as inspiration for a different kind of pharmacotherapeutic capable of inducing profound and lasting interruption of drug intake and relapse in human and animal subjects. The central hypothesis for this new direction in drug discovery is that these lasting behavioral effects are driven by desirable neuroplasticity and neuro-restorative alterations of relevant circuitry induced by drugs like ibogaine. In support of this hypothesis, it has been reported that ibogaine induces expression of GDNF in rats, which in turn resets the function of dopaminergic reward circuitry.^45^ We have recently replicated this result and shown that ibogaine also induces gene expression of BDNF in the prefrontal cortex.^46^ It thus appears that multiple neurotrophic systems are activated by ibogaine that may drive synaptic and circuit restoration, which in turn underlies the observed interruption of addiction phenotypes. This hypothesis is further supported by the observations in this study, where oxa-noribogaine induced marked elevation of GDNF protein levels in the VTA and mPFC, as well as BDNF protein levels in the mPFC, five days after a single administration. Previous reports largely focused on gene expression profiling within 24 h; in contrast, our results indicate neurotrophin protein elevation up to 5 days post a single-dose pharmacological intervention.

We note that neurotrophic factors play a complex role in the neurobiology of addiction. Neurotrophins likely mediate neuroadaptations to chronic drug use and development of certain dimensions of substance use disorder phenotypes (e.g., drug sensitization and incubation of craving). Contradictory effects on drug use have been reported depending on the reinforcing drug and experimental design of neurotrophin manipulations.^39,47^ However, targeted induction or injection of certain neurotrophins, GDNF in particular, has the ability to reverse many aspects of already established addiction-like phenotypes. For example, injection of GDNF in the VTA reverses persistent molecular (e.g., increased protein levels of tyrosine hydroxylase) and behavioral (reward expression or drug intake) markers induced by chronic morphine administration. Further, chronic morphine intake reduces GDNF signaling in the VTA (determined by the phosphorylation status of its receptor), which is reversed by GDNF induction in the same brain region.^40^ Strong evidence has also been accumulated for GDNF in the VTA as an inhibitor of alcohol intake and other aspects of alcohol dependence in rodents.^48^ We therefore propose a neurotrophin hypothesis of addiction treatment by iboga compounds, where 1) neurotrophins mediate in part the neuroplastic (or hyperplastic) processes induced by chronic use of reinforcing drugs, leading to pathological addiction states, but 2) targeted induction of neurotrophins by compounds like ibogaine or oxa-noribogaine facilitates interruption or weakening of such states by re-opening neuroplasticity windows in specific brain circuits.

Sampling GDNF and BDNF protein levels via the quantitative ELISA assays used in our study provides a relatively straightforward experimental system for further study of this hypothesis and testing of advanced compounds with well characterized pharmacology. The linkage of GDNF protein induction to KOR agonism is intriguing and highlights the potential of atypical KOR agonists that lack adverse effects such as aversion and sedation. One limitation in our study was the use of unconditioned animals for the neurotrophin examination, not subjects trained to self-administer morphine or fentanyl. These follow-up questions will be addressed in future studies.

Considering ibogaine’s unique clinical effects, as well as its structural and pharmacological complexities, the iboga system provides a rich discovery platform.^21,25^ We here described the oxa-iboga compounds as a new class of iboga alkaloids created by a single but highly consequential structural permutation of the iboga skeleton. Oxa-noribogaine accentuates KOR agonistic activity, enhances therapeutic-like activity in models of OUD, and addresses cardiotoxicity, in comparison to noribogaine. A single reset dose of oxa-noribogaine leads to suppression of morphine SA for at least 1 week, while repeated dosing increases its efficacy, enables dose tapering, and leads to a persistent suppression of morphine intake. The lasting effects on morphine self-administration are not seen in food responding. Further, a single dose of oxa-noribogaine also reduces fentanyl intake for at least 5 days, which correlates with elevated GDNF protein levels in the VTA and mPFC, indicating a lasting therapeutic-like effect against different opioids, including fentanyl which has rapidly become the primary driver of the opioid epidemic.

We also show that oxa-noribogaine decreases the reinforcing efficacy of morphine, both acutely and in the long term, and markedly reduces cue-induced reinstatement responding for fentanyl and morphine in a relapse model.

These results support the rationale for further development of oxa-noribogaine, or a related analog, as a promising experimental candidate for OUD treatment. However, there are limitations in the present work that will need to be addressed in future studies. First, only male rats were used in the opioid self-administration and reinstatement experiments, while sex differences in KOR-mediated physiological and pharmacological effects have been reported.^27^ We do not expect major sex differences in the effect of oxa-noribogaine on opioid self-administration assays, as in mice there are small to no differences between females and males in a series of behavioral readouts in response to oxa-noribogaine; including assays used for target engagement/therapeutic-like effects (tail-flick test), adverse effects (open field), and reward-related responses (CPP/CPA). Further, the efficacy and duration of effect of noribogaine in male rats, as observed in the current study, mirrors perfectly these effects in female rats reported in previous studies.^14^ Second, we did not examine the effect of oxa-noribogaine on somatic withdrawal signs during or after the self-administration experiments. As no overt somatic symptoms were present in our studies, a more systematic behavioral mapping would be required. We expect that the analgesic effects of oxa-noribogaine will be highly beneficial in alleviating mechanical hypersensitivity (measured via the von Frey test), as a model of hyperalgesic phenotype in opioid-dependent and post-dependent human subjects.^49^ Third, we did not explore alternative intervention designs where, for example, oxa-noriboigaine would be administered during abstinence. It is plausible that induction of GDNF and other neurotrophins in this phase could facilitate a neurobiological incubation process resulting in increased drug responding and reinforcement.^36,45^ In our view this is an unlikely scenario, if the anecdotal evidence from ibogaine clinics provides any clues in this direction. Fourth, we did not conduct direct comparison of oxa-noribogaine’s effect on the opioid versus food reinforcers in a drug-food choice paradigm. Instead, this comparison was obtained in separate cohorts each trained to one reinforcing agent. From a translational viewpoint, the suppression of opioid intake over days and weeks while subjects feed and grow normally is most relevant. However, for a detailed behavioral analysis the comparison between opioid and food in a direct choice procedure will be required.^50,51^ These important aspects of preclinical characterization discussed above will be examined in future studies.

As with all preclinical SUD studies, however, there is the question of predictive validity of the animal models for candidate therapeutics, where SA paradigms have a mixed track record.^4,52^ In the present study, the starting point is a drug with significant clinical effects (although not yet confirmed in controlled studies), which was reverse translated to provide a benchmark for preclinical evaluation of new compounds, in terms of both efficacy and safety. In direct comparison to noribogaine, oxa-noribogaine is superior in terms of both maximal efficacy (extent of intake suppression) and duration of its after-effects. For another comparison, 18-methoxycoronaridine (18MC) is a synthetic iboga analog that has progressed through preclinical and Phase I clinical safety studies, but its clinical efficacy has not yet been reported. In preclinical assays in rats, 18MC showed efficacy and duration of effect on morphine intake comparable to ibogaine and noribogaine,^53^ and thus, an oxa-iboga analog likely offers a superior iboga candidate for SUD treatment. Ibogaine has also served, in part, as inspiration for the creation of 1,2,3,4,5,6-hexahydroazepino[4,5-b]indoles, for example compound U-22394A (“U” for Upjohn Company, also known as PNU-22394) that has been in clinical tests but not for SUD indications.^54^ An analog of PNU-compounds (named TBG) has recently been examined in a heroin SA test in rats and showed no lasting effect on heroin SA beyond the acute and non-selective suppression of heroin and sucrose intake.^55^ However, the PNU-compounds are pharmacologically distinct from iboga, as they are direct ligands of multiple 5HT receptors and lack activity at KOR and other known iboga targets, while iboga alkaloids have no direct interactions with 5HT receptors within a relevant concentration range. The PNU-compounds are structurally and pharmacologically related to lorcaserin (5HT2C > 5HT2A preferring agonist), which prior to its market withdrawal generated excitement in SUD research (e.g., showing suppression of heroin SA in rats^56^ and cocaine SA in monkeys^57^). Lorcaserin maintained effective suppression of cocaine SA for an extended period of time by daily dosing prior to SA sessions (14 days), however the suppression effect completely dissipated as soon as the treatment ended.^57^ This example highlights the difference between drugs that are effective acutely, and those - like oxa-iboga and iboga - that may produce lasting or even persistent alterations of SUD-related phenotypes.

The oxa-iboga compounds act as potent KOR agonists *in vitro* and *in vivo*, but exhibit atypical behavioral features compared to standard kappa psychedelics. Generally, KOR agonists are acutely pro-depressive and aversive; in drug SA studies, the effects vary depending on the experimental design.^58^ For example, mixing in KOR agonist nalfurafine to self-administered oxycodone decreased the opioid intake in rats,^59^ whereas pretreatment with nalfurafine or U50,488 resulted in an increased cocaine intake in mice.^60^ However, the long-term effects of kappa psychedelics in drug SA studies have not been systematically examined. Oxa-noribogaine, although a potent KOR agonist, does not show acute pro-depressive or aversive effects, and induces desirable long-term effects in OUD models. It is not clear what mechanistically imparts these atypical features, whether differential KOR signaling^61–64^ and/or polypharmacology.^65,66^ The present studies show a clear differentiation of oxa-iboga from standard KOR agonists via markedly decreased receptor signaling efficacy in a functional assay that approximates compounds’ intrinsic signaling efficacy. Oxa-noribogaine does not show any marked preference between Gi activation and β-arrestin2 recruitment. We therefore propose the low signaling efficacy as a working hypothesis for the unique behavioral effects of oxa-noribogaine, such as the lack of aversion or pro-depressive effects. This proposal is in line with a renewed interest in low efficacy agonists at MOR, as an approach to attenuate the notorious adverse effects of such compounds.^67–69^ The inherited iboga pharmacological activity may also contribute to the *in vivo* pharmacological profile of oxa-noribogaine.

In summary, we propose a mechanistic model where KOR activation, as the primary molecular event, largely mediates the acute behavioral effects and triggers neurotrophic and neuro-repair signaling pathways in several circuits (e.g. prefrontal control of limbic systems and reward circuitry), which underpin, at least in part, the observed long-term effects in OUD models. The oxa-iboga alkaloids are a new class of compounds that maintains and enhances the ability of iboga compounds to effect lasting alteration of addiction-related states while addressing iboga’s cardiac liability. As such, these compounds represent exciting candidates for a new kind of anti-SUD pharmacotherapeutic.

## Supporting information

Supplementary_Statistical_Table

## Acknowledgements

This work was supported by Columbia University (CU, D. Sames), The High Point University (HPU, S. Hemby), the National Institute on Drug Addiction (NIDA) of the National Institute of Health (NIH), grants R01DA050613 (D. Sames and S. Hemby), R33DA045884 (S. Majumdar), R33DA038858 (V. Katritch), and the National Institute of Mental Health (NIMH), grant R21MH116462 (J. Pintar), and the Hope for Depression Research Foundation (J. A. Javitch). This research was funded in part through the NIH/NCI Cancer Center Support Grant P30 CA008748 to MSKCC. V.H. acknowledges the Experientia Foundation Postdoctoral Fellowship. The authors would like to thank William Nguyen, Ky Truong, Lana Rasoul, Alexa Stafford, Guy Page and Dr. Richard Kondo (AnaBios Corporation) for technical assistance and helpful discussions, Dr. Fereshteh Zandkarimi (CU) for measuring the high-resolution mass spectra of novel compounds and to Prof. Gerard F. Parkin and David A. Sambade (CU) for solving the X-ray crystal structure of oxa-ibogaine. Min H. (Jimmy) Kyaw (CU) reproduced the palladium-catalyzed cyclization procedure under supervision by V.H. A.H. was supported by the Summer Research fellowship (SuRF) at HPU. We thank Dr. Ignacio Carrera (Universidad de la República, Uruguay) for helpful discussions on iboga pharmacology, and The Psychoactive Drug Screening Program (PDSP) at UNC at Chapel Hill for performing receptor panel screening. Certain graphics (Fig. 4a) were created using BioRender.com. Research reported in this publication was supported by the Office of The Director, National Institutes of Health of the National Institutes of Health under Award Number S10OD026749. The content is solely the responsibility of the authors and does not necessarily represent the official views of the National Institutes of Health.

## Author contributions

D.S. conceptualized and supervised the work. V.H. contributed to the design, performed scale-up synthesis, developed palladium-catalyzed synthetic approach, supervised pharmacological characterization including data interpretation and summarized the collected data for publication. A.C.K. contributed to the design, synthesis, and pharmacological characterization of compounds, including the design and first synthesis of oxa-noribogaine, carried out early synthetic work including reaction development, optimization, and compound characterization and supervised initial pharmacological characterization and data interpretation. B.B. contributed to the scale-up synthesis, contributed to the design and performed mice experiments conducted at Columbia University (tail-flick, open field, and forced swim tests) and interpreted the corresponding results. S.M. and L.S. carried out self-administration and conditioned place preference experiments performed at HPU. A.H. trained rats for self-administration studies and conducted the food maintained responding experiments. M.G.W and M.N. performed the BRET functional assays. A.H. performed the opioid binding assays. M.A. performed the tail-flick mice assay comparing WT and KO mice and J.E.P. supervised the tail-flick mice assays with KO mice. C.H. carried out the serotonin transporter inhibition assay. N.A.-G. provided expert consultation for cardiotoxicity studies. S.A.Z. carried out the docking studies and V.K. supervised the docking studies. M. Y. oversaw the *in vivo* experiments at Columbia University and provided expert guidance. J.A.J. supervised the BRET functional assays and interpretation of the corresponding results. S.M. supervised the binding assays, functional assays and early *in vivo* mice tests carried out at MSKCC. S.E.H. designed, guided and supervised the CPP tests in mice, neurotrophin protein brain level determination in rats, self-administration and reinstatement studies in rats, statistical analyses and interpretation of the corresponding results. D.S. wrote the manuscript with significant help from V.H. and S.E.H. All authors contributed to editing of the manuscript.

## Materials & Correspondence

Correspondence and requests for materials should be addressed to D.S.

## Reporting summary

Further information on research design is available in the Nature Research Reporting Summary linked to this paper.

## Data availability

Raw data are available from authors upon reasonable request. CCDC deposition #2215015 contains the supplementary crystallographic data (oxa-ibogaine **10a**) for this paper. These data can be obtained free of charge from The Cambridge Crystallographic Data Centre via www.ccdc.cam.ac.uk/structures.

## Code availability

Custom-written data analysis codes are available from authors upon reasonable request.

## Extended Data Figures

**Extended Data Fig. 1.**
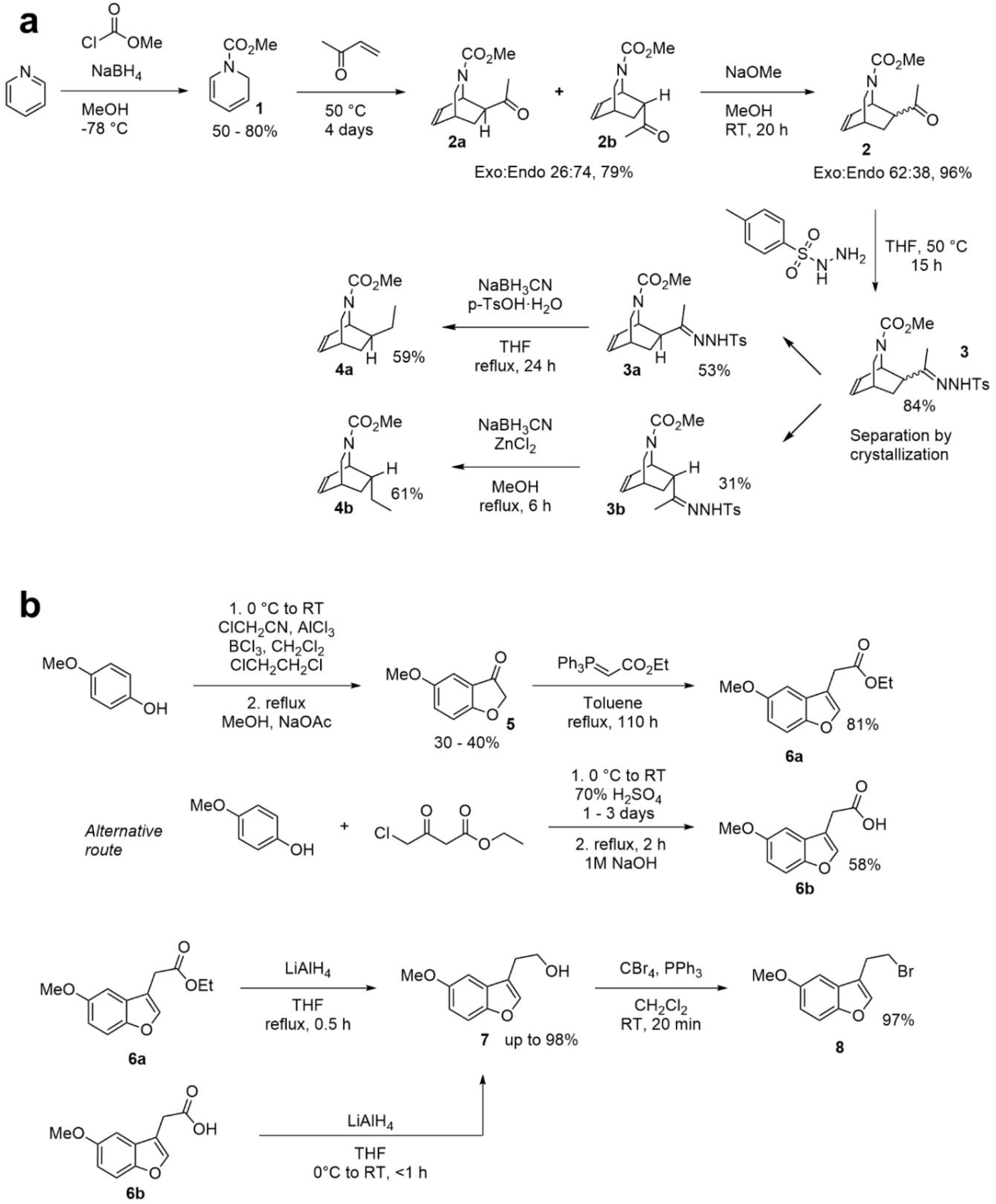
Synthesis of oxa-iboga analog intermediates. **a,** The isoquinuclidine ring was assembled via a Diels-Alder reaction starting from dihydropyridine 1. **b**, The benzofuran intermediate was prepared using two slightly different reaction pathways, with the alternative route proving to be a more convenient choice for a multigram synthesis.

**Extended Data Fig. 2.**
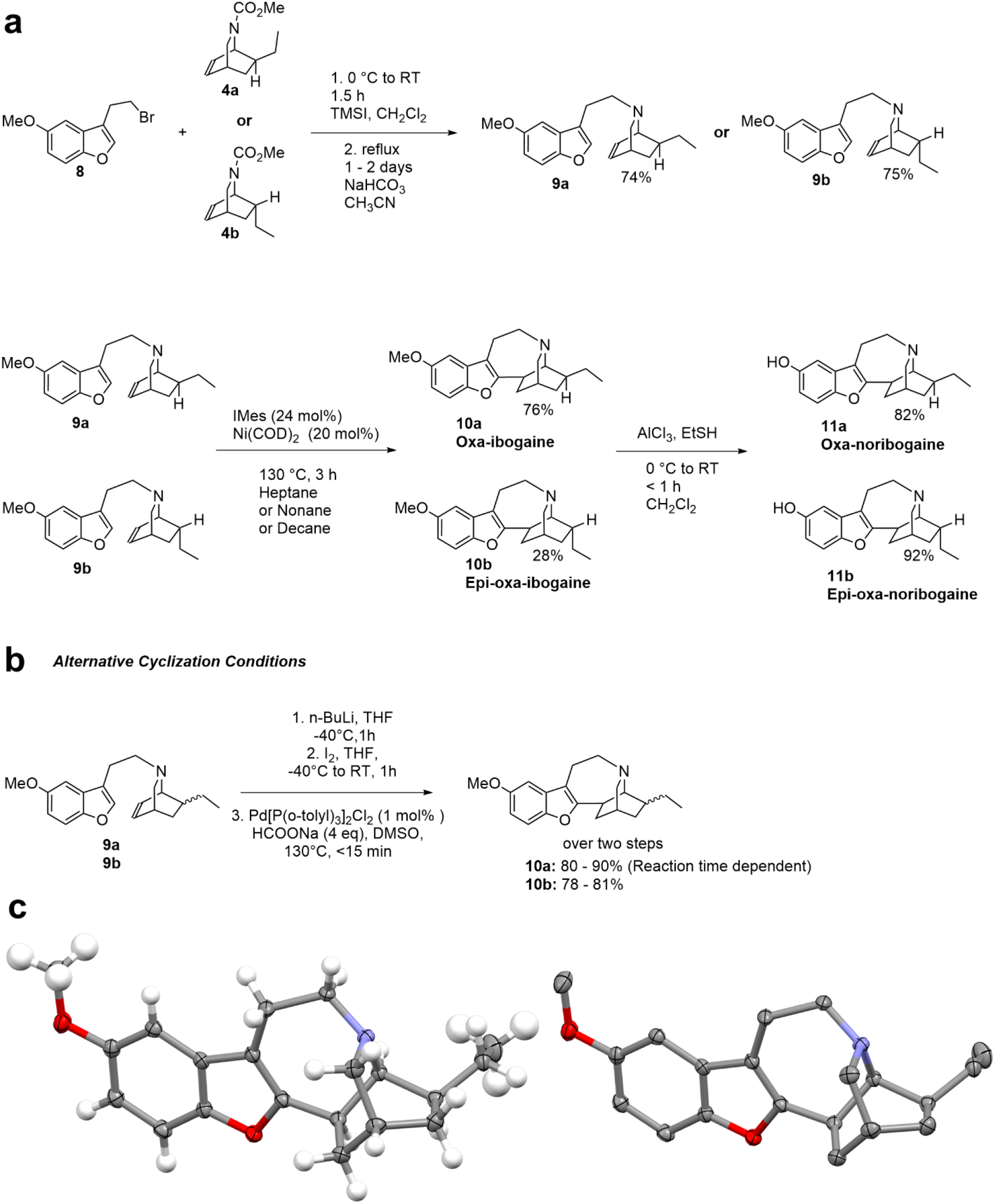
Synthesis of oxa-iboga analogs. **a**, Formation of the 7-membered azepine ring can be achieved using a nickel-catalyzed coupling between the benzofuran and isoquinuclidine moieties or **b**, via a sequence of lithiation/iodination followed by reductive Heck coupling. **c**, An X-ray crystallography structure of racemic oxa-ibogaine (one enantiomer shown) in thermal ellipsoid style (50% probability). Atom color coding: H white, C gray, O red, N blue.

**Extended Data Fig. 3.**
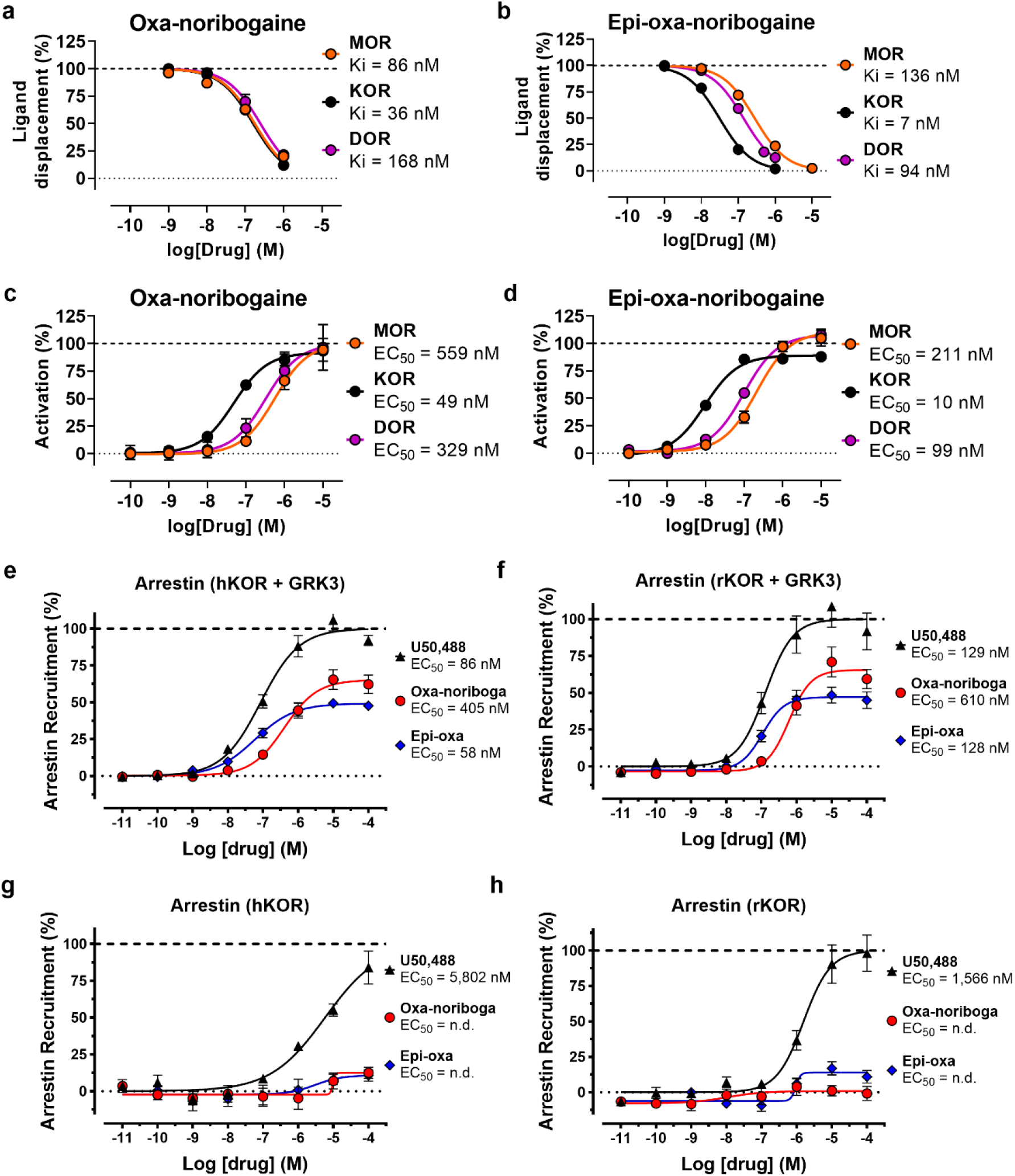
*In vitro* pharmacological characterization of noribogaine and oxa-iboga analogs. **a**, Binding affinity measured by radioligand (*[^125^I]-IBNtxA*) displacement experiments at mouse opioid receptors (mMOR, mKOR and mDOR) for oxa-noribogaine and **b**, epi-oxa-noribogaine. **c**, Agonist activity at mouse opioid receptors determined by [^35^S]GTP*γ*S assay for oxa-noribogaine and **d**, epi-oxa-noribogaine. **e**, Oxa-iboga alkaloids recruit β-arrestin with partial efficacy compared to U50,488 in HEK-293 cells expressing either human (hKOR) or **f**, rat (rKOR) kappa opioid receptors. **g**, **h**, The efficacy and potency are severely diminished if no additional G-protein-coupled receptor kinase 3 (GRK3) is added into the test system. Data are presented as mean ± SEM (n ≥ 3).

**Extended Data Fig. 4.**
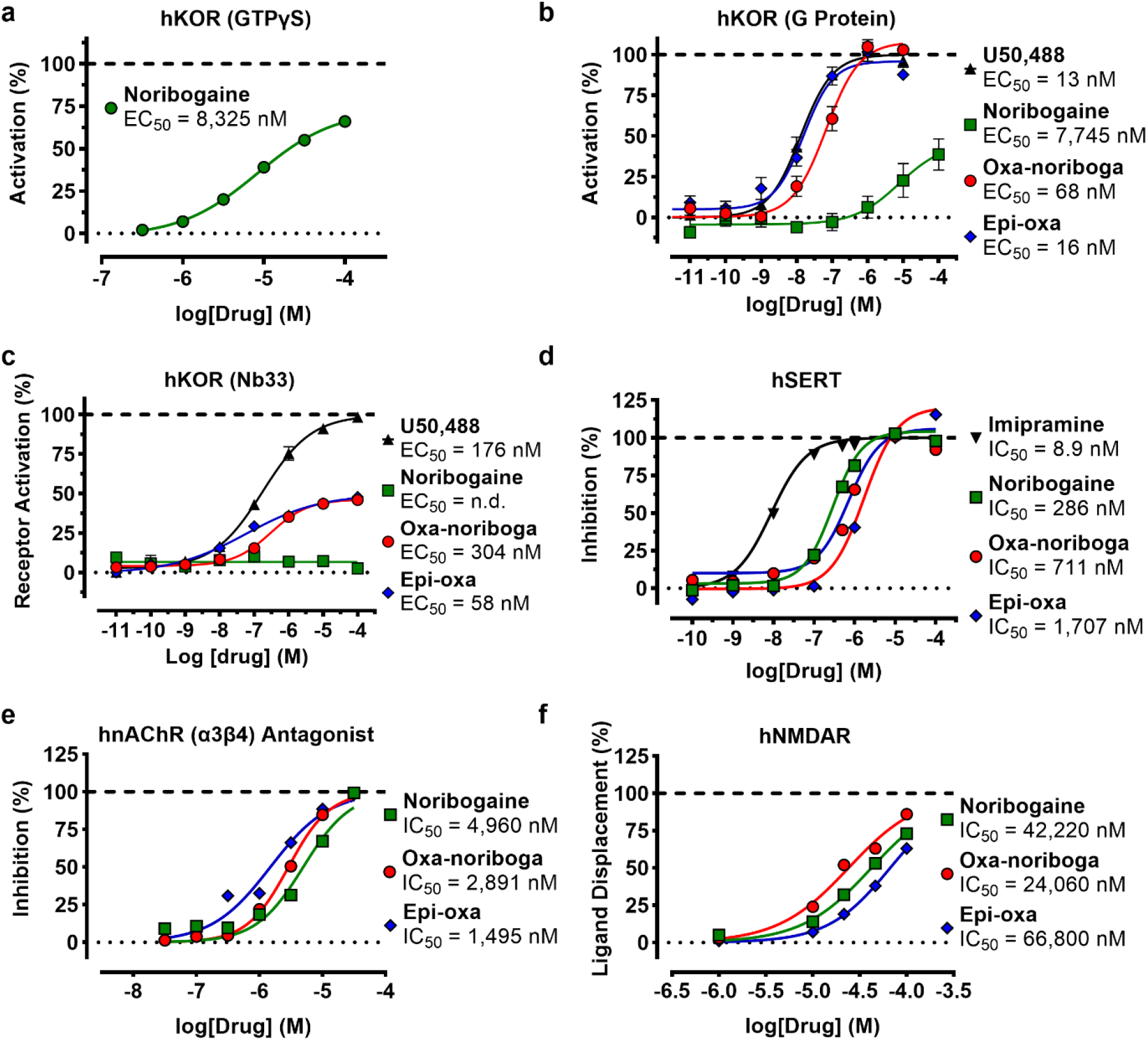
*In vitro* pharmacological characterization of noribogaine and oxa-iboga analogs continued. **a**, Agonist activity of noribogaine at human KOR determined by [^35^S]GTP*γ*S assay. **b**, Human KOR agonist activity (G protein BRET assay) for noribogaine and oxa-noribogaine analogs. **c**, Human KOR agonist activity determined using a nanobody sensor (Nb33 BRET assay) for noribogaine and oxa-noribogaine analogs. **d**, Inhibition of human serotonin transporter (hSERT) by noribogaine and oxa-iboga analogs. **e**, Inhibition of human nicotinic acetylcholine receptor (nAChR α3β4) by iboga alkaloids as determined by electrophysiological assays. **f,** Binding affinity (radioligand displacement assay) of iboga alkaloids at rat NMDA receptor. Data are presented as mean ± SEM.

**Extended Data Fig. 5.**
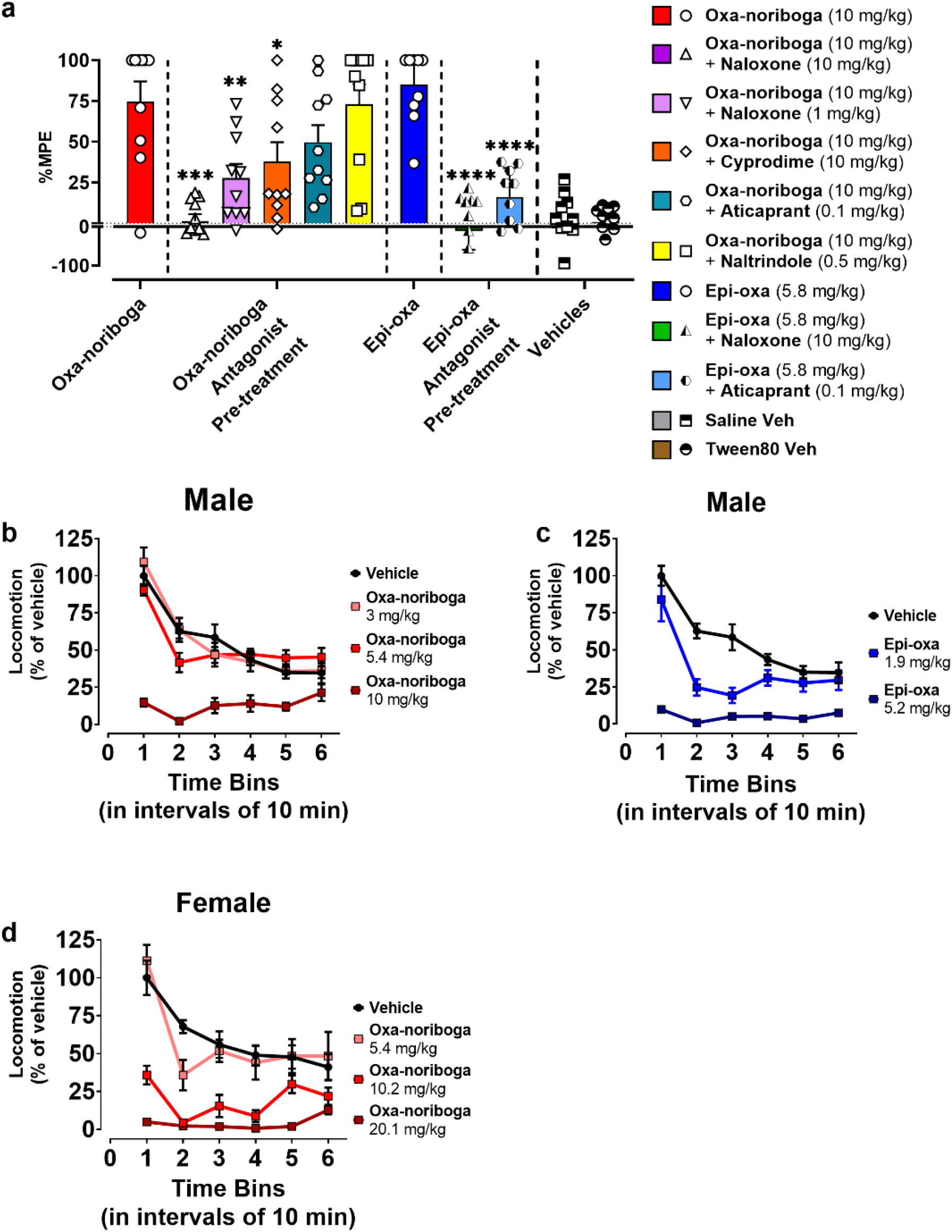
I*n vivo* characterization of oxa-noribogaine analogs by tail-flick and open field tests. **a**, Effect of pre-treatment with antagonists naltrindole (DOR selective), naloxone (non-selective at 10 mg/kg, MOR preferring at 1 mg/kg), cyprodime (MOR selective) and aticaprant (KOR selective) on oxa-noribogaine induced nociception in male mice and naloxone (10 mg/kg) and aticaprant (0.1 mg/kg) on epi-oxa-noribogaine. **b**, Escalating dose effect of oxa-iboga analogs on locomotion in OF test (male mice). Oxa-noribogaine (3.0 mg/kg = ED_50_, 5.4 mg/kg = ED_80_ and 10 mg/kg > ED_95_) and **c** epi-oxa-noribogaine (1.9 mg/kg = ED_50_ and 5.2 mg/kg = ED_80_). **d**, Escalating dose effect of oxa-noribogaine (5.4 mg/kg = male ED_80_, 10.2 mg/kg = ED_80_ and 20.1 mg/kg > ED_95_) on locomotion in OF test (female mice). Data are presented as mean ± SEM, specific statistical tests, information on reproducibility, and P values are reported in Methods and in Supplementary Statistics Table, \**P* < 0.05, \*\**P* < 0.01, \*\*\**P* < 0.001.

**Extended Data Fig. 6.**
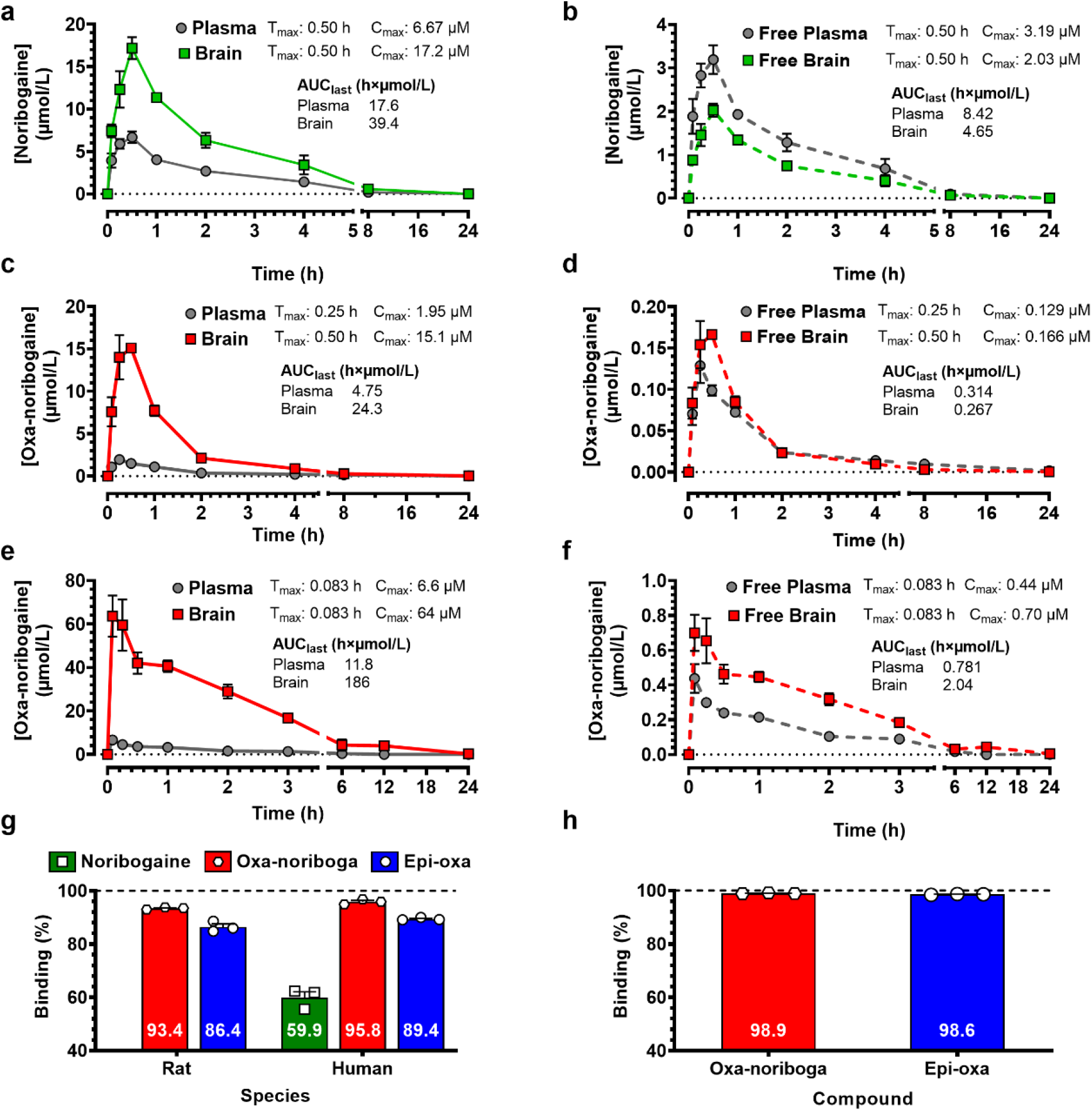
Pharmacokinetic (PK) distribution (mice, rats), and plasma protein (rats, human) and brain tissue (rat) binding of noribogaine and oxa-iboga analogs. Noribogaine (s.c. 10 mg/kg) PK in male C57BL/6 mice, **a** total and **b** estimated free concentrations. **c**, Oxa-noribogaine exhibits rapid and high brain penetration in PK (male C57BL/6 mice, s.c. 10 mg/kg, brain/plasma = 5.5). Approximately 85% of the compound is cleared in the first 2 hours and only traces remain after 8 hours. **d**, Estimated free plasma and brain drug concentrations. Assuming linearity of PK below 10 mg/kg, C_max(brain)_ at an ED_50_ analgesic dose (3 mg/kg) is expected to be ~ 49 nM, which matches very well the *in vitro* KOR activation potency (EC_50_ = 41 nM in BRET assay, EC_50_ = 49 nM, in [^35^S]GTPγS assay). Oxa-noribogaine (i.p. 40 mg/kg) PK in male Wistar rats, **e**, total and **f**, estimated free concentrations. Rat plasma protein and brain tissue binding data were used to estimate free drug fractions in mice. The Cmax and area under the curve (AUC, drug exposure) values for estimated free drugs are much smaller for oxa-noribogaine in comparison to noribogaine which is relevant for interpreting the oxa-noribogaine’s superior efficacy in addiction models. **g**, Rat and human plasma protein binding and **h** rat brain tissue binding data of noribogaine and oxa-iboga analogs. Data are presented as mean ± SEM.

**Extended Data Fig. 7.**
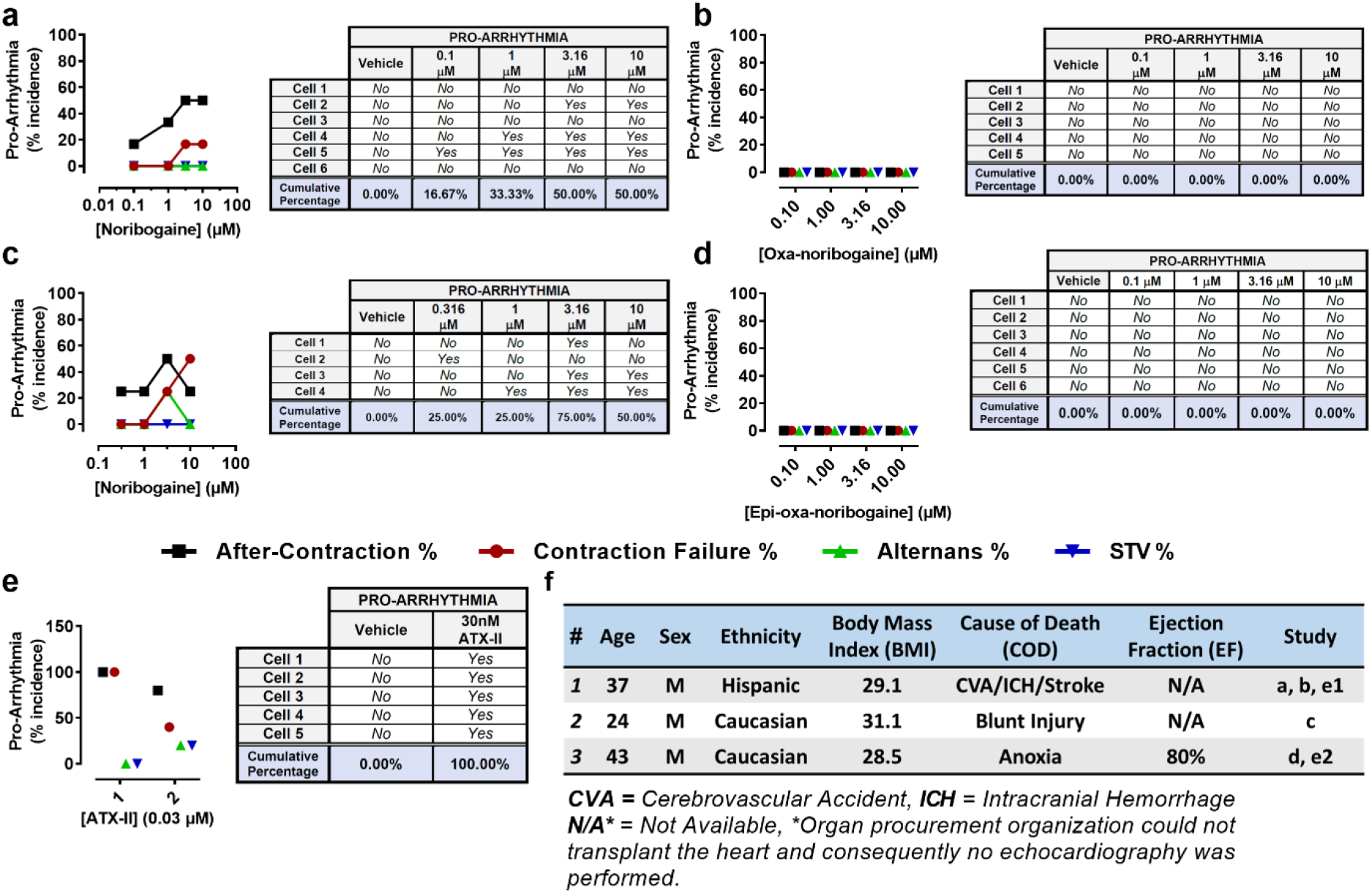
Pro-arrhythmia risk of noribogaine and oxa-iboga analogs examined in human primary cardiomyocytes. Noribogaine, **a**, was compared to oxa-noribogaine, **b**, using cardiomyocytes isolated from the same donor heart. **c**, Comparable and concentration-dependent pro-arrhythmia risk was detected for noribogaine across 2 donors. **d**, Epi-oxa-noribogaine shows no risk of pro-arrhythmia. **e**, Anemone toxin, ATX-II, was used as a positive control. **f**, Donor information.

**Extended Data Fig. 8.**
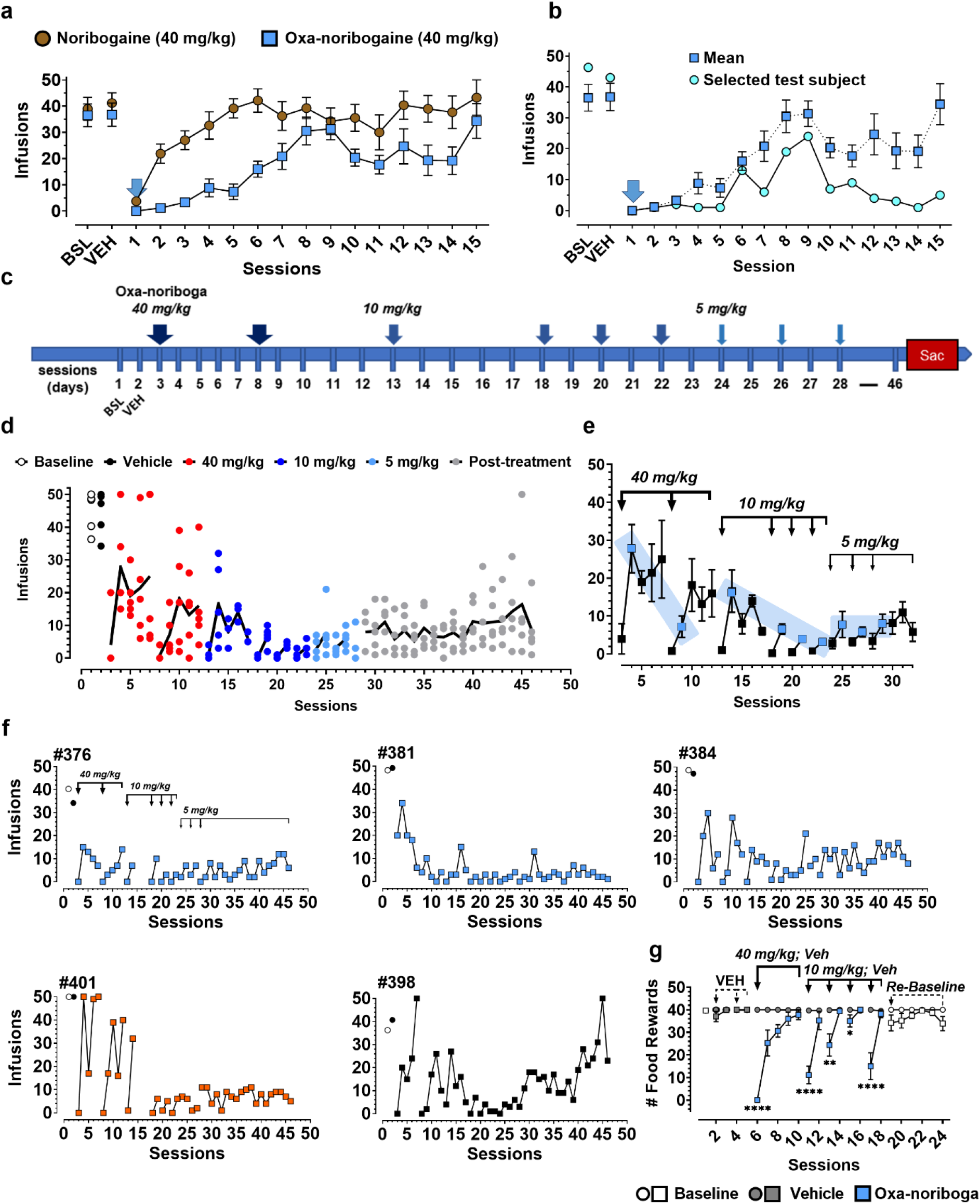
Morphine self-administration studies in rats. **a**, Complete time profile of a morphine SA study (Figure 4b) for noribogaine (40 mg/kg) and oxa-noribogaine (40 mg/kg) treatments. **b**, Morphine intake for one test subject that showed a profound and lasting morphine intake suppression (>14 days) after a single dose of oxa-noribogaine before session 1 (40 mg/kg, i.p.). The mean response of the entire cohort is shown for comparison. **c**, Experimental design of the repeated dosing regimen in morphine SA initiated by 40 mg/kg (“psychedelic reset dose”), followed by 10 and 5 mg/kg doses (“maintenance doses”) of oxa-noribogaine. **d**, Cumulative effect of the regimen of reset (40 mg/kg) and maintenance (10 and 5 mg/kg) doses of oxa-noribogaine resulted in sustained suppression of morphine intake. Individual data points are visualized with mean values depicted using solid lines. **e**, Section of panel **d** (or Figure 4e) where blue highlights indicate the morphine intake on days after oxa-noribogaine administration (after-effects), when oxa-noribogaine is essentially eliminated from the plasma and brain. Repeated administration of oxa-noribogaine increases its suppression efficacy on days after its administration, enabling dose tapering. **f**, Treatment response of individual subjects in the morphine SA study. Subjects *#376, #381* and *#384* strongly responded to the initial 40 mg/kg reset doses leading to a lasting suppression of morphine intake. Subject *#401* required multiple doses to subdue morphine intake and induce a long-term suppression, while subject *#398* responded to the treatment and showed lasting effects, but started to return toward basal morphine responding in the last 10 days of the experiment. **g**, Food maintained responding observed after a 40 mg/kg dose followed by four repeated 10 mg/kg doses of oxa-noribogaine. While the administration significantly suppresses food responding acutely (40 mg/kg: P < 0.0001, responses vary for repeated 10 mg/kg doses), the response of test subjects was no different than the control group subjects on the days after administration. No significant deviation of responding was observed between the test and the control group subjects over a 7 day period after the treatment was completed. Data are presented as mean ± SEM, specific statistical tests, information on reproducibility, and P values are reported in Methods and in Supplementary Statistics Table, \**P* < 0.05, \*\**P* < 0.01, \*\*\**P* < 0.001, \*\*\*\**P* < 0.0001.

**Extended Data Fig. 9.**
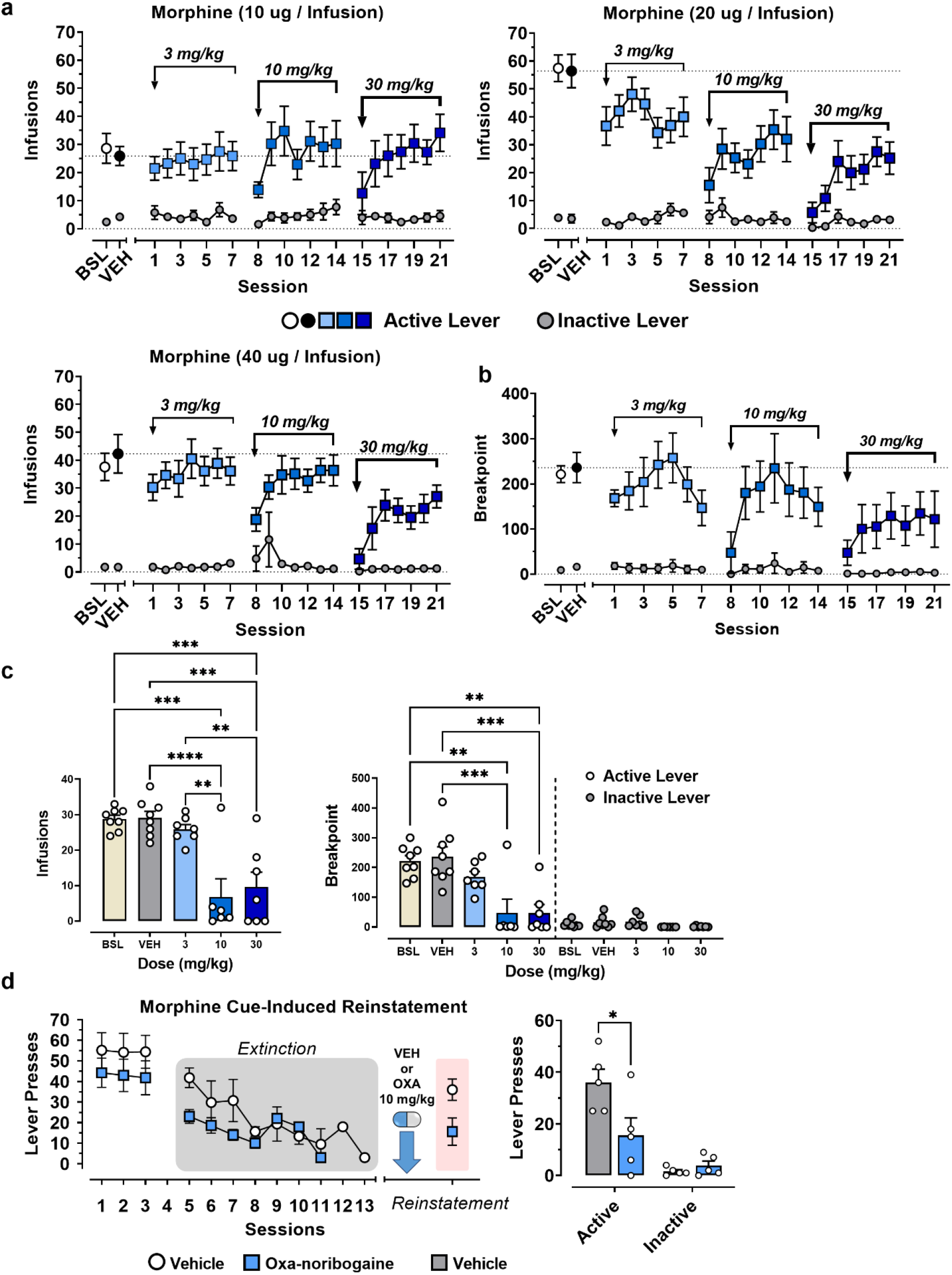
Morphine self-administration, extinction and reinstatement studies in rats. **a**, Complete time profiles of dose-response study utilizing escalating repeated doses of oxa-noribogaine (3, 10 and 30 mg/kg) in three animal test groups receiving different doses of morphine (10, 20 and 40 μg/inf). Both active and inactive lever responses are included. **b**, Time profile of a progressive ratio morphine SA study with both active and inactive lever responses included. **c**, Bar graph visualization of an acute effect of escalating dose of oxa-noribogaine (3, 10 and 30 mg/kg) on the morphine SA in the progressive ratio study. **d**, Time profile and active/inactive lever responses for the morphine cue-induced reinstatement study. Data are presented as mean ± SEM, specific statistical tests, information on reproducibility, and P values are reported in Methods and in Supplementary Statistics Table, \**P* < 0.05, \*\**P* < 0.01, \*\*\**P* < 0.001, \*\*\*\**P* < 0.0001.

