## Supplementary_Information for "Novel Class of Psychedelic Iboga Alkaloids Disrupts Opioid Use"

**Supplemental Material**
*(59 pages)*

#### Contents

Supplementary Tables 3

Methods 7

Data analysis and statistics 7

Broad Receptor Panel Assay 7

Commercial Assays 7

hSERT Inhibition Assay 7

Radioligand Competition Binding Assays (Mouse Opioid Receptors) 7

[$^{35}$S]GTPγS Functional Assay (Mouse Opioid Receptors) 8

BRET Functional Assays (G-protein, β-arrestin and Nb33 recruitment) 8

Cardiotoxicity Assay in Adult Human Primary Cardiomyocytes (AnaBios Corporation) 9

*In Vivo* Experiments 9

Materials and formulation 10

Tail-flick Test 11

Tail-flick Test (KO animals) 11

Tail-flick Test (antagonism pre-treatment) 11

Open Field Test 11

Forced Swim Test 12

Place Conditioning Preference 12

General procedures for intravenous drug self-administration and food responding studies in rodent animal models 12

Effect of oxa-noribogaine on morphine self-administration. 13

Effect of oxa-noribogaine on morphine self-administration under a progressive ratio schedule of reinforcement 14

Effect of oxa-noribogaine on cue-induced reinstatement of responding previously maintained by morphine or fentanyl 14

Effect of acute administration of oxa-noribogaine, epi-oxa-noribogaine and noribogaine on morphine self-administration 14
Effect of acute oxa-noribogaine administration on fentanyl self-administration 15
Food Maintained Responding 15
Effect of repeated administration of oxa-noribogaine on morphine self-administration 15
GDNF and mBDNF protein expression Studies 15
Pharmacokinetic Studies (PK, Sai Life Sciences Limited) 16
Plasma Protein and Brain Tissue Binding (Sai Life Sciences Limited) 16
Estimated free drug concentrations, target engagement, and correlation with in vitro pharmacology 17
Ibogaine Analogs Docking Studies on KOR 17
Synthesis 18
X-Ray structure determination of Oxa-ibogaine 10a. 27
Supplementary References 28
Appendix 1. High Resolution Mass Spectra 32
Appendix 2. NMR Spectra 36

### Supplementary Tables

Table S1: Binding affinities (*Radioligand Displacement*) [$K_i$, (95% CI) in nM] for oxa-iboga alkaloids at mouse opioid receptors (n = 3).

| Receptor | Oxa-noribogaine | Epi-oxa-noribogaine |
|---|---|---|
| ***mMOR*** | 86 (65; 115) | 136 (111; 166) |
| ***mKOR*** | 36 (29; 44) | 6.9 (5.7; 8.3) |
| ***mDOR*** | 168 (127; 223) | 94 (85; 104) |

*For radioligand experiments fitted curves were constrained to top = 100 and bottom = 0.*

Table S2: Agonist activity (*GTPγS assay*) [$EC_{50}$, (95% CI) in nM], {%$E_{max}$} of iboga alkaloids at mouse opioid receptors (n ≥ 3).

| Receptor | Oxa-noribogaine | Epi-oxa-noribogaine |
|---|---|---|
| ***mMOR*** | 559 (331, 920) {101} | 211 (139, 322) {111} |
| ***mKOR*** | 49 (37, 64) {92} | 9.6 (7.2, 12.8) {89} |
| ***mDOR*** | 329 (91, 117) {100} | 99 (75, 130) {108} |

*Efficacy data were obtained using agonist induced stimulation of [$^{35}$S]GTPγS binding assay. Efficacy is represented as $EC_{50}$ (nM) and percent maximal stimulation ($E_{max}$) relative to standard agonist DAMGO (MOR), DPDPE (DOR), or U50,488 (KOR) at 1000 nM.*

Table S3: Agonist activity (*G protein BRET assay*) [$EC_{50}$, (95% CI) in nM], {%$E_{max}$} of iboga alkaloids at human and rat kappa opioid receptor (n = 3).

| Receptor | (±)-U50,488 | Noribogaine | Oxa-noribogaine | Epi-oxa-noribogaine |
|---|---|---|---|---|
| ***hKOR*** | 13 (8, 20) {100} | 7,745 (284; 21,050) {46} | 68 (42; 109) {108} | 16 (10; 26) {96} |
| ***rKOR*** | 18 (11; 32) {100} | 6,239 (2,657; 14,650) {54} | 43 (17; 126) {78} | 12 (7; 23) {77} |

Table S4: Agonist activity (*Nb33 BRET assay*) [$EC_{50}$, (95% CI) in nM], {%$E_{max}$} of iboga alkaloids at human and rat kappa opioid receptor (n = 3).

| Receptor | (±)-U50,488 | Noribogaine | Oxa-noribogaine | Epi-oxa-noribogaine |
|---|---|---|---|---|
| ***hKOR*** | 176 (113, 284) {100} | n.d. {<3% at 10 μM} | 304 (171; 567) {46} | 58 (18; 336) {50} |
| ***rKOR*** | 202 (159; 257) {100} | n.d. {<4% at 10 μM} | 484 (304; 801) {55} | 62 (35; 112) {36} |

Table S5: β-Arrestin recruitment (*BRET assay*) [$EC_{50}$, (95% CI) in nM], {%$E_{max}$} of iboga alkaloids at human and rat kappa opioid receptor (n = 3), with and without addition of G-protein coupled receptor kinase 3 (GRK3).

| Receptor | (±)-U50,488 | Oxa-noribogaine | Epi-oxa-noribogaine |
|---|---|---|---|
| ***hKOR + GRK3*** | 86 (52, 141) {100} | 405 (216; 748) {65} | 49 (31; 108) {50} |
| ***rKOR + GRK3*** | 129 (64; 276) {100} | 610 (314; 997) {66} | 108 (60; 194) {47} |
| ***hKOR*** | 5,802 (1,515; 3,260,000) {100} | n.d. {13% at 100 μM} | n.d. {11% at 100 μM} |
| ***rKOR*** | 1,566 (841; 3,784) {100} | n.d. {11% at 100 μM} | n.d. {<0% at 100 μM} |

Table S6: Noribogaine functional assays [$EC_{50}$/$IC_{50}$, (95% CI) in nM], {%$E_{max}$} (n = 1).

| Assay (Species) | Receptor | Noribogaine |
|---|---|---|
| *GTPγS (human)* | *hKOR* (Agonist) | 8,325 (6143; 11,280) {73} |
| | *h5-HT$_{2A}$* (Agonist) | n.d. {3% at 25 μM} |
| *IP1 (human)* | *h5-HT$_{2A}$* (Agonist) | n.d. {<0% at 25 μM} |
| | *h5-HT$_{2B}$* (Agonist) | n.d. {<0% at 10 μM} |
| | *h5-HT$_{2B}$* (Antagonist) | n.d. {10% at 10 μM} |

Table S7: Agonist activity (*G protein BRET assay*) [$EC_{50}$ in nM], {%$E_{max}$} of oxa-ibogamine analogs demonstrating the importance of the phenolic -OH (position 10) on KOR activity (n = 4). Fitted curves were constrained to top = 100% and bottom = 0%.

| Assay (Species) | Receptor | Oxa-ibogamine | Epi-oxa-ibogamine |
|---|---|---|---|
| *G protein BRET (human)* | *hKOR* | >10,000 {56% at 100 µM} | ~10,000 {46% at 100 µM} |

Table S8: Off target agonist activity (*IP1 assay*) of oxa-noribogaine [$EC_{50}$ in nM], {%$E_{max}$} at human serotonin 2 receptors (n = 1).

| Assay (Species) | Receptor | Oxa-noribogaine |
|---|---|---|
| *IP1 (human)* | *h5-HT$_{2A}$* (Agonist) | n.d. {2% at 10 µM} |
| | *h5-HT$_{2B}$* (Agonist) | n.d. {<0% at 10 µM} |
| | *h5-HT$_{2C}$* (Agonist) | n.d. {<0% at 10 µM} |

Table S9: Binding affinities for hNOP, rNMDAR and inhibition of hnAChR (α3β4) ion channels [$IC_{50}$, (95% CI) in nM] by iboga alkaloids (n = 1). Fitted curves were constrained to top = 100% and bottom = 0%.

| Assay | Receptor | Noribogaine | Oxa-noribogaine | Epi-oxa-noribogaine |
|---|---|---|---|---|
| *Radioligand Displacement* | *hNOP* | n.d. {<0% at 10 µM} | n.d. {<0% at 10 µM} | n.d. {5% at 10 µM} |
| | *rNMDAR* | 42,220 (35,820; 49,770) | 24,060 (17,500; 33,090) | 66,800 (63,950; 69,790) |
| *Electrophysiology* | *hnAChR (α3β4)* | 4,960 (3,343; 7,360) | 2,891 (2,622; 3,188) | 1,495 (1,041; 2,147) |

Table S10: Inhibitory activity [$IC_{50}$ (95% CI) in nM] {%$E_{max}$} of iboga alkaloids at human serotonin transporter (hSERT, n = 3).

| Assay | Imipramine | Noribogaine | Oxa-noribogaine | Epi-oxa-noribogaine |
|---|---|---|---|---|
| *Fluorescence Uptake Inhibition* | 8.9 (7.0; 11) {100} | 286 (216; 372) {104} | 711 (560; 908) {106} | 1,707 (1,282; 2,394) {120} |

Table S11: Binding affinities [$K_i$ ± SEM, nM] for targets identified by broad PDSP screening of oxa-noribogaine. Exact Ki values were determined only for targets identified in the primary binding screening.

| Primary Binding Screen at 10 µM | $K_i$ ± SEM (Determined only for ligand displacement >50% at 10 µM) | | |
|---|---|---|---|
| *5-HT1A, 5-HT1B, 5-HT1D, 5-HT1E, 5-HT2A, 5-HT2B, 5-HT2C, 5-HT3, 5-HT4, 5-HT5A, 5-HT6, 5-HT7A, A1, A2A, α1A, α1B, α1D, α2A, α2B, α2C, β1, β2, β3, BZP, D1, D2, D3, D4, D5, DAT, DOR, GABAA, H1, H2, H3, H4, KOR, M1, M2, M3, M4, M5, MOR, NET, PBR,SERT, σ1, σ2, AMPA, NR2B, M4 D, NTS1, mGluR5, α2β2, α2β4, α3β2, α3β4, α4β2, α4β2 (Rat Brain), α4β4, α7, NOP, OT, V1A, V1B, V2, BZP Rat Brain Site* | **α2A (n = 3)** | **α2C (n = 3)** | **α3β2 (n = 1)** |
| | 1,452 ± 538 | 2,435 ± 476 | ~2,207 |
| | **α3β4 (n = 1)** | **α4β4 (n = 1)** | **α7 (n = 1)** |
| | ~2,802 | ~3,485 | ~1,293 |
| | **(σ2 (n = 3)** | **SERT (n = 3)** | **D5 (n = 3)** |
| | 4,488 ± 358 | 1,157 ± 391 | 4,217 ± 866 |
| | **DOR (n = 3)** | **KOR (n = 3)** | **MOR (n = 3)** |
| | 411 ± 30 | 1.6 ± 0.3 | 56 ± 15 |

Table S12: Pharmacokinetics data determined for noribogaine in male C57BL/6 mice following a single subcutaneous administration (Dose: 10 mg/kg, n = 3).

| Matrix | $T_{max}$ (h) | $C_{max}$ (ng/mL)[a] | $AUC_{last}$ (h×ng/mL) | $AUC_{inf}$ (h×ng/mL) | $T_{1/2}z$ (h) | CLf (mL/min/kg) | VZ/F (L/kg) |
|---|---|---|---|---|---|---|---|
| **Plasma** | 0.50 | 1979.32 | 5221.91 | 5229.14 | 2.76 | 31.87 | 7.62 |
| **Brain** | 0.50 | 5093.45 | 11672.46 | 12112.71 | 1.71 | 13.76 | 2.04 |

[a] *Brain conc. expressed as ng/g, density of brain tissue was considered as 1 which is equivalent to plasma density.*

Table S13: Pharmacokinetics data determined for oxa-noribogaine in male C57BL/6 mice following a single subcutaneous administration (Dose: 10 mg/kg, n = 3).

| Matrix | $T_{max}$ (h) | $C_{max}$ (ng/mL)[a] | $AUC_{last}$ (h×ng/mL) | $T_{1/2}z$ (h) | CLf (mL/min/kg) | Vd/F (L/kg) | Brain-Kp($C_{max}$) | Brain-Kp($AUC_{last}$) |
|---|---|---|---|---|---|---|---|---|
| **Plasma** | 0.25 | 581.24 | 1413.06 | 6.10 | 112.73 | 59.53 | N/A | N/A |
| **Brain** | 0.50 | 4492.48 | 7227.68 | 4.76 | N/A | N/A | 7.72 | 5.11 |

[a] *Brain conc. expressed as ng/g, density of brain tissue was considered as 1 which is equivalent to plasma density.*

Table S14: Pharmacokinetics data determined for oxa-noribogaine in male Wistar rats following a single intraperitoneal administration (Dose: 40 mg/kg, n = 3).

| Matrix | $T_{max}$ (h) | $C_{max}$ (ng/mL)[a] | $AUC_{last}$ (h×µg/mL) | $T_{1/2}z$ (h) | CLf (mL/min/kg) | Vd/F (L/kg) | Brain-Kp($C_{max}$) | Brain-Kp($AUC_{last}$) |
|---|---|---|---|---|---|---|---|---|
| **Plasma** | 0.08 | 1973.00 | 3520 | 1.26 | 189.29 | 20.62 | N/A | N/A |
| **Brain** | 0.08 | 18919.68 | 55220 | 3.59 | N/A | N/A | 9.60 | 15.69 |

[a] *Brain conc. expressed as ng/g, density of brain tissue was considered as 1 which is equivalent to plasma density.*

Table S15: Rat plasma protein binding of Oxa- and Epi-oxa-noribogaine (n = 3).

| Compound | % Bound ± SD | % Unbound | % Recovery | % Stability (4 h) |
|---|---|---|---|---|
| **Oxa-noribogaine** | 93.4 ± 0.4 | 6.6 | 94 | 104 |
| **Epi-oxa-noribogaine** | 86.4 ± 1.9 | 13.6 | 88 | 111 |

Table S16: Human plasma protein binding of noribogaine, Oxa- and Epi-oxa-noribogaine (n = 3).

| Compound | % Bound ± SD | % Unbound ± SD | % Recovery | % Stability (4 h) |
|---|---|---|---|---|
| **Noribogaine** | 59.9 ± 3.7 | 40.1 | 75 | 105 |
| **Oxa-noribogaine** | 95.8 ± 0.9 | 4.2 | 97 | 116 |
| **Epi-oxa-noribogaine** | 89.4 ± 0.5 | 10.6 | 98 | 108 |

Table S17: Rat brain tissue binding of Oxa- and Epi-oxa-noribogaine (n = 3).

| Compound | % Bound ± SD | % Unbound | % Recovery | % Stability (4 h) |
|---|---|---|---|---|
| **Oxa-noribogaine** | 98.9 ± 0.1 | 1.1 | 120 | 80 |
| **Epi-oxa-noribogaine** | 98.6 ± 0.1 | 1.4 | 106 | 97 |

Table S18: Tail-flick test characterization of antinociceptive properties of U50,488, oxa- and epi-oxa-noribogaine [Base Latency, (95% CI) in s; $ED_{50}$, (95% CI) in mg/kg], in male WT C57BL/6 mice.

| Compound | (±)-U-50,488 (n = 5)* | Oxa-noribogaine (n = 20) | Epi-oxa-noribogaine (n = 9) |
|---|---|---|---|
| **Base Latency (s)** | 3.7 (3.2; 4.1) | 2.6 (2.5; 3.9) | 2.0 (1.2; 2.7) |
| **$ED_{50}$ (mg/kg)** | 2.2 (1.6; 3.1) | 3.0 (2.6; 3.4) | 1.9 (1.4; 2.5) |

**5 naive animals were used for each dose, baseline latency was determined for each test subject*

Table S19: Tail-flick test characterization of antinociceptive properties of U50,488, oxa- and epi-oxa-noribogaine [Base Latency, (95% CI) in s; $ED_{50}$, (95% CI) in mg/kg], in female WT C57BL/6 mice.

| Compound | (±)-U-50,488 (n = 5)* | Oxa-noribogaine (n = 10) | Epi-oxa-noribogaine (n = 10) |
|---|---|---|---|
| **Base Latency (s)** | 2.9 (2.6; 3.2) | 2.3 (1.8; 2.8) | 2.3 (1.7; 2.9) |
| **$ED_{50}$ (mg/kg)** | 6.3 (4.4; 9.0) | 4.9 (3.9; 6.3) | 9.7 (6.9; 13.7) |

**5 naive animals were used for each dose, baseline was determined for each test subject*

Table S20: Tail-flick test comparison of antinociceptive properties of oxa- and epi-oxa-noribogaine in wild type (WT), mu (MOR-KO) and kappa (KOR) knockout (KO) mouse models [baseline, (95% CI) in s; $ED_{50}$, (95% CI) in mg/kg].

**Oxa-noribogaine**

| **Sex** | **Male** | | | **Female** | |
|---|---|---|---|---|---|
| **Model** | **WT (n = 10)** | **KOR-KO (n = 8)** | **MOR-KO (n = 6)** | **WT (n = 5)** | **KOR-KO (n = 4)** |
| **Base Latency (s)** | 2.5 (2.3, 2.7) | 2.7 (2.4, 3.0) | 2.8 (2.4, 3.1) | 2.9 (2.4, 3.3) | 3.5 (2.6, 4.4) |
| **$ED_{50}$ (mg/kg)** | 3.1 (2.5; 3.8) | 20.6 (16.5; 25.7) | 7.0 (4.3; 11.3) | 3.1 (2.5; 3.8) | 20.6 (16.5; 25.7) |

**Epi-oxa-noribogaine**

| **Sex** | **Male** | | |
|---|---|---|---|
| **Model** | **WT (n = 5)** | **KOR-KO (n = 5)** | **MOR-KO (n = 7)** |
| **Base Latency (s)** | 2.3 (1.9, 2.6) | 2.2 (2.0, 2.4) | 3.4 (2.7, 4.0) |
| **$ED_{50}$ (mg/kg)** | 1.3 (0.9; 1.7) | n.d. $\gg$100 | 2.0 (1.5; 2.6) |

### Methods

##### Data analysis and statistics

Nonlinear curve fitting and statistical analyses were performed using models available in GraphPad Prism (9.4 or 8.3, San Diego, CA), unless otherwise noted. Data are represented as mean ± SEM, unless stated otherwise, with asterisks indicating significance $*P < 0.05$, $**P < 0.01$, $***P < 0.001$ and $****P < 0.0001$.

##### Broad Receptor Panel Assay

Binding constants ($K_i$) at the selected human receptors, ion channels and transporters were generously determined using radioligand displacement experiments by the National Institute of Mental Health's Psychoactive Drug Screening Program, Contract # HHSN-271-2008-00025-C (NIMH PDSP).[1] The NIMH PDSP is Directed by Bryan L. Roth MD, PhD at the University of North Carolina at Chapel Hill and Project Officer Jamie Driscoll at NIMH, Bethesda, MD, USA. For experimental details please refer to the PDSP website http://pdsp.med.unc.edu/.

##### Commercial Assays

KOR [$^{35}$S]GTP$\gamma$S agonist assay (for noribogaine), rat NMDA (Glutamate, MK-801) and human NOP (Orphanin ORL1) radioligand displacements, human nicotinic acetylcholine receptor (nAChR α3β4) inhibition studies were performed by contracted research organization (items: 333200, 233010, 260600 and CYL8057IF2). Experiments were conducted as described on https://www.eurofinsdiscoveryservices.com/.

##### hSERT Inhibition Assay

Stably transfected hSERT-HEK cellular cultures were maintained in Dulbecco's Minimal Essential Medium (DMEM) with GlutaMAX (Gibco) with the following additions: 10% (v/v) Fetal Bovine Serum (FBS, Atlanta Biologicals), 100 U/mL Penicillin 10 µg/mL Streptomycin (Gibco) and 500 µg/mL Geneticin (G418) (Sigma). Singly transfected cells were seeded at a density of $0.09 \times 10^6$ cells/well in poly-D-Lysine (Sigma) coated white solid-bottom 96-well plates (Greiner). Cells were grown for 44 hours in aqueous media at 37 °C under 5% $CO_2$ atmosphere. At the beginning of the experiment, the cellular growth solution was aspirated, and individual cells were rinsed with 150 µL of 1 × Dulbecco's Phosphate Buffered Saline (PBS; HyClone). 63 µL of Experimental Media (DMEM without phenol red but with 4.5 g/L of D-Glucose (Gibco), 1% (v/v) FBS (Atlanta Biologicals), 100 U/mL Penicillin, and 10 µg/mL Streptomycin (Gibco)) with 2 × tiered concentrations of inhibitor (vehicle DMSO and control inhibitor: Imipramine) were added.[2] After pre-incubation period (one hour), 63 µL of Experimental Media containing 2 × various concentrations of tested inhibitor (or vehicle) along with a specified amount of fluorescent substrate $APP^+$ (1.1 µM)[3] was added to the wells. Fluorescent probe was allowed to uptake for 30 min, the contents of each well were aspirated and consequently, rinsed twice with 120 µL of PBS. A final solution of 120 µL of PBS was finally added to all corresponding wells for cell maintenance before measuring fluorescence uptake (BioTek H1MF plate reader, $APP^+$ excitation and emission wavelengths 417 and 502 nm). Recorded inhibitor values were first subtracted from vehicular values to quantify the respective fluorescence uptake. Data were analyzed using the dose-response-inhibitor nonlinear curve fitting model (log[inhibitor] vs response (four parameters).

##### Radioligand Competition Binding Assays (Mouse Opioid Receptors)

Assays were performed as previously reported.[4] Briefly, IBNtxA and [$^{125}$I]BNtxA were synthesized at MSKCC as previously described.[5,6,8] Na$^{125}$I was purchased from Perkin-Elmer (Waltham, MA). [$^{125}$I]BNtxA binding was carried out in membranes prepared from Chinese Hamster Ovary (CHO) cells stably expressing murine clones of MOR, DOR, and KOR, as previously described.[5–7] Binding incubations were performed at 25 °C for 90 min in 50 mM potassium phosphate buffer, pH 7.4, containing 5 mM magnesium sulfate. After the incubation, the reaction was filtered through glass-fiber filters (Whatman Schleicher & Schuell, Keene, NH) and washed three times with 3 mL of ice-cold 50 mM Tris-HCl (pH 7.4) on a semiautomatic cell harvester. Nonspecific binding was defined by addition of levallorphan (8 µM) to matching samples and was subtracted from total binding to yield specific binding. Protein concentrations were determined using the

Lowry method with BSA as the standard.[9] Data were analyzed using the binding-competitive nonlinear curve fitting model (One site - Fit $K_i$).

##### [$^{35}$S]GTPγS Functional Assay (Mouse Opioid Receptors)

Assays were performed as previously reported.[4] Briefly, [$^{35}$S]GTPγS binding was performed on membranes prepared from transfected cells stably expressing opioid receptors in the presence and absence of the indicated compound for 60 min at 30 °C in the assay buffer (50 mM Tris-HCl (pH 7.4) 3 mM $MgCl_2$, 0.2 mM EGTA, and 10 mM NaCl) containing 0.05 nM [$^{35}$S]GTPγS; 2 μg/mL each leupeptin, pepstatin, aprotinin, and bestatin, and 30 μM GDP, as previously described.[10] After the incubation, the reaction was filtered through glass fiber filters (Whatman Schleicher & Schuell, Keene, NH) and washed three times with 3 mL of ice-cold buffer (50 mM Tris-HCl, pH 7.4) on a semiautomatic cell harvester. Filters were transferred into vials with 3 mL of Liquiscint (National Diagnostics, Atlanta, GA), and the radioactivity in vials was determined by scintillation spectroscopy in a Tri-Carb 2900TR counter (PerkinElmer Life and Analytical Sciences). Basal binding was determined in the presence of GDP and the absence of drug. Data were normalized to 1000 nM DAMGO, DPDPE, and U50,488 for MOR, DOR, and KOR binding, respectively. Data were analyzed using the dose-response-stimulation nonlinear curve fitting model (log[agonist] vs. response (three parameters)).

##### BRET Functional Assays (G-protein, β-arrestin and Nb33 recruitment)

*Material:* HEK-293T cells were obtained from the American Type Culture Collection (Rockville, MD) and were cultured in a 5% $CO_2$ atmosphere at 37 °C in Dulbecco's Modified Eagle Medium (high glucose #11965; Life Technologies Corp.; Grand Island, NY) supplemented with 10% Fetal Bovine Serum (FBS, #35-010-CV, Corning, Corning, NY, USA), 100 U·mL$^{-1}$ penicillin (#30-002-CI, Corning, Corning, NY, USA), and 100 μg·mL$^{-1}$ streptomycin (#30-002-CI; Corning, Corning, NY, USA). The following chemicals were used without further modification: coelenterazine H (#DC-001437, Dalton Pharma Services, Toronto, ON, Canada), PEI (#NC1014320, Polysciences, Warrington, PA, USA) and (±)-U-50488 HCl (Tocris Biosciences, Minneapolis, MN, USA).

*DNA Constructs (G-protein, β-arrestin, and Nb33):* The rat KOR (rKOR) was provided by Dr. Lakshmi Devi at Mount Sinai School of Medicine. The human KOR (hKOR) was obtained from the Missouri S&T Resource Center. The GRK3, $G\alpha_{oB}$ with Renilla luciferase 8 (RLuc8) inserted at position 91 ($G\alpha_{oB}$-RLuc8), and $G\beta_1$ ($\beta_1$) were provided by C. Galés.[11,12] Venus-Arrestin2 and $G\gamma_2$, which was fused to the full-length mVenus at its N-terminus via the amino acid linker GSAGT (mVenus-γ2), were constructed in house. The expression vectors coding for rat and human KOR tagged at the C-terminus with Nanoluc (KOR-nluc) were constructed using standard techniques in molecular biology and confirmed by DNA sequencing (Genewiz, South Plainfield, NJ, USA). Briefly, three DNA inserts were PCR amplified, one coding for the N-terminal signal peptide and flag tag, one coding for KOR, and one coding for nanoluc. The inserts were ligated and cloned into a pcDNA3.1 (+) vector (#V79020, ThermoFisher Scientific, Waltham, MA, USA). The plasmid coding for the nanobody33Venus (Nb33) construct[13] was a gift from Dr. Meritxell Canals at the University of Nottingham.

*Transfection (G-protein, β-arrestin, and Nb33):* The following cDNA amounts were transfected into HEK-293T cells (4 x $10^6$ cells/plate) in 10-cm dishes using polyethylenimine (PEI) in a 1.5:1 ratio (diluted in DMEM, Life Technologies). *G-protein β-γ release*: 2.5 μg KOR, 0.1 μg $G\alpha_{oB}$RLuc8, 6.2 μg $\beta_1$, 6.2 μg mVenus-γ2. *Arrestin recruitment*: 0.2 μg KOR-nluc, 15 μg Arrestin-2-Venus, with/without 5 μg GRK3. Cells were maintained in the HEK-293T media described above. After 24 hours the media was changed, and the experiment was performed 48 hours after transfection. *Nb33 recruitment:* A total of 5 μg of cDNA was transiently transfected into HEK-293T cells (2 × $10^6$ cells per plate) in 10 cm dishes (1 μg receptor-nluc, and 4 μg Nb-33-Venus), using PEI in a 6:1 ratio (diluted in DMEM). Cells were maintained in the HEK-293T media described above. Experiments were performed 48 hours after transfection.

*BRET:* Transfected cells were dissociated and re-suspended in phosphate-buffered saline (PBS). Approximately 200,000 cells/well were added to a black-framed, white-well, 96-well plate (#60050; Perkin Elmer; Waltham, MA). At time zero, the luciferase substrate coelenterazine H (5 μM) was added to each well. Ligands were added after 5 min, then BRET signal was measured 5 min later for G-protein, and 10 min later for Nb33 and β-arrestin recruitment. BRET measurements were performed using a PHERAstar FS plate reader (BMG Labtech, Cary, NC, USA). The BRET signal was calculated as the ratio of the light emitted by the mVenus acceptor (510–540 nm) over the light emitted by the NanoLuc donor (475 nm). This drug-induced BRET signal was normalized using the $E_{max}$ of U-50,488 as the maximal response at KOR.

Data were analyzed using the dose-response-stimulation nonlinear curve fitting model (log[agonist] vs. response (four parameters)). All experiments were repeated in three independent trials each with triplicate determinations.

**Cardiotoxicity Assay in Adult Human Primary Cardiomyocytes (AnaBios Corporation)**

Cardiotoxicity of noribogaine and oxa-iboga analogs was assessed according to published procedure.[14,15] Briefly, adult human primary ventricular myocytes were isolated from ethically consented donor hearts that were enzymatically digested using a proprietary protocol. Cardiomyocytes were placed in a perfusion chamber mounted on the stage of inverted Motic AE31E (IonOptix) or Olympus IX83P1ZF (MyoBLAZER) microscope and continuously perfused at approximately 2 mL/min with recording buffer heated to 35 ± 1 °C using an in-line heater from Warner Instruments (IonOptix & MyoBLAZER) and allowed to equilibrate for 5 minutes under constant perfusion. The cells were field stimulated with supra-threshold voltage at a 1 Hz pacing frequency, with a bipolar pulse of 3 ms duration, using a pair of platinum wires placed on opposite sides of the chamber connected to a MyoPacer stimulator. Starting at 1 V, the amplitude of the stimulating pulse was increased until the cardiomyocytes started generating contractility transients, and a value 1.5× threshold was used throughout the experiment. Cardiomyocytes were then imaged at 240 Hz using an IonOptix MyoCam-S CCD camera (IonOptix) or at 148 Hz using an Optronis CP70-16-M/C-148 (MyoBLAZER) camera. Digitized images were displayed within the IonWizard acquisition software (IonOptix) or MyoBLAZER acquisition software. The longitudinal axis of the selected cardiomyocyte was aligned parallel to the video raster line, by means of a cell framing adapter. Optical intensity data was collected from a user-defined rectangular region placed over the cardiomyocyte image. The optical intensity data represented the bright and dark bands corresponding to the Z-lines of the cardiomyocyte. The IonWizard software or MyoBLAZER Analysis software analyzed the periodicity in the optical density of these bands by means of a fast Fourier transform algorithm.

Compound test solutions were formulated from stock solutions within 30 min prior to experimental application to the cells. Test solutions were applied after vehicle control (120 s interval, 1 Hz stimulation) in an increasing concentration order (in 300 s intervals, 1 Hz stimulation) and experiment was terminated after wash control (300 s interval, 1 Hz stimulation).

Positive control 30 nM ATX-II (toxin from anemonia sulcate) was applied after vehicle control (120 s interval, 1 Hz stimulation) and the data were recorded (300 s interval, 1 Hz stimulation).

An aftercontraction (AC) was visually identified as spontaneous secondary change in the slope of the contractility transient that occurred before the next stimulus-induced contraction and that produced an abnormal and unsynchronized contraction. Contraction Failure (CF) was also visually identified when an electrical stimulus was unable to induce a contraction. Alternans and Short-Term Variability (STV) are visualized in Poincaré plots of Contraction Amplitude variability. STV ($STV = \Sigma|CAn+1-CAn|\ (20\times\sqrt{2})^{-1}$) was calculated with the last 20 transients of each control and test article concentration period. Alternans were identified as repetitive alternating short and long contractility amplitude transients. STV values were normalized to the vehicle control value of each cell. AC, CF and Alternans were plotted and expressed as % of incidence of cells exhibiting each of the signals normalized by the total number of cardiomyocytes.

*Noribogaine concentration selection:* The concentration range was selected on the basis of these considerations: the free plasma noribogaine $C_{max}$ values were estimated from the reported clinical data of total plasma concentrations[16] and our own human plasma protein binding data (Extended data Fig. 3g); approximately 100 nM noribogaine (60 mg oral dose) produced a mild QT effect and represents a relatively safe plasma level, while 300-400 nM (180 mg oral dose) induced concerning levels of QT prolongation (>500 ms), whereas low micromolar concentrations can be reached after detox therapeutic doses of ibogaine (>8 mg/kg) which have been associated with risk of arrhythmias.

***In Vivo* Experiments**

All experimental procedures involving animals were approved by the Columbia University, Memorial Sloan Kettering Cancer Center, Rutgers University or High Point University Institutional Animal Care and Use Committee (IACUC) and adhered to principles described in the National Institutes of Health Guide for the Care and Use of Laboratory Animals. Animals received regular veterinary care (weekly by institutional veterinarians) including daily health monitoring (by experimenters) of the animals (observing home cage behaviors, nesting, and body weight). All procedures were designed to minimize any stress/distress.

*Tail-flick, open-field and forced swim test*: Healthy adult C57BL/6J mice male (8-15 weeks, 29-35 g) and female (8-15 weeks, 19-26 g) were purchased from the Jackson Laboratory (Bar Harbor, ME) and housed

5 mice per cage with food and water available *ad libitum*. Mice were maintained on a 12-h light/dark cycle (lights on 7:00-19:00) and all testing was done in the light cycle. Temperature was kept constant at 22 ± 2 °C, and relative humidity was maintained at 50 ± 5%.

*Conditioned placed preference:* Healthy adult male and female C57BL/6 mice (10-15 weeks, 25-32 g) were purchased from the Jackson Laboratory (Bar Harbor, ME) and housed in groups of 4-6 in AllerZone MicroIsolator cages placed in RAIR HD Enviro-Gard™ Ventilated racks with HEPA Filtered Air Units with food and water available *ad libitum*. Vivarium conditions were recorded daily, and the automated climate control ensured constant temperature in the range of 20-24 °C and humidity 40-60%. Mice were maintained on a 12-h light/dark cycle with lights on at 18:00 and experimental sessions took place during the dark phase of the cycle.

*Self-administration and neurotropic factors expression studies*: Adult male Fisher F-344 rats (CDF Strain, 90-150 days, 230-280 g) were purchased from Charles River Laboratories (Wilmington, MA) and housed in groups of 2 in AllerZone MicroIsolator cages placed in RAIR HD Enviro-Gard™ Ventilated racks with HEPA Filtered Air Units prior to surgery. Following surgery rats were housed individually in acrylic cages with food and water available *ad libitum*. Vivarium conditions were recorded daily, and the automated climate control ensured constant temperature in the range of 20-24 °C and humidity 40-60%. Rats were maintained on a 12-h light/dark cycle with lights on at 18:00, and experimental sessions took place during the dark phase of the cycle.

*Pharmacokinetic studies*: Healthy adult male C57BL/6 mice (8-12 weeks, 18-36 g) or healthy adult male Wistar rats (8-12 weeks, 250-280 g) were procured from Global (India). Three mice or rats were housed in each cage. Temperature and humidity were maintained at 22 ± 3 °C and 30-70%, respectively and illumination was controlled to give a sequence of 12 h light and 12 h dark cycle. Temperature and humidity were recorded by auto-controlled data logger system. All animals were provided laboratory rodent diet (Envigo Research private Ltd, Hyderabad). Reverse osmosis water treated with ultraviolet light was provided *ad libitum*.

**Materials and formulation**

Drugs for *in vivo* experiments were either purchased from commercial sources and used as received or synthesized in the Sames laboratory as described in the synthesis section (noribogaine and oxa-iboga analogs - prepared by the nickel-mediated coupling). Commercial chemicals: U50,488 hydrochloride (Tocris Bioscience), cyprodime hydrochloride (Tocris Bioscience), naloxone hydrochloride (Alfa Aesar), naltrindole hydrochloride (MedChemExpress), imipramine hydrochloride (Alfa Aesar), aticaprant free base (MedChemExpress), morphine sulfate (Gallipot, Inc., St. Paul, MN), penicillin G procaine (Butler Company, Columbus, OH), propofol (Patterson Veterinary Supply, Inc., Loveland, CO), fentanyl citrate (Fagron, Inc., St Paul, MN), ketamine hydrochloride (Ketaset), xylazine (Xylamed, Patterson Veterinary Supply, Inc., Greely, CO).

*Pharmacology experiments:* The compound solutions were prepared on the same day as testing from pure solid material. Solids were dissolved in UPS grade 0.85% saline with addition of 2 molar equivalents of glacial acetic acid. Heat and sonication were used to assist in fully dissolving the solid to obtain a clear solution. Aticaprant was solubilized using 7% (v/v) Tween 80 in saline with addition of 2 molar equivalents of glacial acetic acid. Compound solutions were prepared at a concentration allowing for a volume of injection (190 - 350 µL) based on the body weight of the animal.

*Self-administration and protein expression experiments*: Test compounds (noribogaine and oxa-iboga) were dissolved in vehicle (2 molar equivalents of acetic acid in water) at a concentration of (20 mg/mL) to allow for injection volume of (2 mL/kg). For example, a 40 mg/kg dose administered to a 300 g rat required injecting 12 mg of test substance in 0.6 mL of vehicle (0.5mg/mL/kg). Heat and sonication were used to assist in fully dissolving the solid to obtain a clear solution.

*Pharmacokinetic studies:* Noribogaine hydrochloride for subcutaneous (s.c.) administration was dissolved in DMSO and the solution was diluted with normal saline to obtain formulation in 5% DMSO, 95% (v/v) normal saline (formulation strength 2 mg/mL). The final mixture was vortexed and sonicated to obtain a clear solution. Oxa-noribogaine was solubilized using 0.85% saline with addition of 2 molar equivalents of glacial acetic acid, formulations were vortexed and sonication to obtain a clear solution (formulation strength 2 mg/mL for s.c. and 4 mg/mL for i.p.).

**Tail-flick Test**

Tests were performed as previously reported.[17] Mice were moved to the testing room 30 minutes before the experiment to allow for acclimation. The body weight of each mouse and base tail-flick value were recorded. Mice were administered a 1 mg/kg s.c. dose of compound solution. After injection mice were returned to the home cage and allowed to rest for 30 minutes. Thirty minutes post injection the tail-flick measurement was taken using thermal stimulation via IR on a Ugo Basile unit set to 52 PSU (to achieve a baseline between 2 and 3 seconds and 10 seconds was used as a maximum latency to prevent tissue damage). Mice were then administered 3 mg/kg s.c. dose, allowed to rest for 30 minutes, followed by another tail-flick measurement. This process was repeated for doses 10 and 30 mg/kg in increasing order. Tail-flick latencies for the different doses were expressed as percentage of maximum potential effect (% MPE) by subtracting the experimental value by the base tail flick value then dividing by the difference between the maximum possible latency (10 seconds) and the base tail-flick value and finally multiplying by 100.

*Note*: $ED_{50}$ value for U50,488 was determined using four separate groups of naive mice (5 animals for each dose, baseline latency was determined for each animal prior to drug administration), as cumulative dosing of U50,488 in the same animal resulted in a rapid development of tolerance.

*Data Analysis:* $ED_{50}$ values were calculated using the dose-response-stimulation nonlinear curve fitting model (log[agonist] vs. response (four parameters)) with applied constraints Top = 100 and Bottom = 0.

**Tail-flick Test (KO animals)**

For analgesic testing in knockout animals MOR-1 Exon-1 KO[18] and KOR-1 Exon-3 KO[19] mice on a C57 background were bred in the Pintar laboratory at Rutgers University. All mice used were opioid naïve. Analgesia was tested in wild-type and KO animals by the radiant heat tail-flick technique using an IITC Model 33 Tail Flick Analgesia Meter as previously described.[20,21] The intensity was set to achieve a baseline between 2 and 3 seconds. Tail flick antinociception was assessed as an increase in baseline latency, with a maximal 10 s latency to minimize damage to the tail. Data were analyzed as percent maximal effect, % MPE, which was calculated according to the formula: % MPE [(observed latency − baseline latency)/(maximal latency − baseline latency)] × 100. Compounds were administered subcutaneously (s.c.) and analgesia was assessed at the peak effect (15 minutes). Mice were tested for analgesia with cumulative subcutaneous doses of the drug until the mouse can withstand the maximal latency. Once the mouse reached the maximal latency, the mouse was no longer given higher doses.

*Data Analysis:* $ED_{50}$ values were calculated using the dose-response-stimulation nonlinear curve fitting model (log[agonist] vs. response (four parameters)) with applied constraints Top = 100 and Bottom = 0. Statistical significance (WT vs KO-model) was assessed using the extra-sum-of-squares F test (alpha = 0.05).

**Tail-flick Test (antagonism pre-treatment)**

Mice were transferred to the testing room 30 minutes before the experiment to allow for acclimation. The weight of each mouse and base tail-flick value were recorded. Mice were administered the selected antagonist s.c. Antagonists were prepared and used at the following doses: cyprodime 10 mg/kg, naloxone 10 mg/kg, naloxone 1 mg/kg, naltrindole 0.5 mg/kg, and aticaprant 0.1 mg/kg. After injection, mice were returned to the home cage and allowed to rest for a duration of time specific to the antagonist being used. The wait time after administration of antagonist are as follows: cyprodime 30 min, naloxone 30 min, naltrindole 45 min and aticaprant 20 min. Mice were then administered oxa-noribogaine s.c. prepared at the dose 10 mg/kg. After injection, mice were returned to the home cage to rest for a duration of time specific to the experimental compound being used. The wait time after administration of oxa-noribogaine was 30 min. Afterwards, the tail-flick measurement was taken as described above.

*Data Analysis:* Statistical analysis was performed using the unpaired t-test, two-tailed with Welch's correction.

**Open Field Test**

Mice were transferred to the testing room 30 minutes before the experiment to allow for acclimation. Body weight of each mouse was recorded. For experiments with antagonist, pre-treatment, the dosing and after application wait times were the same as for the tail-flick test. Mice were administered an $ED_{50}$, $ED_{80}$ or >$ED_{95}$ doses of test compound (volume of injection 290 - 350 µL based on body weight). After injection mice were returned to the home cage and allowed to rest for 30 minutes. Each mouse was then gently placed in the center of a clear Plexiglas arena (27.31 × 27.31 × 20.32 cm, Med Associates ENV-510) lit with dim light (~5 lux) and allowed to ambulate freely for 60 minutes. The locomotion of the mouse was tracked by infrared beams embedded along the X, Y, Z axes of the area and automatically recorded. Data was collected on Activity Monitor by Med Associates.

*Data Analysis:* Statistical analysis was performed using the unpaired t-test, two-tailed with Welch's correction to analyze total locomotion and a two-way ANOVA with Šidák's multiple comparison test for *post hoc* comparison to analyze locomotion across time bins.

**Forced Swim Test**

One week after arriving in the facility, mice were handled for approximately 5 minutes by a male experimenter. All experiments were carried out by the same male experimenter who performed the handling. On the day of FST testing, mice were weighed and transferred to the testing room 30 minutes prior to experimentation. Mice were administered 0.85% saline (s.c.), imipramine 15 mg/kg (i.p.), oxa-noribogaine $ED_{80}$ (5.4 mg/kg) or >$ED_{95}$ 10 mg/kg dose s.c. 15 minutes after receiving saline or imipramine injection, or 30 minutes after receiving oxa-noribogaine injection, each subject mouse was gently placed into cylinder (plexiglass 40 cm tall, diameter 22 cm) filled with water (height 18 cm) kept at 23.8 – 25.0 °C. Mice were allowed to swim for 6 minutes after which they were removed, dried, and returned to their home cage. Video footage was analyzed with Noldus FST scoring software to obtain immobility scores. The mouse's view of the FST cylinder was obstructed prior to initiating the experiment. Only the last 4 minutes of the 6 minutes test were analyzed for time spent immobile. All FSTs were performed between the hours of 12:00 and 18:00.

*Data Analysis:* Statistical analysis was performed using the unpaired t-test, two-tailed with Welch's correction.

**Place Conditioning Preference**

Place conditioning was conducted in a three-chamber apparatus with two equal sized chambers (16.8 × 12.7 × 12.7 cm) that differed in color (black, white) and flooring (mesh, bar), separated by a middle chamber (7.2 × 12.7 × 12.7 cm) with computer-controlled doors and equipped with infrared diodes (Med Associates). Mice were acclimated to the room for three days before the pretest phase.

*Pretest phase*: Mice were placed in the middle chamber; the doors were raised, and the animal had free access to the chamber for 20 minutes. Dependent measures included the amount of time spent in each compartment, activity in the compartments, and entries into the compartments. No statistically significant difference was observed in the amount of time spent between the compartments during the pretest phase. Using an unbiased approach, the drug-paired chamber was randomly assigned for each mouse. Mice spending more than 65% time in either compartment or 33% in the middle compartment during the pre-conditioning session were excluded from the study. *Conditioning phase:* Conditioning began the day following the pretest phase and consisted of three two-day pairing cycles. On the first, third and fifth days of conditioning, mice received either cocaine (10 mg/kg, i.p.), morphine (20 mg/kg, i.p.), oxa-noribogaine (10 or 40 mg/kg, i.p.) or epi-oxa-noribogaine (10 and 40 mg/kg, i.p.) and were confined to the drug paired compartment for 30 minutes. On the second, fourth and sixth days, mice were administered saline (1 mL/kg, i.p.) and confined to the vehicle paired compartment.

*Testing phase:* The following day, mice were placed in the middle compartment, doors were raised, and the mice were allowed access to the entire chamber for 15 minutes. Dependent measures were identical to the pretest phase.

*Data Analysis:* A CPP score was calculated as the difference in time spent in the drug-paired minus the vehicle paired compartment for both the pretest and testing phases. Data were analyzed using paired t-tests or two-way ANOVA with Šidák's multiple comparison test for *post hoc* comparison.

**General procedures for intravenous drug self-administration and food responding studies in rodent animal models**

*Operant self-administration apparatus:* Rats were transferred to operant conditioning chambers (ENV-008CT; Med Associates, St. Albans, VT) enclosed in sound-attenuating cubicles (ENV-018; Med Associates). The front panel of the operant chambers contained two response levers (4 cm above the floor and 3 cm from the side walls), a cue light (3 cm above the lever) and a food chute centered on the front wall (2 cm above the floor) that was connected to a food pellet dispenser (ENV-023; Med Associates) located behind the front wall and a tone generator to mask extraneous noise. A syringe pump (PHM-100; Med Associates) holding a 20-ml syringe delivered infusions. A counter-balanced arm containing the single channel liquid swivel was located 8-8.5 cm above the chamber and attached to the outside of the front panel. An IBM compatible computer was used for session programming and data collection (Med Associates Inc., East Fairfield, VT).

*Lever training:* Subjects were transferred to the operant chambers for daily experimental sessions and responding was engendered and maintained by delivery of food pellets (45 mg pellets: Noyes, Lancaster, NH) under a fixed ratio (FR 1) schedule of reinforcement. The lever lights were illuminated when the schedule was in effect. Completion of the response requirement extinguished lights, delivered food, and was followed by a 20-second timeout (TO) period during which all lights were extinguished, and responses had no scheduled consequences. After the TO, the lights were illuminated, and the FR schedule was again in effect. Sessions lasted 20 minutes or until 50 food pellets were delivered.

*Intravenous jugular surgery:* After operant responding was acquired and maintained by food, subjects surgically implanted with an intravenous jugular catheter. Venous catheters were inserted into the right jugular vein following administration of ketamine (90 mg/kg; i.p.) and xylazine (5 mg/kg; i.p.) for anesthesia as described previously.[22–24] Catheters were anchored to muscle near the point of entry into the vein. The distal end of the catheter was guided subcutaneously to exit above the scapulae through a Teflon shoulder harness. The harness provided a point of attachment for a spring leash connected to a single-channel fluid swivel at the opposing end. The catheter was threaded through the leash and attached to the swivel. The other end of the swivel was connected to a syringe (for saline and drug delivery) mounted on a syringe pump. Rats were administered penicillin G procaine (75,000 units in 0.25 mL, i.m.) and allowed a minimum of 5 days to recover before self-administration studies were initiated. Catheter patency was maintained by hourly infusions of 0.2 ml of 0.9% saline (w/v) with 1.7 U/ml of heparin equivalent to 0.34U heparin/infusion while in the home cage only. Infusions of propofol (6 mg/kg; i.v.) were manually administered as needed to assess catheter patency.

*Self-administration training:* Following recovery, rats were transferred to their respective operant chambers for daily self-administration sessions. Before each session, the swivel and catheter were flushed with 500 µL of heparinized saline before connecting the catheter to the syringe mounted on the syringe pump outside of the sound attenuating chamber via a 20 ga luer hub and 28 ga male connector. At the start of each self-administration session, the stimulus light above the active lever was illuminated and both the active and inactive levers were extended. A response on the active lever (FR1) resulted in a 20 sec time out (FR1:TO 20 sec) during which time the subject received a 200 µL intravenous drug infusion (over the first six seconds), lever light extinguished, levers are retracted, tone is generated, and the house light was illuminated. At the end of the TO, the levers were extended, lever light illuminated, tone silenced, and the house light extinguished. The self-administration session continued for 2 hours or until 70 infusions were delivered. Self-administration studies were initiated once stable responding was achieved (defined as total number of infusions ± <20% of the mean of the three previous sessions).[24–26]

**Effect of oxa-noribogaine on morphine self-administration**

Separate groups of rats were trained to self-administer morphine (10, 20 or 40 µg/infusion) under an FR1 schedule of reinforcement. After stable responding was established, rats were administered vehicle (2 mL/kg, i.p.) 30 min prior to the subsequent experimental session. Four days following vehicle administration, rats were administered oxa-noribogaine in an ascending dose manner (3, 10 or 30 mg/kg, i.p.), administered 30 min prior to the beginning of the session. Responding was assessed for seven days following each dose. For each morphine dose, baseline and vehicle active and inactive lever responding was compared using paired t-tests, two tailed. The acute effects of oxa-noribogaine on responding (active and inactive lever) maintained by each morphine dose was assessed independently using a two-way ANOVA with Tukey's multiple comparison test. The effects of oxa-noribogaine on morphine self-administration on active and inactive lever responding for the seven days following each dose of oxa-noribogaine were assessed using a mixed-model analysis of repeated measures data (due to missing data points) with Šidák's multiple comparison test for *post hoc* comparison.

**Effect of oxa-noribogaine on morphine self-administration under a progressive ratio schedule of reinforcement**

A separate cohort of rats were trained to self-administer morphine (20 µg/infusion) under FR1 for seven days and then the schedule was changed to progressive ratio schedule as previously reported by Grasing and colleagues.[27] The number of responses required to obtain each successive injection of morphine was calculated according to the following equation:

$$= \mathrm{Round}(10 * -e^{|0.035*(\mathrm{Step\ number}-1)|} - 10 + 0.5)$$

Results are rounded to the nearest integer value and the step number is the number of ratios completed.

After seven days of IVSA under the PR schedule, rats were administered vehicle (2 mL/kg, i.p.) 30 min prior to the subsequent experimental session. Four days following vehicle administration, rats were administered oxa-noribogaine in an ascending dose manner (3, 10 and 30 mg/kg, i.p.), administered 30 min prior to the beginning of the session. Responding was assessed for seven days following each dose. The "breakpoint" was defined as the cumulative number of responses on the active lever for the last completed ratio. The acute effects of each dose of oxa-noribogaine on active lever and inactive lever responding were analyzed using a one-way ANOVA with Tukey's post hoc analysis. The effects of oxa-noribogaine on morphine self-administration on active and inactive lever responding for the seven days following each dose of oxa-noribogaine as well as the number of infusions were assessed using a mixed-model analysis of repeated measures data (due to missing data points) with Šidák's multiple comparison test for *post hoc* comparison.

**Effect of oxa-noribogaine on cue-induced reinstatement of responding previously maintained by morphine or fentanyl**

Two groups of rats were trained to self-administer morphine (20 µg/infusion) or fentanyl (0.625 µg/infusion) in 2-h sessions under FR1 schedule of reinforcement. After a minimum of two weeks of self-administration and three days of stable responding immediately prior to the extinction session, rats underwent operant extinction training in 2 h session in the absence of the drugs and their associated cues until responding was < 20% of baseline responding maintained by intravenous self-administration of the drug. During the extinction phase, levers were available, but responding on the active lever did not result in cue light presentation or drug delivery.

The day following the last extinction session, each group was divided into two subgroups to receive either vehicle (2 mL/kg, i.p.) or oxa-noribogaine (10 mg/kg, i.p.) 30 min prior to the 60 min session to test for cue-induced reinstatement of morphine or fentanyl. During the assessment of cue-induced responding, cues previously associated with morphine or fentanyl self-administration were presented following each response on the previously active lever. Responses on the previously inactive lever were recorded but did not result in stimulus presentation.

Baseline responding (three days prior to extinction) on the active and inactive levers were compared separately for the morphine and fentanyl groups receiving either vehicle or oxa-noribogaine using a two-way ANOVA with time as the repeated measure with Šidák's multiple comparison test for *post hoc* comparison. For the cue-induced reinstatement test session, responding on the active and inactive levers were analyzed for the morphine and fentanyl groups separately receiving either vehicle or oxa-noribogaine using an unpaired t-test.

**Effect of acute administration of oxa-noribogaine, epi-oxa-noribogaine and noribogaine on morphine self-administration**

Rats were trained to self-administer morphine (10 µg/infusion) as described in the general self-administration training. After stable responding was established, rats were administered vehicle (2 mL/kg, i.p.) 15 min prior to the subsequent experimental session. Three days following vehicle administration, rats were administered oxa-noribogaine (10 or 40 mg/kg, i.p.), epi-oxa-noribogaine (40 mg/kg, i.p.) or noribogaine (40 mg/kg, i.p.), administered 15 min prior to the beginning of the session.

**Effect of acute oxa-noribogaine administration on fentanyl self-administration**

Rats were trained to self-administer fentanyl (625 ng/ infusion) as described in the general self-administration training. After stable responding was established, rats were administered vehicle 15 min prior to the subsequent experimental session. Three days following vehicle administration, rats were administered oxa-noribogaine (40 mg/kg, i.p.) administered 30 min prior to the beginning of the session.

**Food Maintained Responding**

For food maintained responding studies food access was restricted such that rats were maintained at 90% of their normal body weight until responding stabilized, at which point rats were provided 12-15 g of rodent chow in addition to the food pellets earned during the experimental session.

Rats were lever trained as described in the general methods. Responding was maintained as described with notable difference in timeout (TO) period 6-min and sessions lasting 2 hours or until a maximum of 20 pellets was delivered. Once responding stabilized, subjects were assigned randomly to receive oxa-noribogaine (10 and 40 mg/kg, i.p.), epi-oxa-noribogaine (40 mg/kg, i.p.), noribogaine (10 and 40 mg/kg, i.p.) or vehicle administered intraperitoneal 15 min prior to the initiation of the session. Responding was considered stable when variation in the number of reinforcers for three consecutive sessions was less than 20%. For the oxa-noribogaine repeated dosing procedure, all doses were administered 15 min prior to placing the rats in their respective operant chambers. Rats were administered oxa-noribogaine (40 mg/kg, i.p.) or vehicle on Day 1, followed by oxa-noribogaine (10 mg/kg, i.p.) on Days 6, 8 10 and 12. For the group receiving oxa-noribogaine, data were analyzed using a two-way ANOVA with Šidák's multiple comparisons test for *post hoc* comparison.

**Effect of repeated administration of oxa-noribogaine on morphine self-administration**

Rats were trained to self-administer morphine (10 µg/infusion; n = 11) as described in the general self-administration training. Following three days of stable responding, rats were administered vehicle 15 min prior to the subsequent experimental session. Five days following vehicle administration, a repeated administration procedure of oxa-noribogaine was initiated as follows: Days 1 and 6: 40 mg/kg, Days 11, 15, 17 and 19: 10 mg/kg, and Days 21, 23 and 25: 5 mg/kg. The respective doses of oxa-noribogaine were administered 15 min prior to the experimental sessions as indicated above.

**GDNF and mBDNF protein expression Studies**

Male Fisher F344 rats were assigned to groups and administered oxa-noribogaine (40 mg/kg; i.p.) or noribogaine (40 mg/kg; i.p.). One days or five days after drug administration, rats from these groups as well as non-treated controls were decapitated, brains removed and placed in a stainless-steel rat brain matrix. Coronal slices were taken and the ventral tegmental area (VTA), nucleus accumbens (NAc) and medial prefrontal cortex (mPFC) were dissected and immediately frozen on dry ice. Total protein was isolated from pulverized tissue from these regions. GDNF and mature BDNF were assayed using the BiosSensis GDNF, Rat, Rapid™ ELISA assay and the BDNF, mature, human, mouse, rat Rapid™ ELISA assay (Biosensis Pty Ltd, SA, Australia). Protease (Thermo Scientific, Rockford, IL), and phosphatase inhibitors (Cocktails 1 and 2, Sigma-Aldrich, St. Louis, MO) were added to the extraction buffers. Samples were sonicated twice for 15-20 seconds and incubated on ice for 30 seconds between sonication. Samples were then centrifuged at 15,000 rpm for 10 minutes at $4^oC$ and the supernatant (total protein lysate) was transferred to a new tube. Protein concentrations were measured using the bicinchoninic acid protein assay kit (Pierce, Rockford, IL, USA) on a spectrophotometer (iD5, Molecular Devices, Sunnyvale, CA). Aliquots of 100 µL of isolated protein from each region were transferred to a 96-well ELISA plates. The abundance of GDNF and mBDNF were normalized to the amount of total protein (pg/mg protein). In a separate cohort, we assessed the ability of the kappa antagonist aticaprant to attenuate the oxa-noribogaine induced increase in GDNF in the VTA and mPFC five days after administration. Rats were administered either vehicle (7% (v/v) Tween 80, 2 molar equivalents glacial acetic acid in physiologic saline) or aticaprant (1 mg/kg; s.c.) followed twenty minutes later by administration of either vehicle (2 molar equivalents glacial acetic acid in physiologic saline) or oxa-noribogaine (40 mg/kg; i.p.). Along with non-treated controls, rats were decapitated, and tissue processed as described above. Data were analyzed by protein and region using one way ANOVA with Tukey's multiple comparison test for *post hoc* comparison.

**Pharmacokinetic Studies (PK, Sai Life Sciences Limited)**

Total of twenty-four male mice (twenty-seven male rats) were used per study (3 animals per each time point). Mice were administered subcutaneously (10 mg/kg dose, s.c.), rats intraperitoneally (40 mg/kg dose, i.p.). The dosing volume for subcutaneous administration was 5 mL/kg and 10 mL/kg for intraperitoneal administration.

*Sample collection:* Blood samples (~60 µL from mice, ~120 from rats) were collected under light isoflurane anesthesia (Surgivet®) from retro orbital plexus from a set of three animals at specified time points into labeled micro-tubes, containing $K_2$EDTA solution (20% $K_2$EDTA solution) as an anticoagulant. Immediately after blood collection, plasma was harvested by centrifugation at 4000 rpm, 10 min at 40 °C and samples were stored at -70 ± 10 ºC until bioanalysis. Following blood collection, animals were immediately sacrificed, the abdominal vena-cava was cut open and whole body was perfused from heart using (10 mL for mice or 20 mL for rats) of normal saline. Brain samples were collected from a set of three animals at specified time points. After isolation, brain samples were rinsed three times in ice cold normal saline (for 5-10 seconds/rinse using (~5-10 mL for mice or ~10-20 mL) of normal saline in disposable petri dish for each rinse) and dried on blotting paper. Brain samples were homogenized using ice-cold phosphate buffer saline (pH - 7.4). Total homogenate volume was three times the tissue weight. All homogenates were stored below -70 ± 10 ºC until bioanalysis. The extraction procedure for plasma and brain samples and the spiked plasma and brain calibration standards were identical. A 25 µL of study sample or spiked plasma calibration standard was added to individual pre-labeled micro-centrifuge tubes followed by 100 µL of internal standard prepared in acetonitrile (Glipizide, 500 ng/mL) was added except for blank, where 100 µL of acetonitrile was added. Samples were vortexed for 5 minutes. Samples were centrifuged for 10 minutes at a speed of 4000 rpm at 4 °C. Following centrifugation, 100 µL of clear supernatant was transferred in 96 well plates and the concentrations of analyte were determined by fit for purpose LC-MS/MS method.

*Data analysis:* Non-Compartmental-Analysis tool of Phoenix WinNonlin® (Version 8.0 for oxa-noribogaine, Version 7.0 for noribogaine) was used to assess the pharmacokinetic parameters. Peak plasma concentration ($C_{max}$) and time for the peak plasma concentration ($T_{max}$) were the observed values. The areas under the concentration time curve ($AUC_{last}$ and $AUC_{inf}$) were calculated by linear trapezoidal rule. The terminal elimination rate constant, ke was determined by regression analysis of the linear terminal portion of the log plasma concentration-time curve. The terminal half-life ($T_{1/2z}$) was estimated by 0.693/ke. Clearance was estimated as Dose/$AUC_{inf}$ and $V_{ss}$ as CL × MRT. Tissue-Kps were calculated using Microsoft excel.

**Plasma Protein and Brain Tissue Binding (Sai Life Sciences Limited)**

Noribogaine, oxa- and epi-oxa-noribogaine plasma protein and brain tissue binding were determined using rapid equilibrium dialysis (RED) followed by LC-MS/MS quantification in MRM mode. Brain tissue homogenate samples were prepared by diluting one volume of whole brain tissue with three volumes of dialysis buffer (phosphate buffered saline pH 7.4 with 0.1 M sodium phosphate and 0.15 M sodium chloride) to yield 4 times diluted homogenate. A 1 mM stock solution of test compounds was prepared in DMSO and diluted 200-folds in rat plasma (male Sprague Dawley, n = 5), human plasma (drug free volunteers; n = 6) or rat brain homogenate to prepare a concentration of 5 µM. The final DMSO concentration was 0.5%. Rapid equilibrium dialysis was performed with RED device containing dialysis membrane with a molecular weight cut-off of 8,000 Daltons (Thermo Scientific). A 200 µL aliquot of positive control 5 µM and test compound at 5 µM (triplicates) were separately added to the plasma/brain homogenate chamber and 350 µL of phosphate buffer saline (pH 7.4, Thermo Scientific) was added to the buffer chamber of the inserts. After sealing the RED device with an adhesive film, dialysis was performed in incubator at 37 °C with shaking at 100 RPM for 4 hours.

*Recovery and stability.* A 50 µL aliquot of positive control and test compounds were added to four 0.5 mL microcentrifuge tubes. Two aliquots were frozen immediately (0 minute sample). The other two aliquots were incubated at 37 °C for 4 hours along with the RED device.

Following dialysis, an aliquot of 50 µL was removed from each well (both plasma or brain homogenate and buffer side) and diluted with equal volume of opposite matrix (dialyzed with the other matrix) to nullify the matrix effect. Similarly, 50 µL of buffer was added to recovery and stability samples. An aliquot of 100 µL was submitted for LC-MS/MS analysis.

*Sample Preparation and Bio-analysis.* A 25 µL aliquot of positive control and test compounds were crashed with 100 µL of acetonitrile containing internal standard and vortexed for 5 minutes. The samples were centrifuged at 4000 RPM at 4 °C for 10 min and 100 µL of supernatant was submitted for LC-MS/MS analysis in MRM mode. The samples were run without calibration curve and the peak area ratios (analyte versus internal standard) obtained was used to determine the fraction of compound bound to plasma and brain proteins.

**Estimated free drug concentrations, target engagement, and correlation with in vitro pharmacology**

Oxa-noribogaine exhibits rapid and high brain penetration (brain/plasma = 5.5, mice) after subcutaneous administration (s.c.). Approximately 85% of the compound is cleared in the first 2 hours and only traces remain after 8 hours. We estimated the free plasma and brain drug concentrations based on plasma protein and brain tissue binding ($C_{max(brain)}$ = 166 nM, 10 mg/kg, s.c.). Assuming linearity of PK below 10 mg/kg, $C_{max(brain)}$ at an $ED_{50}$ analgesic dose (3 mg/kg) is expected to be ~ 50 nM, which matches well the in vitro KOR activation potency ($EC_{50}$ = 41 nM in BRET assay, $EC_{50}$ = 49 nM, in [$^{35}$S]GTPgS assay). Thus the pharmacodynamic readout (analgesia) correlates with the estimated free drug PK measures. Further, the $C_{max}$ and area under the curve (AUC, drug exposure) values for estimated free drugs are much smaller for oxa-noribogaine in comparison to noribogaine, which is relevant for interpreting the oxa-noribogaine's superior efficacy in addiction models.

### Ibogaine Analogs Docking Studies on KOR

The active state KOR X-ray crystal structures corresponding to PDB accession code 6B73[28] was extracted from the RCSB server. Three conserved crystallographic waters close of $H^{6.52}$ and $Y^{3.33}$, elucidated in the high-resolution active state MOR X-ray crystal structure (PDB 5C1M)[29] were transplanted to the active state KOR model and their steric positions inside the active state KOR model were optimized by minimization protocols. All the objects except the receptor protein subunit, the crystallized ligand, and the three transplanted waters were deleted, and this was followed by the addition of hydrogens and optimization of the side-chain residues. Ligands were sketched, assigned formal charges, and energy-optimized prior to molecular docking. The ligand docking box for potential grid docking was defined to contain the extracellular half of the protein, and all-atom docking was performed using the energy minimized structures for all ligands with an effort value of 10. The best-scored docking solutions were further optimized by iterative rounds of minimization and Monte Carlo sampling of the ligand conformation, including the surrounding side-chain residues (within 5 Å of the ligand) and the three water molecules in the KOR orthosteric site. All the above-mentioned molecular modeling operations were performed in ICM-Pro v3.9 molecular modeling and drug discovery suite (Molsoft LLC).[30]

### Synthesis

Reagents and solvents (including anhydrous solvents) were obtained from commercial sources and used without further purification unless otherwise stated. All compounds apart from noribogaine were prepared in racemic form. All reactions were performed in oven- or flame-dried glassware under an argon atmosphere unless otherwise stated and monitored by TLC using solvent mixtures appropriate to each reaction. All column chromatography was performed on silica gel (40-63 μm). Preparative TLC was performed on 20 x 20 cm plates with a 1 mm silica layer. For compounds containing a basic nitrogen, triethylamine ($Et_3N$) was often used in the mobile phase to provide better resolution. In these cases, TLC plates were pre-soaked in the $Et_3N$ containing solvent and then briefly dried before use, to yield an accurate Rf value. Nuclear magnetic resonance spectra were recorded on Bruker 400 or 500 MHz instruments as indicated. Chemical shifts are reported as δ values in ppm referenced to $CDCl_3$ ($^1$H NMR = 7.26 and $^{13}$C NMR = 77.16). Multiplicity is indicated as follows: s (singlet); d (doublet); t (triplet); q (quartet); p (pentet); dd (doublet of doublets); ddd (doublet of doublet of doublets); dt (doublet of triplets); td (triplet of doublets); m (multiplet); br (broad). For compounds containing a carbamate group, complex spectra with split peaks were observed (presence of conformers). As a result of these effects, multiple peaks may correspond to the same proton group or carbon atom. When possible, this is indicated by an "and" joining two peaks or spectral regions. Alternatively, when certain carbon peaks overlap and thus represent two carbons they were indicated by (2C) designation. For easier identification of carbamate containing compounds on the included $^1$H NMR spectra, groups of peaks arising from the hindered rotation were integrated together to obtain whole number integrals. All carbon peaks are rounded to one decimal place. Low-resolution mass spectra were recorded on either a JEOL LCmate (ionization mode: APCI+) or Advion expression-L CMS quadrupole mass spectrometer equipped with ESI and APCI sources. High-resolution mass spectra (HRMS) were acquired on a high-resolution Waters XEVO G2-XS QToF mass spectrometer equipped with a UPC2 SFC inlet, on-board fluidics and an ESI probe.

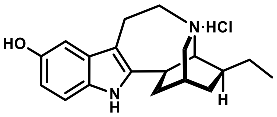


Noribogaine hydrochloride

##### Noribogaine hydrochloride

The compound was synthetized as previously described starting from Voacangine isolated from root bark of *Voacanga africana*.[31]

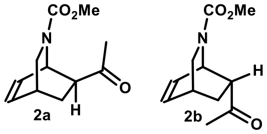


##### Synthesis of *exo/endo*-isoquinuclidine ketone intermediates 2a and 2b

The isoquinuclidine core was synthesized according to the reported literature procedures.[32,33] Briefly, methyl pyridine-1(2*H*)-carboxylate **1** was prepared by treating a mixture of pyridine and sodium borohydride in methanol with methyl chloroformate at -70°C. The crude diene (unstable, slowly decomposing during storage, should be used for next step as soon as possible) was filtered through silica (eluted with 10% diethyl ether in hexanes), concentrated and then heated with methyl vinyl ketone at 50 °C for 5 days to form a mixture of *exo* **2a** and *endo* **2b** compounds in a 26:74 ratio. The mixture was further epimerized using sodium methoxide in methanol to obtain a new ratio of *exo:endo* 62:38.

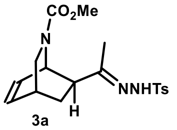
3a

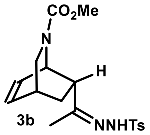
3b

**Synthesis of *exo/endo*-isoquinuclidine tosylhydrazones 3a and 3b**

A mixture of exo/endo-intermediates **2a/2b** (62:38 *exo*:*endo* ratio, 34.93 g, 167 mmol) and *p*-toluenesulfonyl hydrazide (31.10 g, 167 mmol) in THF (136 mL) was heated at 50 °C for 15 h, at which time a white precipitate had formed. The reaction mixture was cooled to room temperature and the white precipitate was collected by filtration, the solid was washed 3× on the filter with ice-cold MeOH, to provide pure *endo*-tosylhydrazone **3b** as a fine white powder (19.55 g, 31%). The filtrate and washings were combined and concentrated to a tan solid, which was recrystallized from MeOH to obtain the pure *exo*-tosylhydrazone **3a** as white plates (33.14 g, 53%).

***exo*-Methyl 7-(1-(2-tosylhydrazono)ethyl)-2-azabicyclo[2.2.2]oct-5-ene-2-carboxylate (3a)**

Mp 162-166 °C (inaccurate due to trapped MeOH); **$^{1}$H NMR (400 MHz, $CDCl_3$)** (spectrum complicated by conformers) δ 7.85 and 7.79 (d, *J* = 8.3 Hz, 2H), 7.35 – 7.28 (m, 3H), 6.46 – 6.37 (m, 2H), 4.74 – 4.68 and 4.61 – 4.57 (m, 1H), 3.52 and 3.34 (s, 3H), 3.02 and 2.94 (dd, *J* = 9.8, 2.2 Hz, 1H), 2.87 and 2.78 (dt, *J* = 9.8, 2.6 Hz, 1H), 2.73 – 2.67 (m, 1H), 2.48 – 2.37 (m, 1H), 2.44 (s, 3H), 2.30 and 2.09 (ddd, *J* = 13.1, 4.4, 2.4 Hz, 1H), 1.89 and 1.80 (s, 3H), 1.42 – 1.26 (m, 1H); **LR-MS** calcd. for $C_{18}H_{24}N_3O_4S^+$ $[M+H]^+$ 378.15, found 377.80.

***endo*-Methyl 7-(1-(2-tosylhydrazono)ethyl)-2-azabicyclo[2.2.2]oct-5-ene-2-carboxylate (3b)**

Mp 181-184 °C; **$^{1}$H NMR (400 MHz, $CDCl_3$)** (spectrum complicated by conformers) δ 7.81 (t, *J* = 7.4 Hz, 2H), 7.32 (d, *J* = 8.0 Hz, 2H), 7.12 (s, 1H), 6.19 (q, *J* = 7.5 Hz, 1H), 6.02 (dd, *J* = 12.0, 5.6 Hz, 1H), 4.87 and 4.77 (d, *J* = 4.3 Hz, 1H), 3.69 and 3.66 (s, 3H), 3.23 (d, *J* = 10.1 Hz, 1H), 3.01 – 2.85 (m, 2H), 2.74 (br s, 1H), 2.45 and 2.44 (s, 3H), 1.87 – 1.66 (m, 1H), 1.71 and 1.69 (s, 3H), 1.59 – 1.47 (m, 1H); **LR-MS** calcd. for $C_{18}H_{24}N_3O_4S^+$ $[M+H]^+$ 378.15, found 377.80.

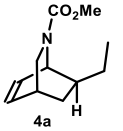
4a

***exo*-Methyl 7-ethyl-2-azabicyclo[2.2.2]oct-5-ene-2-carboxylate (4a)**

*exo*-Tosylhydrazone **3a** (33.02 g, 87.5 mmol), sodium cyanoborohydride (21.99 g, 350 mmol), and *p*-toluenesulfonic acid monohydrate (1.40 g, 7.36 mmol) were combined in THF (250 mL) and refluxed for 21 h. At this time, additional *p*-TsOH·$H_2O$ (0.35 g, 1.84 mmol) was added, and reflux was continued for an additional 4 h. The reaction mixture was then diluted with water (250 mL) and extracted with cyclohexane (3 × 100 mL). The combined organics were washed with water (250 mL), saturated aqueous $NaHCO_3$ (250 mL), and water again (50 mL), dried over $Na_2SO_4$, and concentrated to provide a cloudy, pale-yellow oil. The oil was passed through a short silica column in 7:3 hexanes:EtOAc and the eluate was concentrated to yield pure product as a pale-yellow oil (10.12 g, 59%). The spectral characterization was in agreement with the previously reported literature data.[34]

**$^{1}$H NMR (400 MHz, $CDCl_3$)** δ 6.47 and 6.42 (br dd, *J* = 7 Hz, 1H), 6.34 and 6.30 (br dd, *J* = 8 Hz, 1H), 4.59 and 4.44 (d, *J* = 6 Hz, 1H), 3.67 and 3.66 (s, 3H), 3.20 (td, *J* = 10, 2 Hz, 1H), 2.98 and 2.94 (dt, *J* = 10, 3 Hz, 1H), 2.65 (m, 1H), 1.64 and 1.61 (dt, *J* = 10, 3 Hz, 1H), 1.38 (m, 3H), 1.00 (m, 1H), 0.95 and 0.92 (t, *J* = 7 Hz, 3H); **$^{13}$C NMR (101 MHz, $CDCl_3$)** δ 156.7 and 156.3, 133.8 and 133.7, 133.3 and 133.1, 52.2 and 52.1, 49.1 and 48.8, 48.4 and 48.1, 40.7, 30.8 and 30.6, 29.9, 27.5 and 27.4, 12.1 and 12.0; **LR-MS** calcd. for $C_{11}H_{18}NO_2^+$ $[M+H]^+$ 196.13, found 195.9.

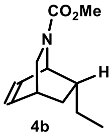

4b

**_endo_-Methyl 7-ethyl-2-azabicyclo[2.2.2]oct-5-ene-2-carboxylate (4b)**

To a suspension of endo-tosylhydrazone **3b** (3.50 g, 9.27 mmol) in MeOH (41 mL) was added a solution of sodium cyanoborohydride (0.83 g, 13.26 mmol) and $ZnCl_2$ (0.90 mg, 6.63 mmol) in MeOH (28 mL) and the resulting mixture was refluxed for 3 h. The reaction was then quenched with 1% aqueous NaOH (200 mL) and extracted with cyclohexane (3 × 50 mL). The combined organics were washed with water (50 mL) and brine (50 mL), dried over $Na_2SO_4$, and concentrated to provide a clear, colorless oil. The crude material was filtered through a short silica column with 1:1 hexanes:EtOAc and the eluate was concentrated to yield pure product as a colorless oil (1.11 g, 61%).

**$^1$H NMR (400 MHz, $CDCl_3$)** δ 6.31 (m, 2H), 4.66 and 4.48 (s, 1H), 3.69 and 3.66 (s, 3H), 3.21 (m, 1H), 2.94 (m, 1H), 2.69 (m, 1H), 1.96 (m, 1H), 1.81 (m, 1H), 1.19 (m, 1H), 1.01 - 0.84 (m, 5H); **$^{13}$C NMR (101 MHz, $CDCl_3$)** δ 155.6 and 155.2, 134.4 and 134.0, 130.6 and 130.0, 52.0 and 51.9, 49.3 and 48.9, 46.9 and 46.5, 40.7 and 40.5, 31.0 and 30.8, 30.0, 28.3 and 28.2, 11.2; **LR-MS** calcd. for $C_{11}H_{18}NO_2^+$ $[M+H]^+$ 196.13, found 195.90.

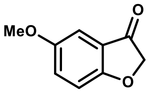

5

**5-Methoxybenzofuran-3(2H)-one (5)**

Benzofuranone **5** was prepared from 4-methoxyphenol according to literature procedure[35] and obtained as orange-tan crystals (yield 30 – 40%).

**$^1$H NMR (500 MHz, $CDCl_3$)** δ 7.24 (dd, $J$ = 9.0, 2.8 Hz, 1H), 7.06 (d, $J$ = 5.8 Hz, 1H), 7.05 (s, 1H), 4.64 (s, 2H), 3.80 (s, 3H); **$^{13}$C NMR (126 MHz, $CDCl_3$)** δ 200.3, 169.5, 155.2, 128.1, 121.2, 114.7, 104.0, 75.6, 56.1; **LR-MS** calcd. for $C_9H_9O_3^+$ $[M+H]^+$ 165.06, found 164.99.

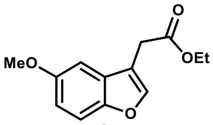

6a

**Ethyl 2-(5-methoxybenzofuran-3-yl)acetate (6a)**

A solution of benzofuranone **5** (4.92 g, 30.00 mmol) and (carbethoxymethylene)triphenylphosphorane (11.50 g, 33.00 mmol) in toluene (100 mL) was refluxed for 110 h and then concentrated in vacuo. The resulting material was triturated with 9:1 hexanes:EtOAc (210 mL) and filtered, and the remaining solids were washed with additional portions of 9:1 hexanes:EtOAc (3 × 90 mL). The combined filtrates were concentrated to give the crude product, which was purified by column chromatography (1:1 hexanes:$CH_2Cl_2$, 5 column volumes → $CH_2Cl_2$, 3 column volumes → 1:1 $CH_2Cl_2$:$Et_2O$, 1 column volume). Collected fractions were concentrated to provide product **6** as a thin, yellow-orange oil (5.68 g, 81%). The spectral characterization was in agreement with the previously reported literature data.[36]

**$^1$H NMR (400 MHz, $CDCl_3$)** δ 7.60 (s, 1H), 7.36 (d, $J$ = 9.0 Hz, 1H), 7.02 (d, $J$ = 2.6 Hz, 1H), 6.91 (dd, $J$ = 8.9, 2.6 Hz, 1H), 4.19 (q, $J$ = 7.1 Hz, 2H), 3.85 (s, 3H), 3.66 (d, $J$ = 1.0 Hz, 2H), 1.28 (t, $J$ = 7.1 Hz, 3H); **$^{13}$C NMR (101 MHz, $CDCl_3$)** δ 170.8, 156.1, 150.4, 143.8, 128.3, 113.42, 113.39, 112.1, 102.3, 61.2, 56.1, 30.1, 14.4; **LR-MS** calcd. for $C_{13}H_{15}O_4^+$ $[M+H]^+$ 235.10, found 235.37.

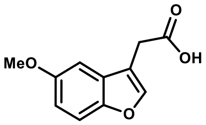

6b

**2-(5-methoxybenzofuran-3-yl)acetic acid (6b)**

Compound **6b** was prepared using slightly modified published reaction conditions.[37] A mixture of 4-methoxyphenol (24.8 g, 200 mmol) and ethyl 4-chloroacetoacetate (32.64 mL, 240.00 mmol) was cooled in

an ice bath and cold (~ 4 °C) 70% aqueous sulfuric acid was added (1 mL of acid per 1 mmol of phenol). Reaction mixture was further stirred for 1-3 days and quenched with ice and ice-cold water (final volume ~500 mL). The resulting beige suspension was further stirred for 1 h and was extracted with $CH_2Cl_2$ (3 × 250 mL). Combined dark brown extracts were dried over $Na_2SO_4$ and filtered through a plug of silica, washing the plug with more $CH_2Cl_2$, until all product eluted (solvent was recycled by evaporation and reused for further elution). Collected fractions were concentrated to yield the coumarin intermediate as a bright yellow solid (35.6 g). The crude intermediate was then suspended in 1M NaOH (500 mL) and heated to reflux for 2 h. After cooling to room temperature pH of reaction mixture was adjusted to ~6 using concentrated (36-38%) aq. hydrochloric acid (~20 mL) and was back adjusted to 7 using addition of solid $NaHCO_3$. Neutral aqueous solution was washed with diethyl ether (2 × 150 mL, removal of unreacted phenol). Aqueous mixture was further acidified to pH ~2 using concentrated (36-38%) aq. hydrochloric acid and the formed suspension was extracted with $CH_2Cl_2$ (4 × 250 mL, product dissolves slowly initially). Combined extracts were dried over $Na_2SO_4$, filtered and concentrated to yield the crude acid **6b** (~90-95% pure) as a beige solid (25.00 g, 58%). The crude material was used as is for the next step. The spectral characterization was in agreement with the previously reported literature data.[37]
**$^1$H NMR (400 MHz, CDCl$_3$)** δ 10.73 (s, 1H), 7.61 (d, *J* = 1.1 Hz, 1H), 7.37 (d, *J* = 8.9 Hz, 1H), 7.00 (d, *J* = 2.6 Hz, 1H), 6.92 (dd, *J* = 8.9, 2.6 Hz, 1H), 3.85 (s, 3H), 3.73 (d, *J* = 1.1 Hz, 2H). **$^1$H NMR (500 MHz, DMSO)** δ 12.42 (s, 1H), 7.84 (d, *J* = 1.2 Hz, 1H), 7.45 (d, *J* = 8.9 Hz, 1H), 7.11 (d, *J* = 2.6 Hz, 1H), 6.90 (dd, *J* = 8.9, 2.6 Hz, 1H), 3.78 (s, 3H), 3.67 (d, *J* = 1.1 Hz, 2H); **LR-MS** calcd. for $C_{11}H_{10}O_4^-$ [M-H]$^-$ 205.1, found 205.4.

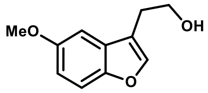

7

**2-(5-Methoxybenzofuran-3-yl)ethanol (7)**
From **6a**: To a suspension of $LiAlH_4$ (2.36 g, 62.09 mmol) in THF (60 mL) at room temperature was carefully added a solution of the ester **6a** (5.60 g, 23.88 mmol) in THF (20 mL), and the mixture was refluxed for 30 min. After cooling to room temperature, the reaction was quenched by the successive addition of $H_2O$ (2.4 mL), 15% aqueous NaOH (2.4 mL), and $H_2O$ again (7.2 mL). The resulting mixture was stirred vigorously until the aluminum salts were white and loose and then filtered, washing the filter cake with $Et_2O$ (3 × 60). The combined filtrate and washings were concentrated to yield the product **7** directly as a yellow oil (4.50 g, 98%). The spectral characterization was in agreement with the previously reported literature data.[38]
From **6b**: Suspension of $LiAlH_4$ (2.85 g, 75.0 mmol) in THF (30.0 mL) was cooled using water/ice-bath and the solution of crude (~90-95% pure) acid **6b** (6.51 g, ~30.00 mmol) in THF (30.0 mL) was added in small portions via canula. Reaction mixture was allowed to warm to room temperature and stirred until no more starting material was detected by TLC (<1 h). The reaction mixture was again cooled using water/ice-bath before adding diethyl ether (not anhydrous, 60 mL) and carefully quenching it by the successive addition of $H_2O$ (2.9 mL), 15% aqueous NaOH (2.9 mL), and $H_2O$ again (8.7 mL). The resulting mixture was stirred vigorously until the aluminum salts were pale and loose, suspension was dried by addition of $MgSO_4$ and filtered, washing the filter cake with diethyl ether until no more product eluted. The combined filtrate and washings were concentrated to yield the product **7** directly as a yellow oil (5.23 g, 90%).
**$^1$H NMR (500 MHz, CDCl$_3$)** δ 7.48 (s, 1H), 7.36 (d, *J* = 8.9 Hz, 1H), 7.00 (d, *J* = 2.6 Hz, 1H), 6.90 (dd, *J* = 8.9, 2.6 Hz, 1H), 3.91 (t, *J* = 6.4 Hz, 2H), 3.85 (s, 3H), 2.91 (t, *J* = 6.4 Hz, 2H), 1.74 (s, 1H); **$^{13}$C NMR (126 MHz, CDCl$_3$)** δ 156.0, 150.5, 143.2, 128.7, 117.0, 113.2, 112.1, 102.3, 61.9, 56.1, 27.2; **LR-MS** calcd. for $C_{11}H_{13}O_3^+$ [M+H]$^+$ 193.09, found 193.35.

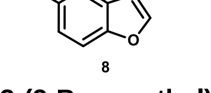

8

**3-(2-Bromoethyl)-5-methoxybenzofuran (8)**
To a solution of the alcohol **7** (2.18 g, 11.34 mmol) and carbon tetrabromide (5.64 g, 17.01 mmol) in $CH_2Cl_2$ (anhydrous, 23 mL) at room temperature was carefully added triphenylphosphine (4.46 g, 17.01 mmol) and the resulting dark orange-brown mixture was left to stir for 20 min. The reaction mixture was filtered through a silica plug to remove baseline impurities, washing the plug with additional $CH_2Cl_2$ until TLC indicated that all product was eluted. The filtrate was then concentrated and purified by column

chromatography (hexanes, 2 column volumes → 20:1 hexanes:$Et_2O$, 2 column volumes → 10:1 hexanes:$Et_2O$, 2 column volumes) to provide a pale-yellow oil that slowly crystallized to a white solid (2.81 g, 97%). The spectral characterization was in agreement with the previously reported literature data.[39]
**$^1$H NMR (500 MHz, $CDCl_3$)** δ 7.51 (s, 1H), 7.37 (d, *J* = 8.9 Hz, 1H), 6.97 (d, *J* = 2.5 Hz, 1H), 6.91 (dd, *J* = 8.9, 2.6 Hz, 1H), 3.86 (s, 3H), 3.64 (t, *J* = 7.4 Hz, 2H), 3.23 (td, *J* = 7.4, 0.7 Hz, 2H); **$^{13}$C NMR (126 MHz, $CDCl_3$)** δ 156.1, 150.4, 143.1, 128.1, 117.8, 113.2, 112.3, 101.9, 56.2, 31.3, 27.7; **LR-MS** calcd. for $C_{11}H_{12}BrO_2^+$ [M+H]$^+$ 255.00 and 257.00, found 255.29 and 257.29.

**General Procedure for Preparation of *N*-benzofuranylethylisoquinuclidines**
To a solution of a carbamate protected isoquinuclidine **4** (1 equivalent) in $CH_2Cl_2$ (anhydrous, 0.125 M, based on 2) at 0 °C was added iodotrimethylsilane (4 equivalents), and the resulting mixture was stirred for 10 min at 0 °C and then at room temperature until TLC indicated that no **4** remained (typically ~1 h). The reaction mixture was then quenched with MeOH (3.0 mL per mmol of **4**) and concentrated to yield the deprotected isoquinuclidine hydroiodide salt in quantitative yield. To this material was added 3-(2-bromoethyl)-5-methoxybenzofuran **8** (1 equivalent) and $NaHCO_3$ (4 equivalents), followed by anhydrous $CH_3CN$ (0.208 M, based on **4**), and the resulting mixture was refluxed until TLC indicated the disappearance of the bromide (typically >24 h). The reaction was then diluted with water, made strongly basic with aqueous NaOH, and extracted with $CH_2Cl_2$ (3×). The combined organics were washed with water, dried over $Na_2SO_4$, and concentrated to provide the crude product, which was purified by column chromatography with an appropriate solvent mixture (as described below for each compound).

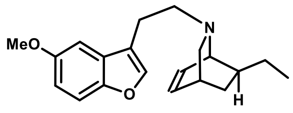

**9a**

***exo*-7-Ethyl-2-(2-(5-methoxybenzofuran-3-yl)ethyl)-2-azabicyclo[2.2.2]oct-5-ene (9a)**
The product **9a** was prepared according to the general procedure and purified by column chromatography (20:1 hexanes:$Et_2O$, 3 column volumes → 20:1 hexanes:$Et_2O$ + 2% $Et_3N$, 5 column volumes) to provide a pale-yellow oil (288 mg, 74%).
*Note (multigram scale preparation) reaction was scaled up to 30 mmol of starting materials **8** and **4a** to yield **9a** (7.02g, 75%).*
**$^1$H NMR (400 MHz, $CDCl_3$)** δ 7.46 (s, 1H), 7.33 (d, *J* = 8.9 Hz, 1H), 6.98 (d, *J* = 2.6 Hz, 1H), 6.87 (dd, *J* = 8.9, 2.6 Hz, 1H), 6.39 – 6.27 (m, 2H), 3.86 (s, 3H), 3.23 (dt, *J* = 5.3, 1.9 Hz, 1H), 3.09 (dd, *J* = 9.1, 2.3 Hz, 1H), 2.84 – 2.63 (m, 3H), 2.57 – 2.48 (m, 1H), 2.48 – 2.40 (m, 1H), 1.94 (dt, *J* = 9.1, 2.6 Hz, 1H), 1.63 – 1.42 (m, 3H), 1.35 – 1.24 (m, 1H), 0.95 – 0.90 (m, 1H), 0.88 (t, *J* = 7.4 Hz, 3H); **$^{13}$C NMR (101 MHz, $CDCl_3$)** δ 155.8, 150.3, 142.6, 133.1, 132.8, 129.2, 119.2, 112.6, 111.8, 102.5, 57.8, 56.3, 56.3, 56.2, 41.3, 31.8, 29.9, 27.4, 23.1, 12.6; **HR-MS** calcd. for $C_{20}H_{26}NO_2^+$ [M+H]$^+$ 312.1964, found 312.1948.

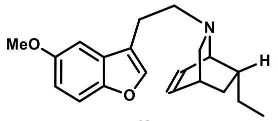

**9b**

***endo*-7-Ethyl-2-(2-(5-methoxybenzofuran-3-yl)ethyl)-2-azabicyclo[2.2.2]oct-5-ene (9b)**
The product **9b** was prepared according to the general procedure and purified by column chromatography (gradient of 5, 10 to 15% of EtOAc in hexanes + 2% $Et_3N$) to provide a pale-yellow oil (582 mg, 75%).
**$^1$H NMR (500 MHz, $CDCl_3$)** δ 7.42 (s, 1H), 7.33 (d, *J* = 8.9 Hz, 1H), 7.00 (d, *J* = 2.6 Hz, 1H), 6.88 (dd, *J* = 8.8, 2.6 Hz, 1H), 6.42 – 6.35 (m, 1H), 6.17 – 6.10 (m, 1H), 3.85 (s, 3H), 3.41 – 3.34 (m, 1H), 3.02 (dd, *J* = 9.7, 2.0 Hz, 1H), 2.88 – 2.70 (m, 3H), 2.57 – 2.48 (m, 2H), 2.07 (dt, *J* = 9.6, 2.7 Hz, 1H), 2.05 – 1.98 (m, 1H), 1.78 (ddd, *J* = 12.2, 9.2, 2.9 Hz, 1H), 1.22 – 1.13 (m, 1H), 1.05 – 0.95 (m, 1H), 0.85 (t, *J* = 7.3 Hz, 3H), 0.79 (ddt, *J* = 12.2, 5.1, 2.8 Hz, 1H); **$^{13}$C NMR (126 MHz, $CDCl_3$)** δ 155.8, 150.3, 142.4, 133.7, 130.2, 129.0, 118.9, 112.7, 111.9, 102.4, 57.8, 57.4, 56.1, 54.5, 40.9, 31.6, 30.7, 28.8, 23.2, 11.8; **HR-MS** calcd. for $C_{20}H_{26}NO_2^+$ [M+H]$^+$ 312.1964, found 312.1955.

**General Procedure for Preparation of Oxa-ibogaine Analogs by Ni(0)-catalyzed Cyclization**

In a glovebox, a vial was charged with Ni(COD)$_2$ (0.20 equivalents) and 1,3- bis(2,4,6-trimethylphenyl)-1,3-dihydro-*2H*-imidazol-2-ylidene (IMes, 0.24 equivalents) followed by heptane (0.100 M, based on Ni(COD)$_2$), and the resulting black solution was stirred at room temperature for 15 min. To this mixture was then added a solution of the substrate **9** (1 equivalent) in heptane (0.333 M, based on **9**), and the reaction vessel was sealed with a Teflon lined solid cap, removed from the glovebox, and heated at 130 °C for 3 h. After cooling to room temperature, the reaction mixture was purified as described below for each compound.

*Note, especially for larger scale using nonane/decane as solvent is preferred to avoid over pressurization of reaction vessel.*

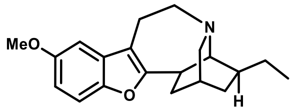


10a

**Oxa-ibogaine (10a)**

Prepared according to the general procedure. The crude reaction mixture was purified directly by column chromatography (30:1 hexanes:EtOAc + 1% $Et_3N$) to yield the crude product as a pale-yellow oil. This material was further purified by preparative TLC (30:1 hexanes:EtOAc + 1% $Et_3N$) to provide the pure product **10a** as a pale-brown oil (118 mg, 76%).

**$^1$H NMR (500 MHz, CDCl$_3$)** δ 7.24 (d, *J* = 8.7 Hz, 1H), 6.86 (d, *J* = 2.6 Hz, 1H), 6.81 (dd, *J* = 8.7, 2.6 Hz, 1H), 3.85 (s, 3H), 3.45 – 3.36 (m, 1H), 3.26 – 3.11 (m, 3H), 3.02 – 2.91 (m, 2H), 2.82 – 2.78 (m, 1H), 2.53 – 2.44 (m, 1H), 2.08 – 2.00 (m, 1H), 1.88 – 1.76 (m, 2H), 1.67 – 1.61 (m, 1H), 1.59 – 1.42 (m, 3H), 1.24 – 1.15 (m, 1H), 0.91 (t, *J* = 7.1 Hz, 3H); **$^{13}$C NMR (126 MHz, CDCl$_3$)** δ 161.1, 155.8, 148.6, 131.4, 111.8, 111.4, 111.0, 101.9, 57.2, 56.2, 53.3, 49.7, 41.5, 41.2, 33.1, 32.3, 27.5, 26.5, 19.5, 11.9; **HR-MS** calcd. for $C_{20}H_{26}NO_2^+$ [M+H]$^+$ 312.1964, found 312.1956.

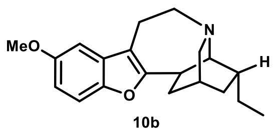


10b

**Epi-oxa-ibogaine (10b)**

Prepared according to the general procedure. The crude reaction mixture was purified directly by column chromatography (9:1 hexanes:EtOAc + 2% $Et_3N$, 4 column volumes → 8:2 hexanes:EtOAc + 2% $Et_3N$, 3 column volumes) to yield the crude product as a yellow-orange oil. This material was further purified by preparative TLC ($Et_2O$ + 1% $Et_3N$) to provide the pure product **10b** as a nearly colorless oil that slowly crystallized to a white solid (27.0 mg, 28%). **$^1$H NMR (400 MHz, CDCl$_3$)** δ 7.24 (d, *J* = 8.8 Hz, 1H), 6.87 (d, *J* = 2.6 Hz, 1H), 6.80 (dd, *J* = 8.8, 2.6 Hz, 1H), 3.85 (s, 3H), 3.43 (ddd, *J* = 13.7, 4.7, 2.3 Hz, 1H), 3.38 – 3.15 (m, 3H), 3.06 (qt, *J* = 9.5, 2.5 Hz, 2H), 2.86 (t, *J* = 2.3 Hz, 1H), 2.46 (dt, *J* = 16.3, 3.1 Hz, 1H), 2.09 – 1.90 (m, 3H), 1.92 – 1.84 (m, 1H), 1.66 – 1.56 (m, 1H), 1.45 – 1.31 (m, 2H), 1.18 – 1.05 (m, 1H), 0.93 (t, *J* = 7.3 Hz, 3H); **$^{13}$C NMR (101 MHz, CDCl$_3$)** δ 161.7, 155.9, 148.4, 131.3, 112.3, 111.5, 111.0, 101.8, 56.5, 56.2, 53.5, 49.1, 42.0, 34.5, 34.1, 31.7, 28.5, 26.4, 19.0, 12.3; **HR-MS** calcd. for $C_{20}H_{26}NO_2^+$ [M+H]$^+$ 312.1964, found 312.1983.

**General Procedure for Preparation of Oxa-noribogaine Analogs by Demethylation**

To a solution of the oxa-ibogaine **10** (1 equivalent) in $CH_2Cl_2$ (0.125 M, based on **10**) at 0 °C was added aluminum chloride (6 equivalents) followed by ethanethiol (18 equivalents), and the resulting mixture was allowed to warm to room temperature and stirred until TLC indicated the complete consumption of starting material (typically <1.5 h). The reaction was then quenched with saturated aqueous $NaHCO_3$ (100 mL per mmol of **10**) and extracted with $CH_2Cl_2$ (4 × to 6 ×, until no further extraction detected by TLC). The combined organic layers were dried over $Na_2SO_4$ and concentrated to provide the crude product. This material was purified by column chromatography with an appropriate solvent mixture (as described below for each compound).

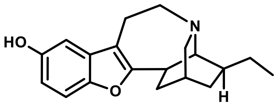

**11a**
**Oxa-noribogaine**

**_rac_-oxa-noribogaine (Oxa-noriboga, 11a)**
The product **11a** was prepared according to the general procedure and purified by column chromatography (1:1 hexanes:EtOAc) to provide a white, foamy solid (24.3 mg, 82%).

*Note (multigram scale preparation starting from **9a**):* In glovebox Ni(COD)$_2$ (0.94 g, 3.42 mmol) and IMes (1.25 g, 4.10 mmol) were balanced into three 40 mL oven-dried scintillation vials (divided in equal portions), sealed with cap with teflon septa and electrical (vinyl) tape and removed from glove box. Decane (11.4 mL per vial) was added into each vial and the black mixture was vigorously stirred for ~15 min (sonication was used to improve initial dissolution of components). Starting material **9** (5.32 g, 17.08 mmol) was dissolved in decane under Argon and divided equally among the reaction vials washing the original container 2 times with fresh solvent (total volume 51.6 mL decane, solution divided equally per vial). Dark mixture was heated to 130°C (135-140 °C vial heating block) and stirred vigorously. After 3 h mixture was cooled to room temperature and directly purified by repeated column chromatography (30:1 hexanes:EtOAc + 1% Et$_3$N). After first column crude material was dissolved in hot hexanes, colored insoluble impurities were filtered off and washed with hexanes. After second column chromatography slightly impure product (5.32 g) was dissolved in CH$_2$Cl$_2$ (71.2 mL) and cooled in ice-water bath. Aluminum chloride (6.83 g, 51.24 mmol) followed by ethanethiol (11.8 mL, 153.72 mmol) were added at 0 °C and the resulting mixture was allowed to warm to room temperature and stirred. After 3 h, reaction mixture was poured into a mixture of saturated solution of sodium bicarbonate (150 mL) and solution of potassium sodium tartrate (Rochelle salt, 2eq. per AlCl$_3$, 28.9 g) in H$_2$O (150 mL) and the mixture was vigorously stirred and shaken until all aluminum salts dissolved. Aqueous phase was further extracted with CH$_2$Cl$_2$ (4 × 100 mL), combined extracts were dried over Na$_2$SO$_4$, desiccant was filtered, and solution was concentrated to a pink foamy solid. Crude material was purified by repeated column chromatography (gradient of 20 to 40% EtOAc in hexanes + 2% Et$_3$N). Slightly impure product was repeatedly dissolved in aqueous ethanol and concentrated to remove all triethylamine trapped as the partial phenolate salt. Material was then dissolved in methanol (~15 mL per 1 g of solid) and acidified by a dropwise addition of concentrated hydrochloric acid (12.1 M, 36-38%), until the solution gave strongly acidic response on pH paper. Solution was concentrated, suspended in acetonitrile, concentrated again and solid thoroughly dried. Crude hydrochloride salt was suspended in acetonitrile (~10 mL per 1 g of solid), suspension was briefly heated to reflux (colored impurities dissolve) and cooled to room temperature. Oxa-noribogaine hydrochloride was collected by filtration, washed with acetonitrile (~5 mL per 1 g of solid) and diethyl ether and air dried. Material was further suspended in saturated sodium bicarbonate solution and repeatedly extracted with mixture of 9:1 CH$_2$Cl$_2$:isopropanol, until no further extraction was detected by TLC. Combined extracts were dried over sodium sulfate, filtered and concentrated. To remove traces of solvents used, the material was dissolved in aqueous ethanol and concentrated. The obtained foamy solid material was crushed to release trapped solvent residue and dried overnight at 45 °C under high vacuum. Oxa-noribogaine was obtained as an off-white (pale grey) amorphous solid (3.49 g, 67% over two steps), yield was corrected for 1.9% of ethanol remaining in the material even after extensive drying.

**$^1$H NMR (500 MHz, CDCl$_3$)** δ 7.19 (d, *J* = 8.6 Hz, 1H), 6.80 (d, *J* = 2.5 Hz, 1H), 6.70 (dd, *J* = 8.6, 2.6 Hz, 1H), 4.74 (br, 1H), 3.45 – 3.33 (m, 1H), 3.21 – 3.09 (m, 3H), 3.00 – 2.91 (m, 2H), 2.81 (d, *J* = 2.2 Hz, 1H), 2.48 – 2.36 (m, 1H), 2.09 – 1.99 (m, 1H), 1.89 – 1.76 (m, 2H), 1.64 (dq, *J* = 13.3, 3.2 Hz, 1H), 1.61 – 1.41 (m, 3H), 1.21 (ddt, *J* = 12.7, 6.5, 2.4 Hz, 1H), 0.91 (t, *J* = 7.1 Hz, 3H); **$^{13}$C NMR (126 MHz, CDCl$_3$)** δ 161.1, 151.4, 148.6, 131.8, 111.64, 111.55, 110.9, 104.2, 57.3, 53.3, 49.6, 41.6, 41.1, 33.0, 32.2, 27.5, 26.4, 19.4, 12.0; **HR-MS** calcd. for $C_{19}H_{24}NO_2^+$ [M+H]$^+$ 298.1807, found 298.1818.

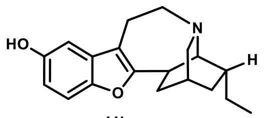


**11b**
**Epi-oxa-noribogaine**

***rac*-epi-oxa-noribogaine (Epi-oxa, 11b)**

The product **11b** was prepared according to the general procedure and purified by column chromatography (20:1 $CH_2Cl_2$:MeOH, 4 column volumes → 20:1 acetone:MeOH, 4 column volumes) to provide a white, foamy solid (13.7 mg, 92%).

**$^1$H NMR (500 MHz, $CDCl_3$)** δ 7.19 (d, *J* = 8.7 Hz, 1H), 6.79 (d, *J* = 2.4 Hz, 1H), 6.71 (dd, *J* = 8.7, 2.5 Hz, 1H), 5.62 (br s, 1H), 3.40 (ddd, *J* = 14.2, 4.6, 2.4 Hz, 1H), 3.34 – 3.26 (m, 2H), 3.20 – 3.08 (m, 2H), 3.02 (d, *J* = 9.8 Hz, 1H), 2.91 (s, 1H), 2.43 (dt, *J* = 16.6, 2.9 Hz, 1H), 2.08 – 1.95 (m, 3H), 1.90 (s, 1H), 1.65 – 1.58 (m, 1H), 1.44 – 1.33 (m, 2H), 1.15 – 1.09 (m, 1H), 0.92 (t, *J* = 7.3 Hz, 3H); **$^{13}$C NMR (126 MHz, $CDCl_3$)** δ 161.5, 152.1, 148.2, 131.4, 112.11, 112.06, 111.0, 104.2, 56.5, 53.6, 49.0, 41.2, 34.0, 33.9, 31.4, 28.4, 26.1, 18.8, 12.3.; **HR-MS** calcd. for $C_{19}H_{24}NO_2^+$ [M+H]$^+$ 298.1807, found 298.1818.

**General Procedure for Preparation of Oxa-ibogaine Analogs by lithiation/iodination and reductive Heck sequence**.

Uncyclized intermediate **9a**/**9b** (1 equivalent) was dissolved in THF (0.5 M based on **9a**/**9b**) under argon atmosphere and the solution was cooled to -40°C in acetone/dry ice bath. Solution of n-butyl lithium in hexanes (2.5 M, 2 equivalents) was added dropwise over 5 – 10 min and the resulting orange solution was stirred 1 h at -40°C. Solution of iodine (1.5 - 1.7 equivalents) in THF (same volume as for **9a**/**9b**) was added ($I_2$ is consumed during addition, but persistent coloration remains after the entire amount is added) and after stirring for 10 min at -40°C the cooling bath was removed, and reaction allowed to warm to room temperature. After stirring at room temperature for 1 h reaction was quenched with addition of saturated $Na_2S_2O_3$ solution (1 mL per 1 mmol of **9a**/**9b**) and the resulting mixture was vigorously stirred until excess iodine was consumed (org. phase discolored to pale orange-brown). Mixture was then poured into water, extracted with diethyl ether (3×), combined extracts were dried over $Na_2SO_4$, filtered and concentrated. Crude material (from **9a** pale orange, from **9b** orange-brown oil, can solidify over time) was used for next step without further purification. Largest scale tested for **9a** (5.05 mmol, 1.57 g) and **9b** (2.79 mmol, 0.87 g).

*Note: Despite using excess n-BuLi and $I_2$ reaction never reached full conversion, even when $I_2$ was added in excess to n-BuLi. Typically, 3 – 10% of unreacted starting material remains and is difficult to separate. No improvement in conversion was observed by increasing the reaction temperature after addition of n-BuLi to 0°C or using freshly opened commercially available anhydrous THF and n-BuLi solution.*

Crude iodo-intermediate (1 equivalent), sodium formate (4 equivalents, powdered and dried overnight on high-vacuum) and bis[tri(o-tolyl)phosphine]palladium(II) chloride (1 mol%, 0.01 equivalent) were combined in dimethyl sulfoxide (0.25 M based on iodo-intermediate) under argon atmosphere. Closed reaction vessel was placed in a pre-heated (130°C) heating adapter. Upon reaching reaction temperature (2 to 10 min, depending on volume and reaction vessel) the mixture (pale orange for **10a** or orange-brown for **10b**) turned dark brown/black. Reaction was further stirred at 130°C for 5 - 60 min. After complete conversion was observed, reaction was cooled to room temperature and the dark mixture was poured to water (~5× volume of DMSO) and extracted with diethyl ether (3-4×, until no further extraction was observed). Combined extracts were dried over $Na_2SO_4$, filtered and concentrated. Obtained crude material was purified as indicated for each derivative.

*Note: In the course of reaction scale-up it was observed that starting material is consumed upon reaching reaction temperature (exact time depends on reaction volume and vessel used). Subsequently, reaction time was reduced to 5 minutes after color changed due to precipitation of reduced palladium was observed. Two repeats for oxa-ibogaine **10a** suggest possible improvement in yield by shortening the reaction time or lowering the reaction temperature. Palladium loading <1 mol% was not tested at this time.*

**Oxa-ibogaine (10a)**

Prepared according to the general procedure, reaction was repeated 3 times on (1.88, 2.0 and 5.05 mmol scale). The crude reaction mixture after cyclization was purified directly by column chromatography (30:1 hexanes:EtOAc + 1% $Et_3N$) to yield the product as an off white solid. The slightly impure material, typically (>95% purity) was used for next step as is, yield of first repeat over two steps (0.47 g, 80%), reaction time 60 min. For second (0.52 g 86%) and third repeat (1.42 g, 90%), reactions were terminated 5 minutes after reduction of palladium was observed, total reaction time <15 minutes. Yields are corrected for presence of uncyclized material **9a** that can be removed after transformation to **11a,** as indicated in the multigram scale Ni-based synthesis note.

Spectral characterization of **10a** is identical to the material prepared by Ni mediated cyclization reaction.

**Epi-oxa-ibogaine (10b)**

Prepared according to the general procedure, reaction was repeated 3 times on (1.0, 1.03, and 6.74 mmol scale). The crude reaction mixture after cyclization was purified directly by column chromatography (gradient of 15 to 20% EtOAc in hexanes + 2% $Et_3N$) to yield the product as an off white solid. Yields over two steps (81%, 252 mg) and (250 mg, 78%). Third repeat (1.32 g, 63%), initial palladium catalyst was reduced upon reaching reaction temperature, but no conversion of iodo-intermediate was detected. Full conversion was achieved by addition of extra 4 equiv. HCOONa and 1 mol% of $Pd[P(o\text{-tolyl})_3]_2Cl_2$ to the same reaction mixture and continued heating for 15 min. During purification a mixed fraction containing 83:17 **9b**:**10b** was also recovered (0.37 g, 19%). Deviation from previous repeats were higher reaction scale, different batch of solvent and reagents (SM, HCOONa and Pd cat.) were pre-dried for 1 h before addition of solvent at 40°C instead of only HCOONa overnight at room temperature.

Spectral characterization of **10b** is identical to the material prepared by Ni mediated cyclization reaction.

### X-Ray structure determination of Oxa-ibogaine 10a

X-Ray suitable crystals were prepared by dissolving racemic oxa-ibogaine in hot acetonitrile and the resulting solution was allowed to cool to room temperature and slowly concentrate.

X-ray diffraction data were collected on a Bruker Apex II diffractometer. The structures were solved by using direct methods and standard difference map techniques and were refined by full-matrix least-squares procedures on F2 with SHELXTL (Version 2014/7)[40–42]. Crystallographic data have been deposited with the Cambridge Crystallographic Data Centre (CCDC #2215015).

| **Property** | **Oxa-ibogaine 10a** | **Property** | **Oxa-ibogaine 10a** |
|---|---|---|---|
| *Formula* | C20H25NO2 | *Radiation type* | MoK α |
| *Formula Weight* | 311.41 | *Wavelength (λ, Å)* | 0.71073 |
| $D_{calc.}$/ $g{\cdot}cm^{-3}$ | 1.272 | *θ min, deg.* | 2.33 |
| $Mu/mm^{-1}$ | 0.081 | *θ max, deg.* | 30.55 |
| *Color* | colorless | *Measured Refl.* | 25983 |
| *Shape* | block | *Independent Refl.* | 4982 |
| $Size/mm^3$ | 0.43×0.11×0.07 | *Reflections with I > 2(I)* | 3804 |
| *Space Group* | $P2_1/n$ | $R_{int}$ | 0.0432 |
| *Crystal System* | Monoclinic | *Parameters* | 210 |
| *a/Å* | 10.9618(18) | *Restraints* | 0 |
| *b/Å* | 8.4800(14) | *Density Max* | 0.434 |
| *c/Å* | 17.660(3) | *Density Min* | -0.211 |
| *α/°* | 90 | *GoF* | *1.045* |
| *β/°* | 97.742(3) | *wR2 (all data)* | *0.1217* |
| *γ/°* | 90 | *wR2* | *0.1100* |
| $V/Å^3$ | 1626.6(5) | *R1 (all data)* | *0.0625* |
| *Z* | 4 | *R1* | *0.0441* |
| *Temperature (K)* | 150(2) | | |

### Appendix 1. High Resolution Mass Spectra

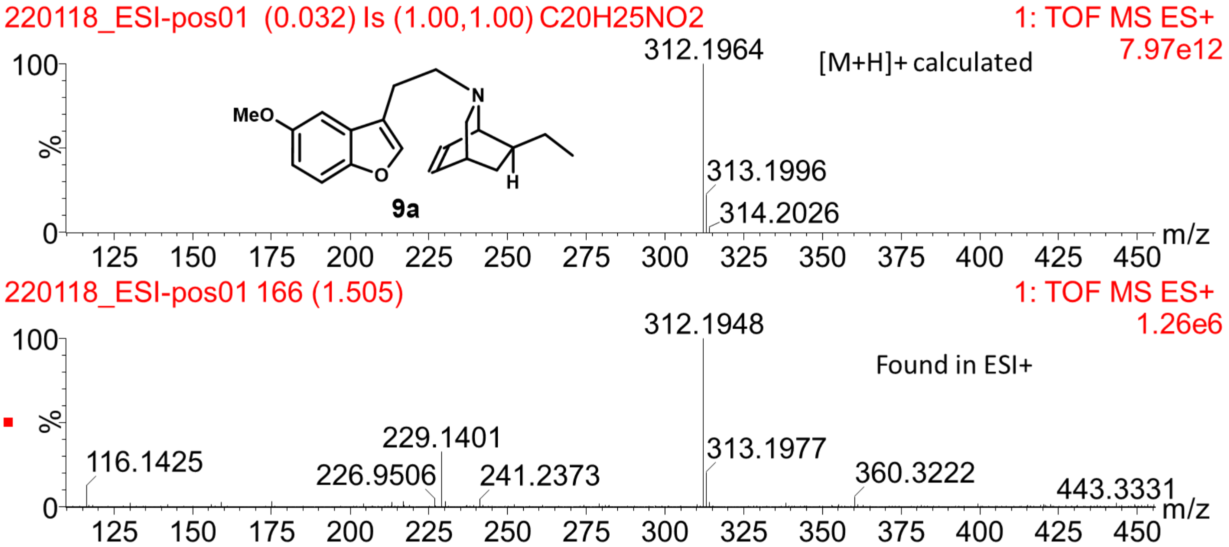


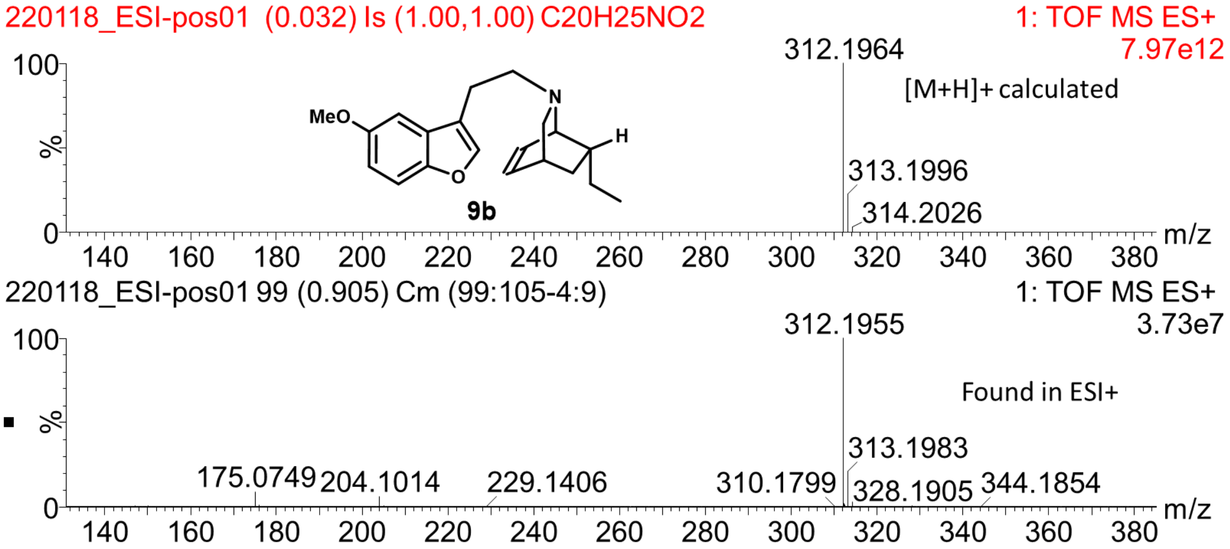

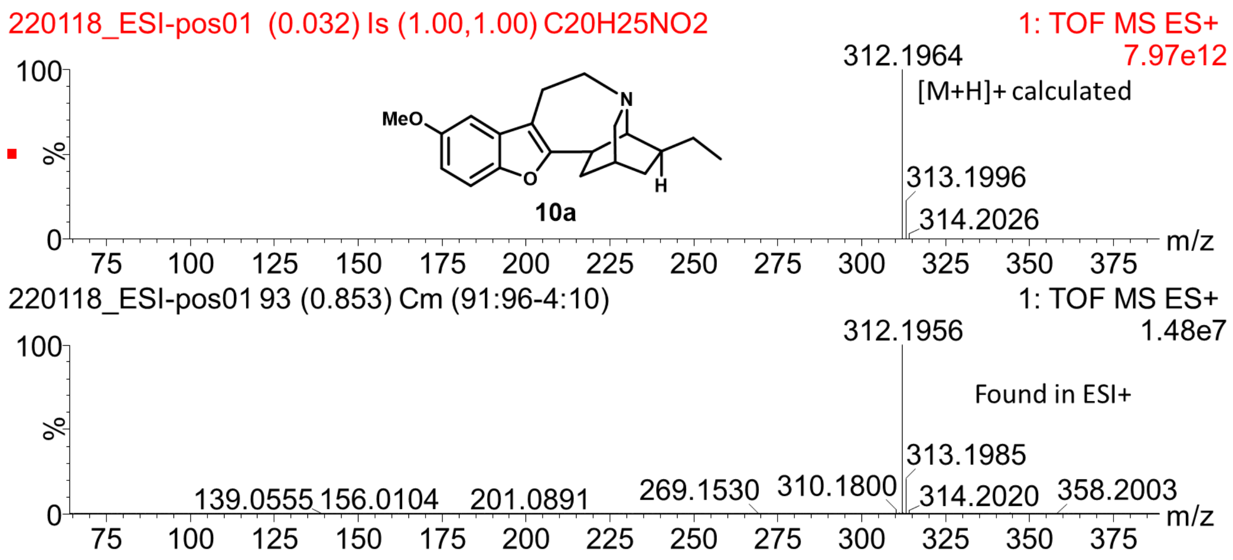
220118_ESI-pos01 (0.032) Is (1.00,1.00) C20H25NO2
1: TOF MS ES+
7.97e12
312.1964
[M+H]+ calculated
313.1996
314.2026
MeO
N
O
H
10a
220118_ESI-pos01 93 (0.853) Cm (91:96-4:10)
1: TOF MS ES+
1.48e7
312.1956
Found in ESI+
313.1985
139.0555
156.0104
201.0891
269.1530
310.1800
314.2020
358.2003
%
m/z

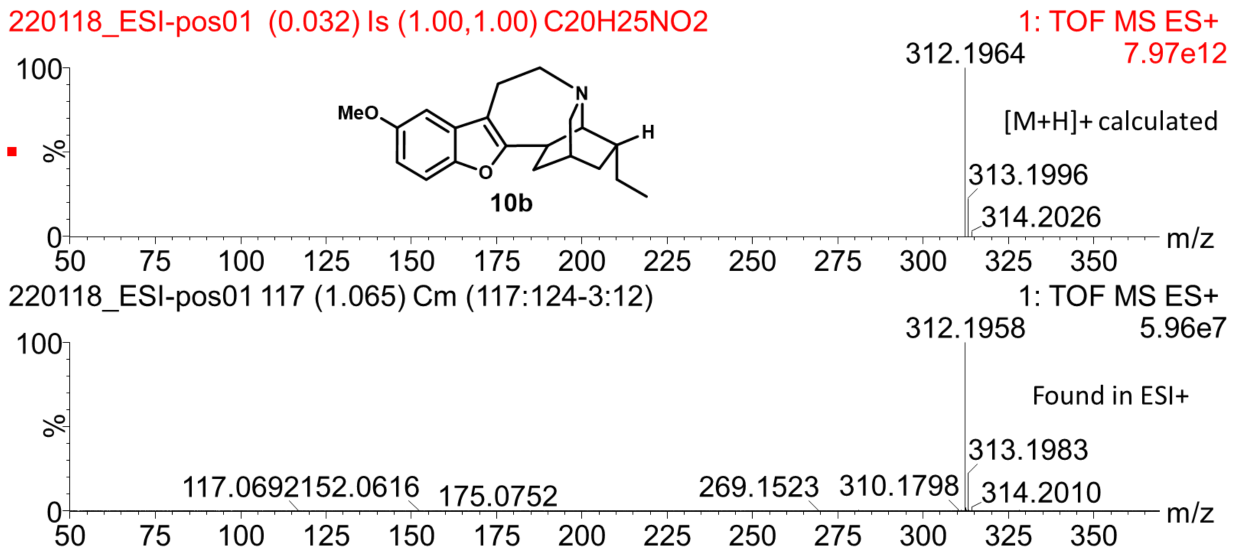
220118_ESI-pos01 (0.032) Is (1.00,1.00) C20H25NO2
1: TOF MS ES+
7.97e12
312.1964
[M+H]+ calculated
313.1996
314.2026
MeO
N
O
H
10b
220118_ESI-pos01 117 (1.065) Cm (117:124-3:12)
1: TOF MS ES+
5.96e7
312.1958
Found in ESI+
313.1983
117.0692
152.0616
175.0752
269.1523
310.1798
314.2010
%
m/z

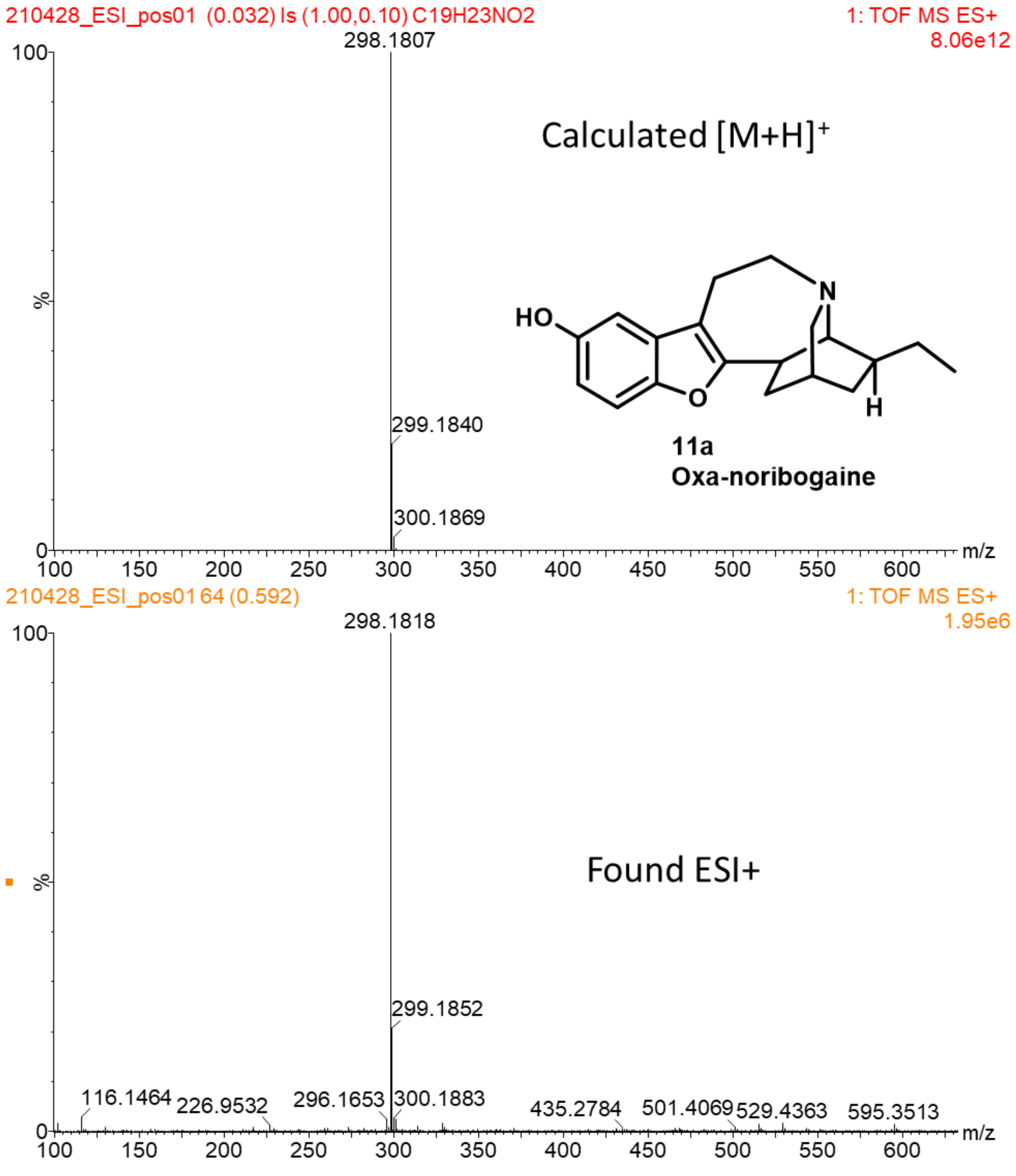
210428_ESI_pos01 (0.032) Is (1.00,0.10) C19H23NO2
1: TOF MS ES+
8.06e12
298.1807
299.1840
300.1869
Calculated [M+H]+
HO
N
O
H
11a
Oxa-noribogaine
%
m/z
210428_ESI_pos01 64 (0.592)
1: TOF MS ES+
1.95e6
298.1818
299.1852
116.1464
226.9532
296.1653
300.1883
435.2784
501.4069
529.4363
595.3513
Found ESI+

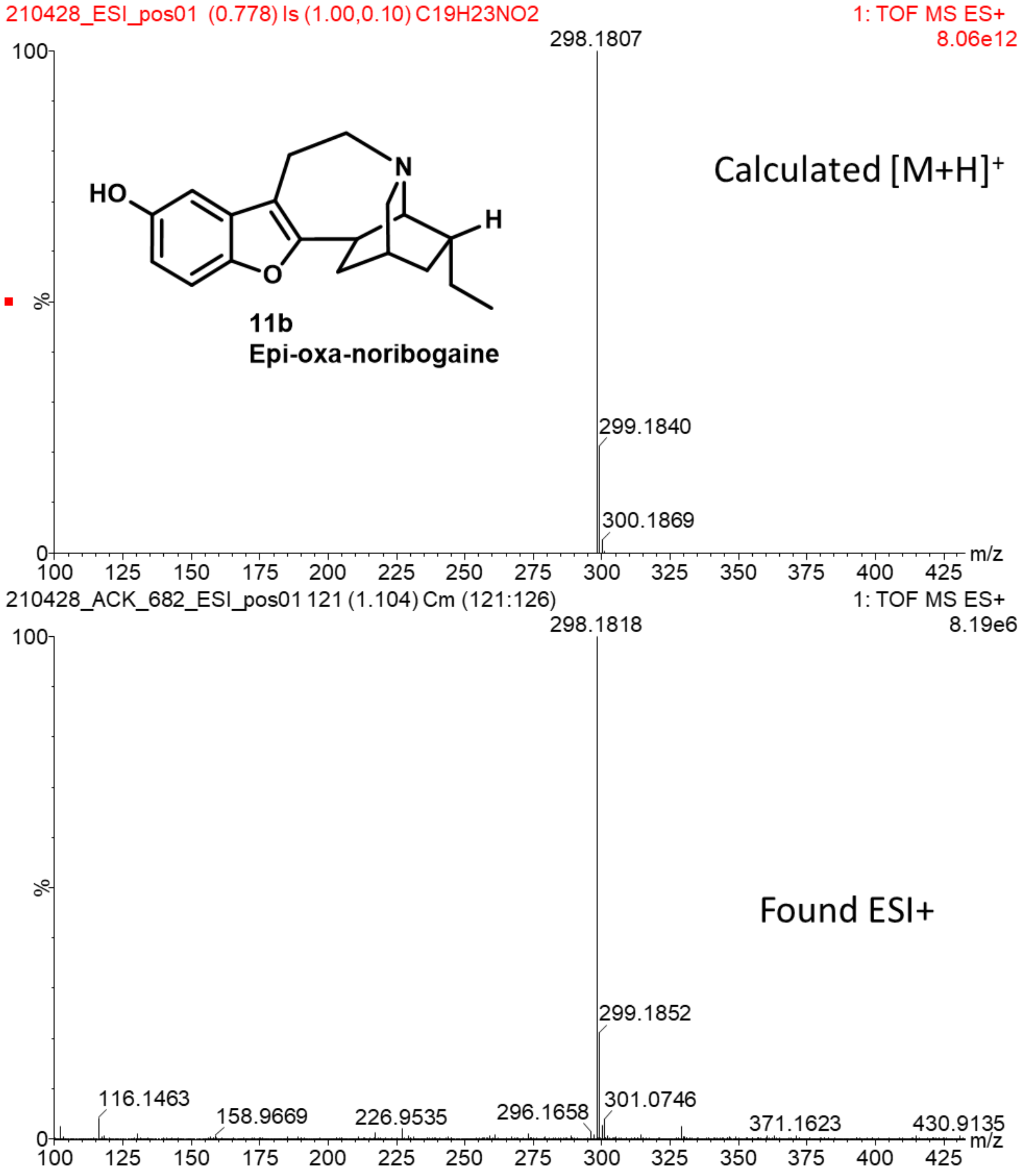
210428_ESI_pos01 (0.778) Is (1.00,0.10) C19H23NO2
1: TOF MS ES+
8.06e12
298.1807
299.1840
300.1869
HO
N
H
O
11b
Epi-oxa-noribogaine
Calculated [M+H]+
m/z
210428_ACK_682_ESI_pos01 121 (1.104) Cm (121:126)
1: TOF MS ES+
8.19e6
298.1818
299.1852
116.1463
158.9669
226.9535
296.1658
301.0746
371.1623
430.9135
Found ESI+
m/z

### Appendix 2. NMR Spectra

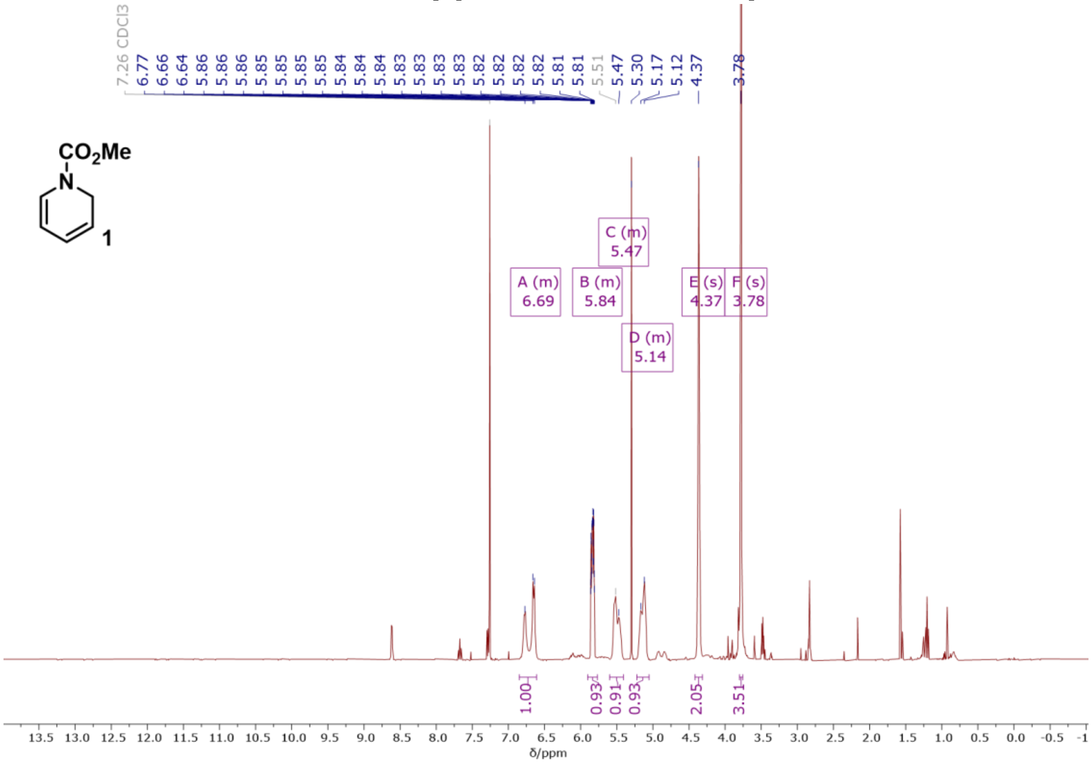


[1]H NMR (400 MHz, $CDCl_3$) spectrum of a crude, unstable intermediate **1**.

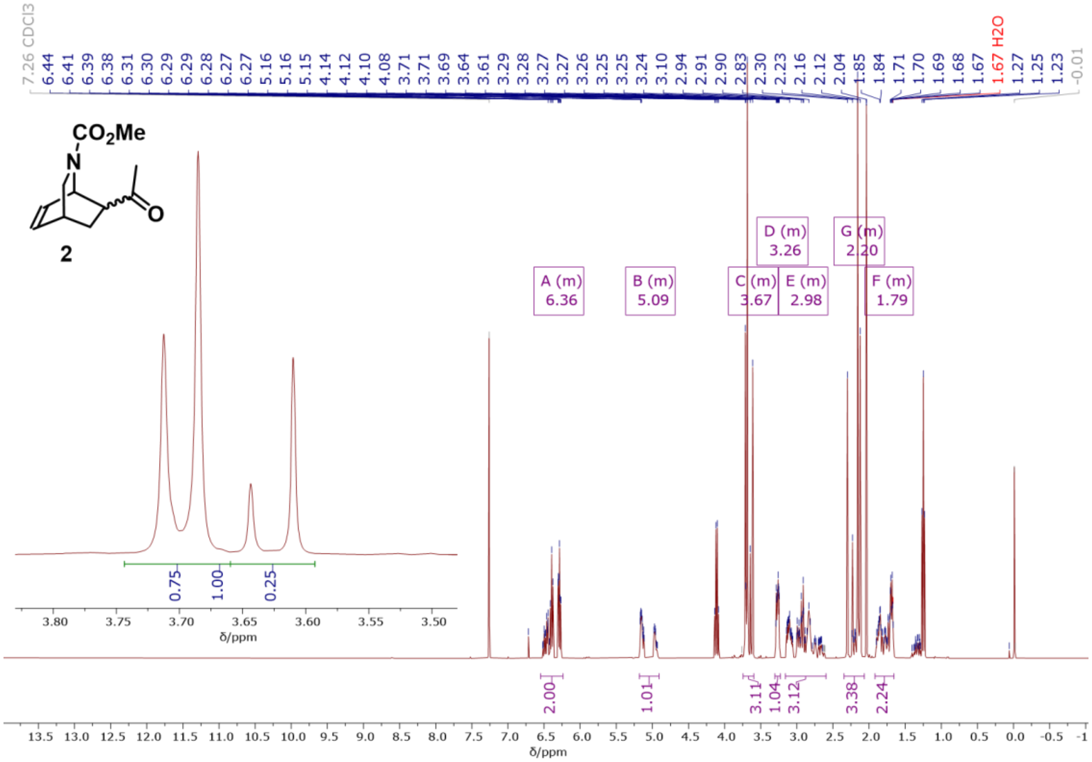


[1]H NMR (400 MHz, $CDCl_3$) spectrum of a mixture of **2a** and **2b** prior to epimerization.

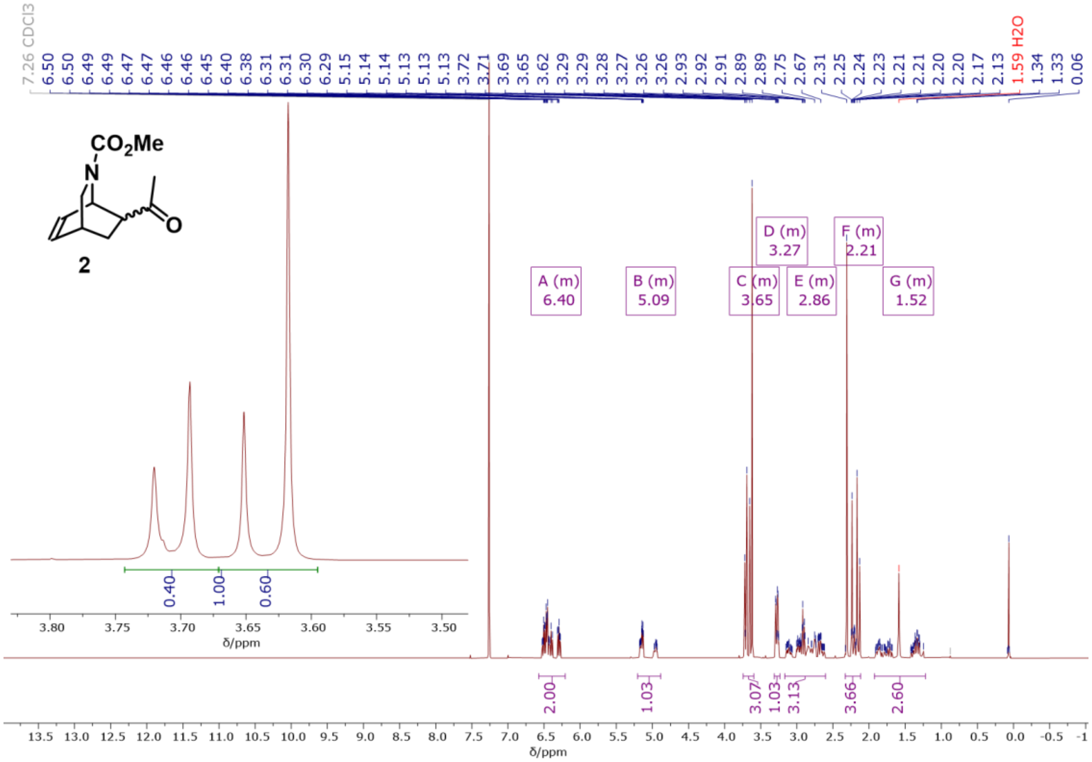


$^1$H NMR (400 MHz, $CDCl_3$) spectrum of a mixture of **2a** and **2b** after epimerization.

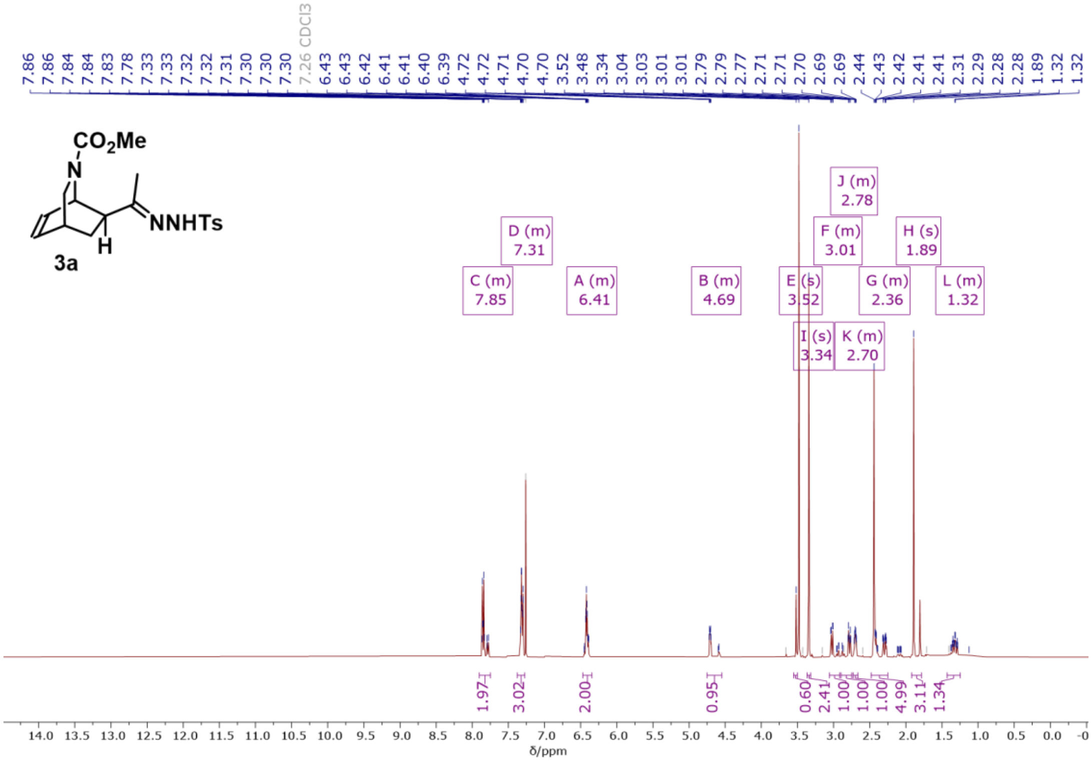


$^1$H NMR (400 MHz, $CDCl_3$) spectrum of compound 3**a**.

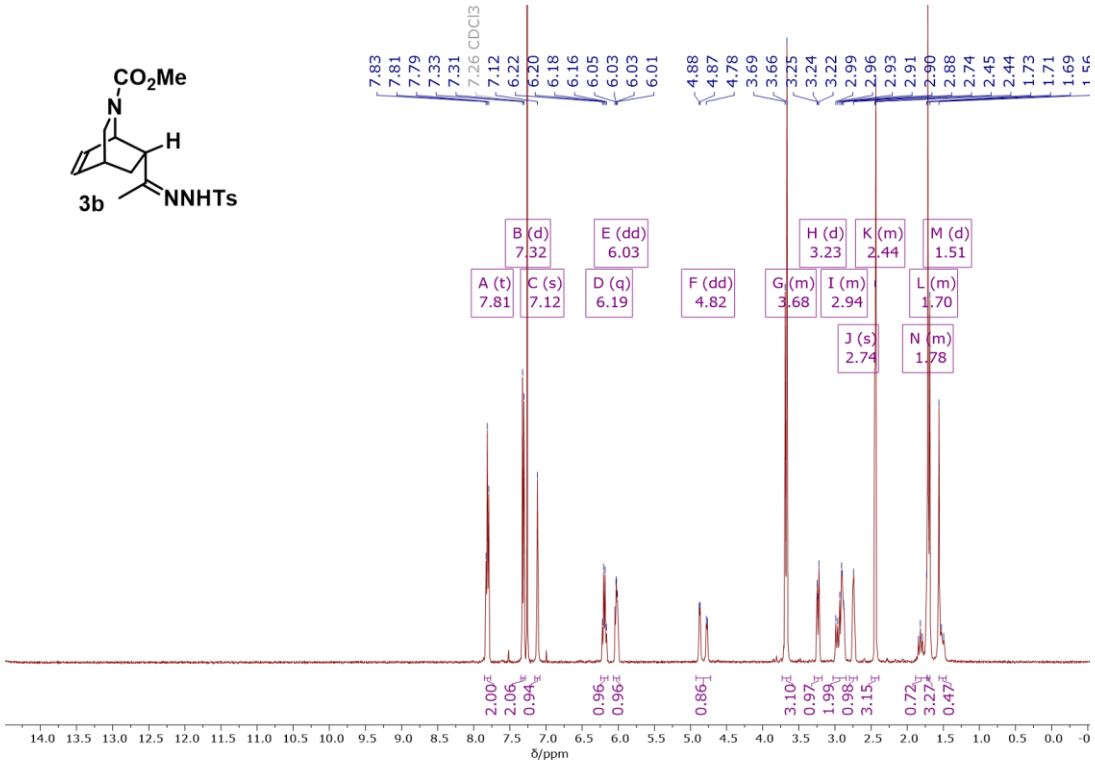


[1]H NMR (400 MHz, $CDCl_3$) spectrum of compound 3**b.**

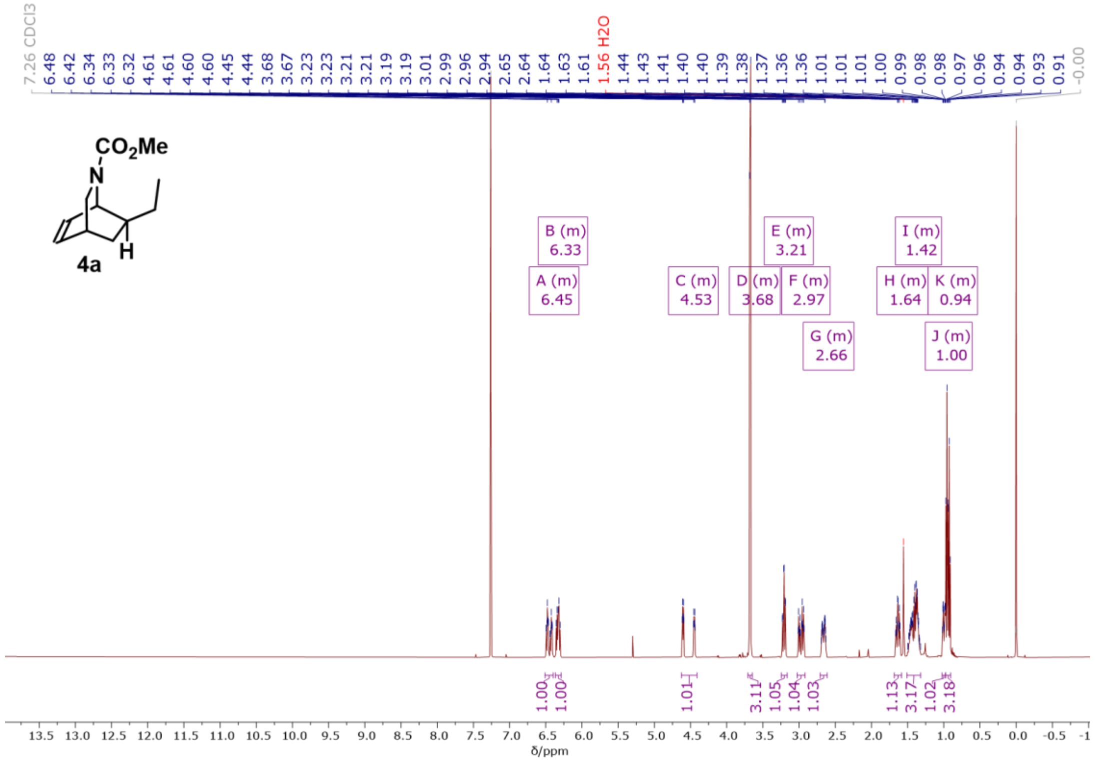


[1]H NMR (500 MHz, $CDCl_3$) spectrum of compound 4**a**.

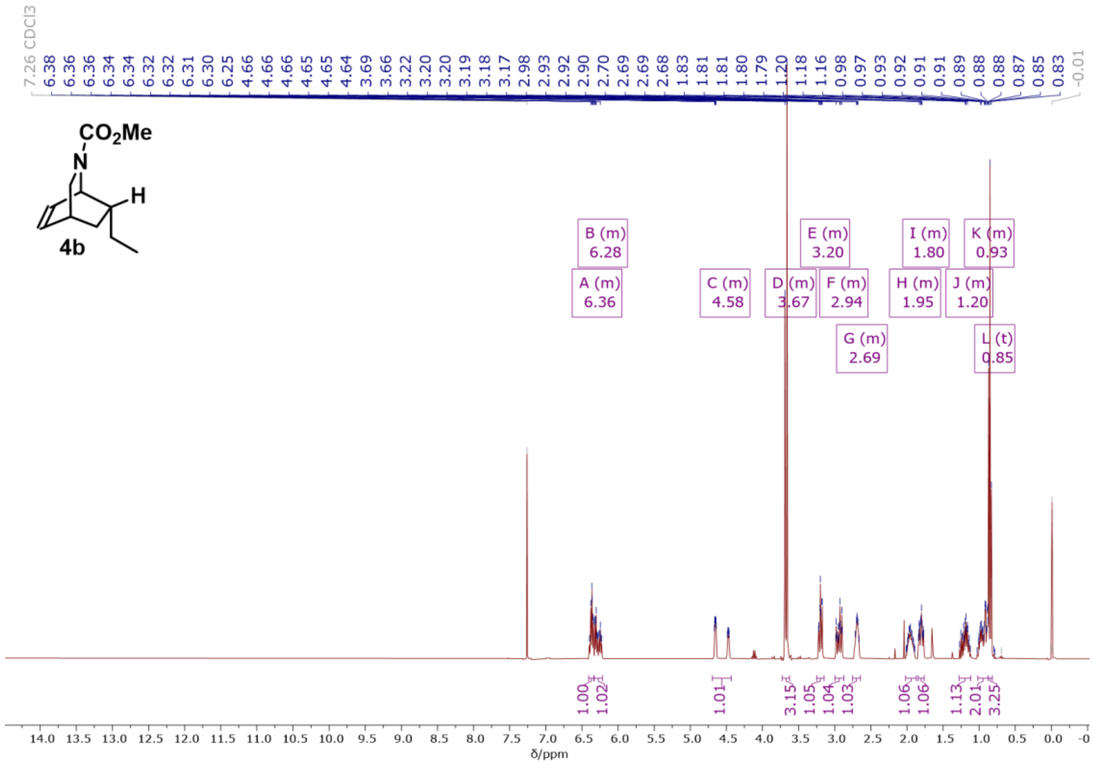

$^1$H NMR (400 MHz, $CDCl_3$) spectrum of compound 4**b.**

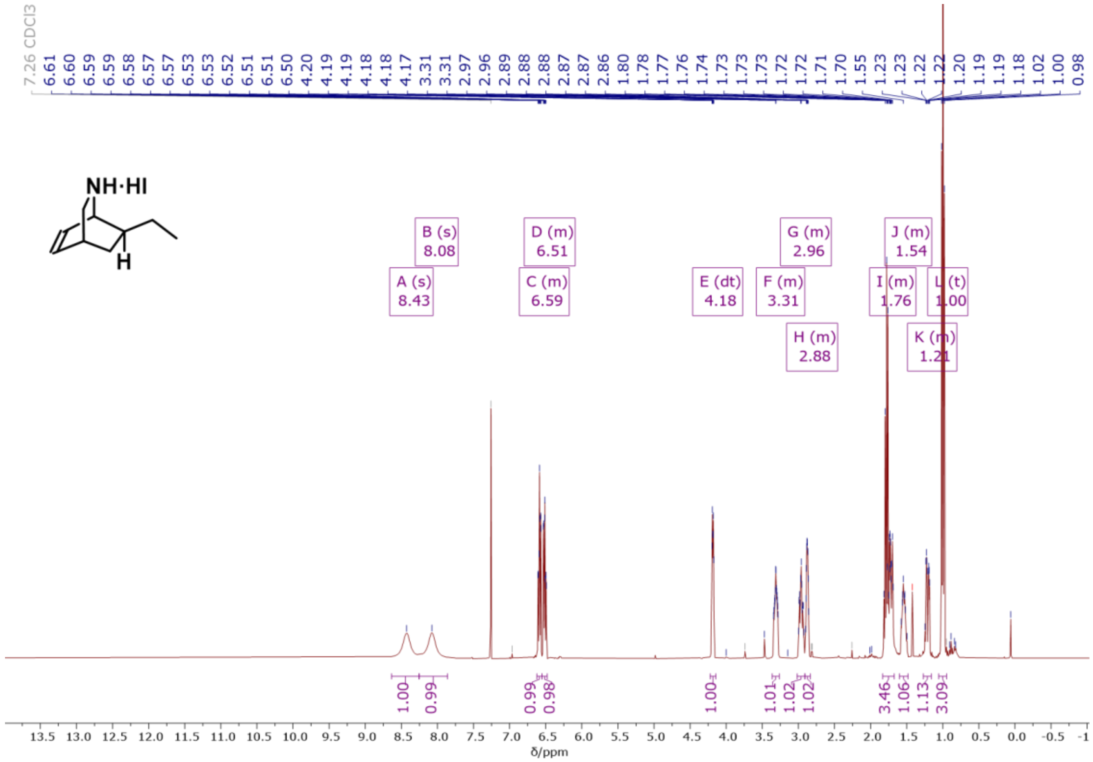

$^1$H NMR (400 MHz, $CDCl_3$) spectrum of exo-ethyl isoquinuclidine hydroiodide intermediate.

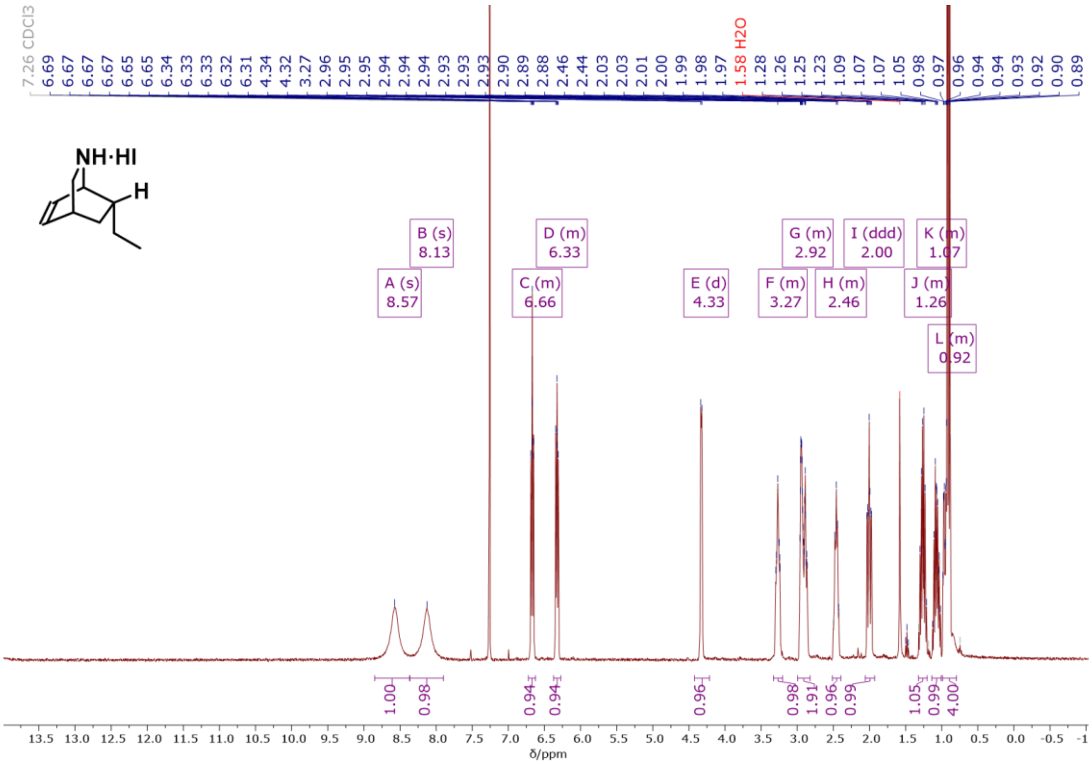


$^{1}$H NMR (400 MHz, $CDCl_3$) spectrum of endo-ethyl isoquinuclidine hydroiodide intermediate.

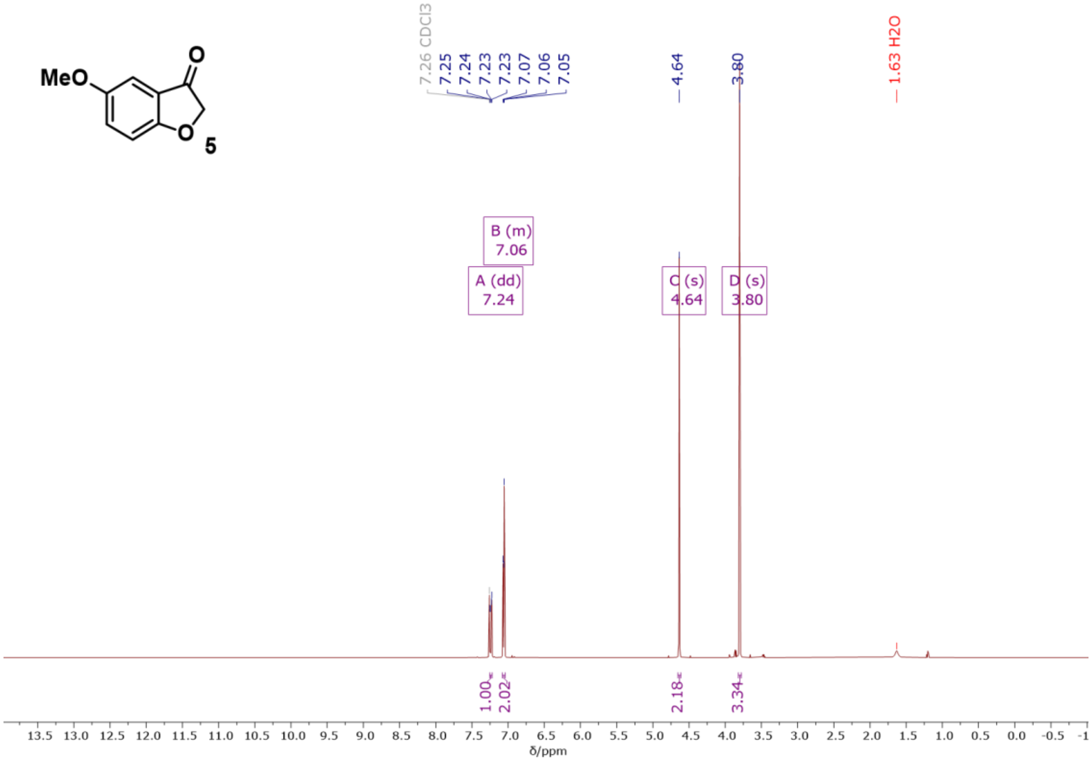


$^{1}$H NMR (400 MHz, $CDCl_3$) spectrum of compound **5**.

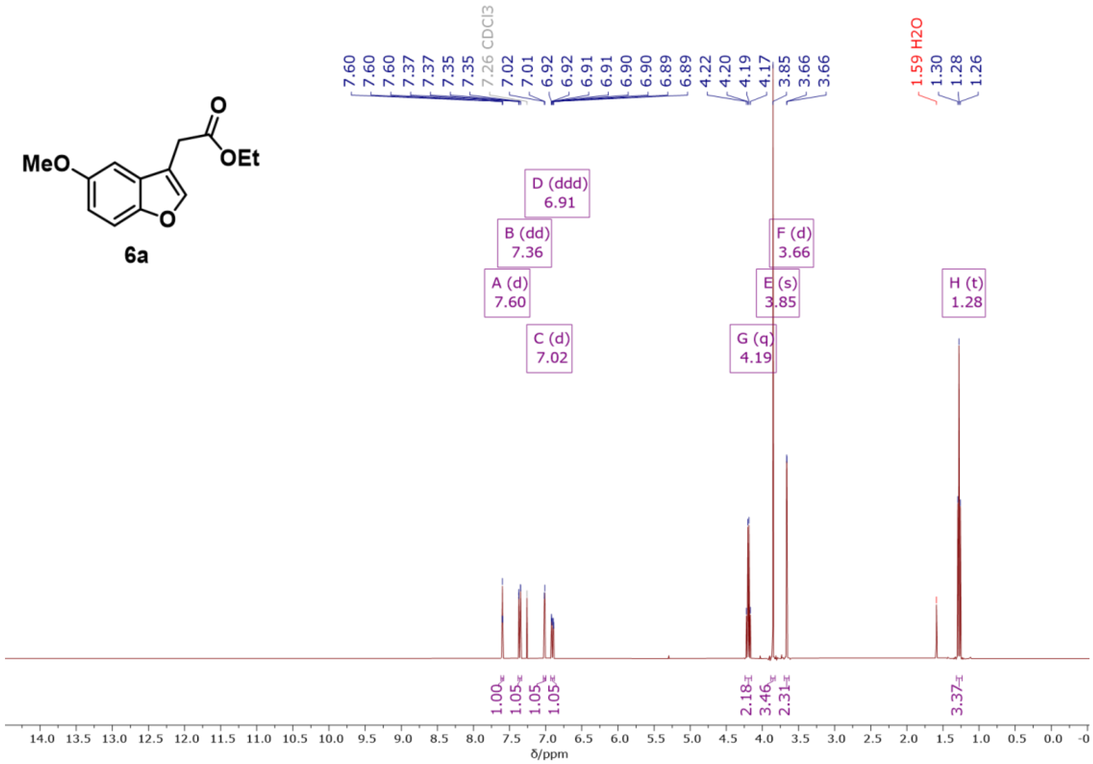


[1]H NMR (400 MHz, $CDCl_3$) spectrum of compound **6a.**

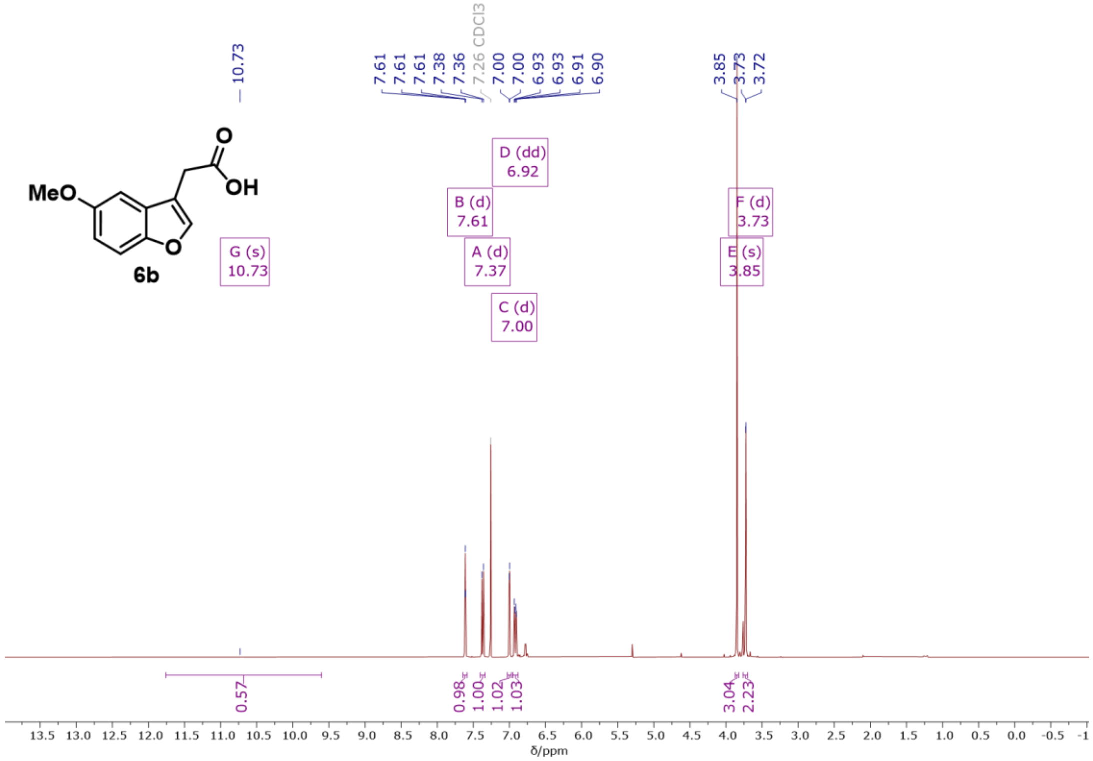


[1]H NMR (400 MHz, $CDCl_3$) spectrum of crude compound **6b**.

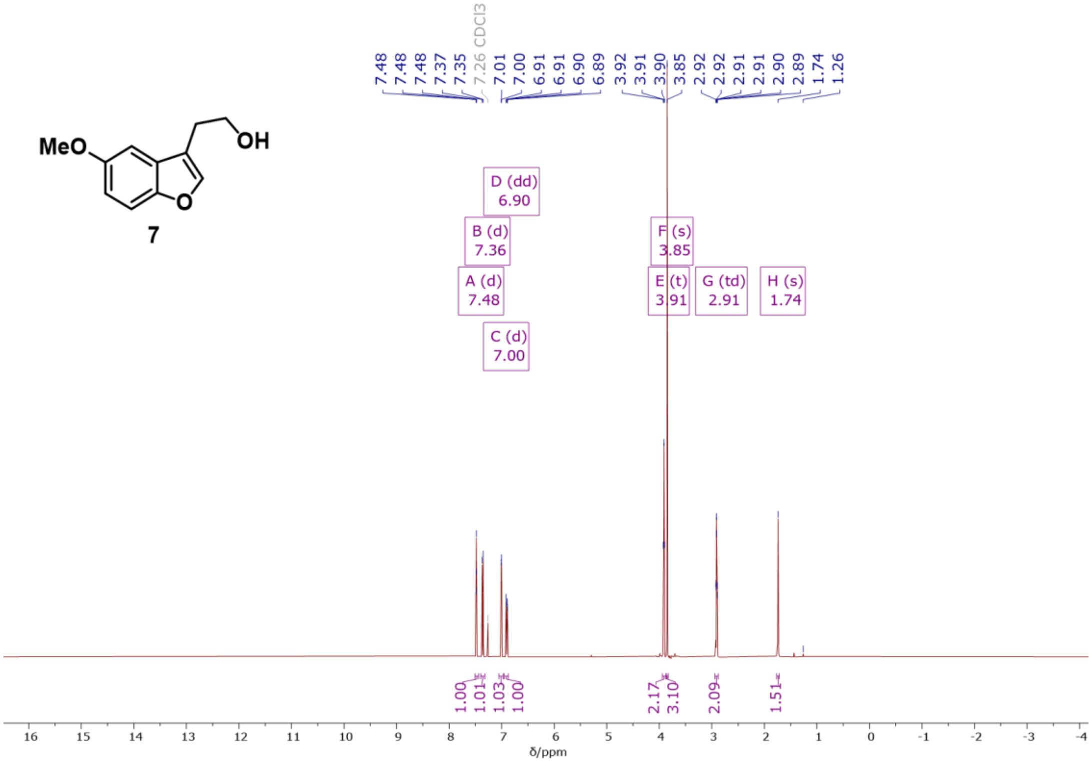


$^1$H NMR (500 MHz, $CDCl_3$) spectrum of compound **7**.

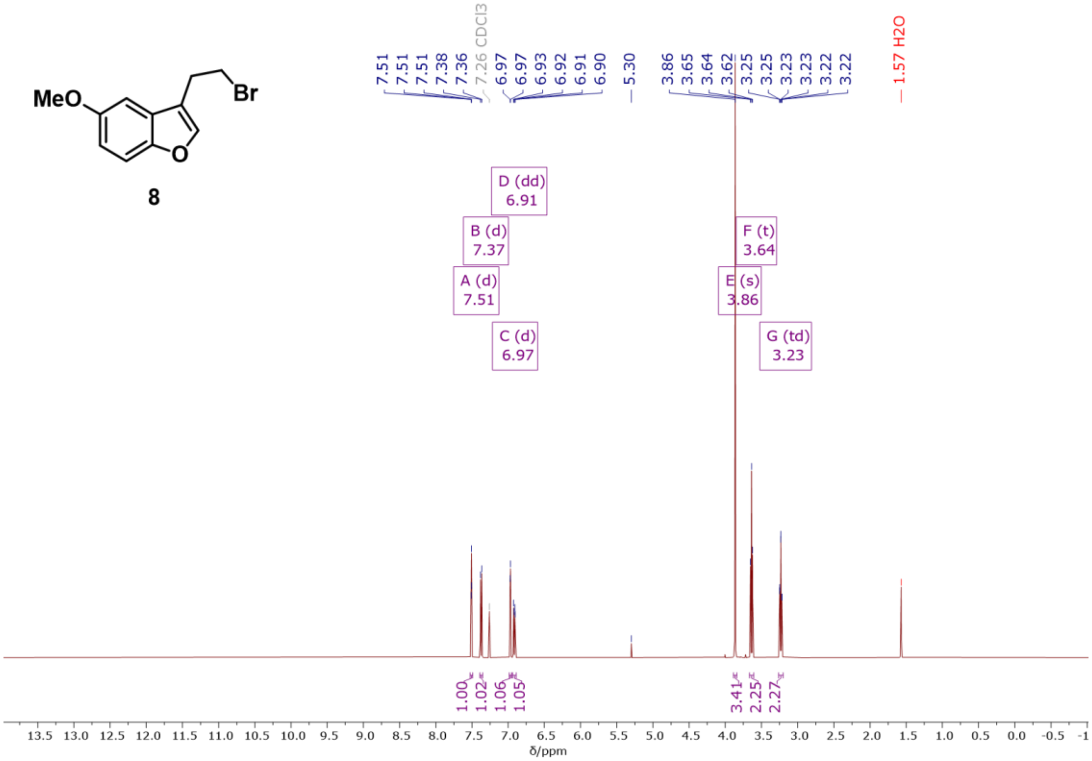


$^1$H NMR (500 MHz, $CDCl_3$) spectrum of compound **8**.

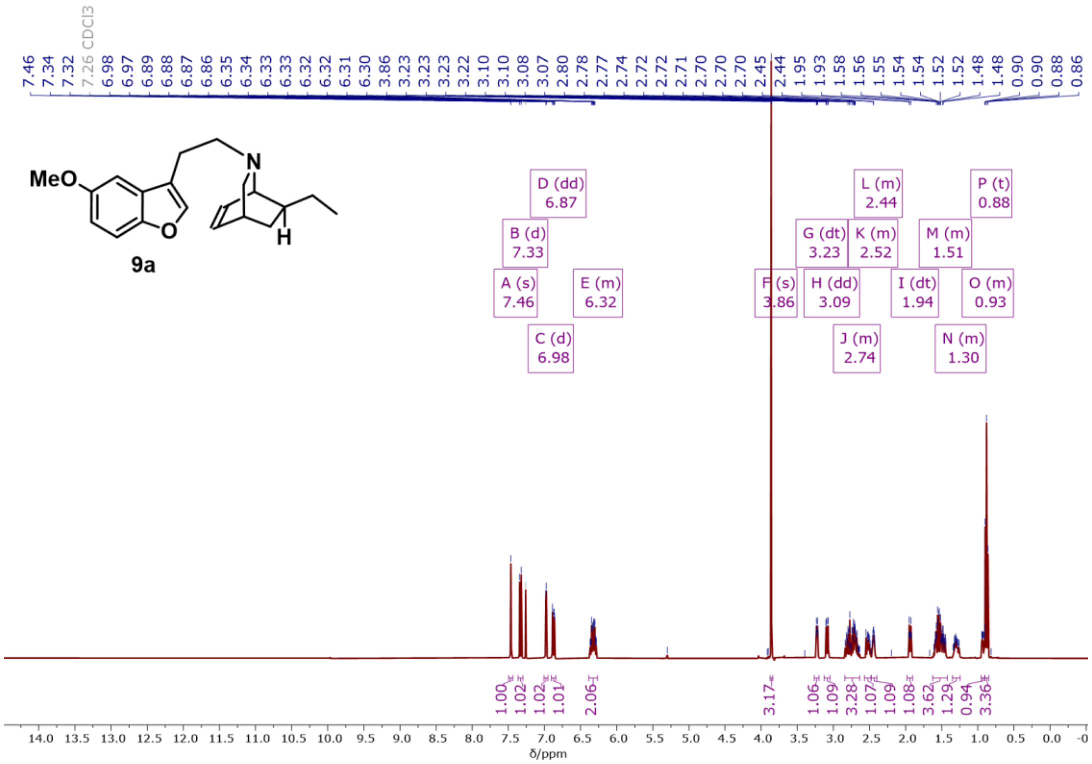


$^{1}H$ NMR (400 MHz, $CDCl_3$) spectrum of compound **9a.**

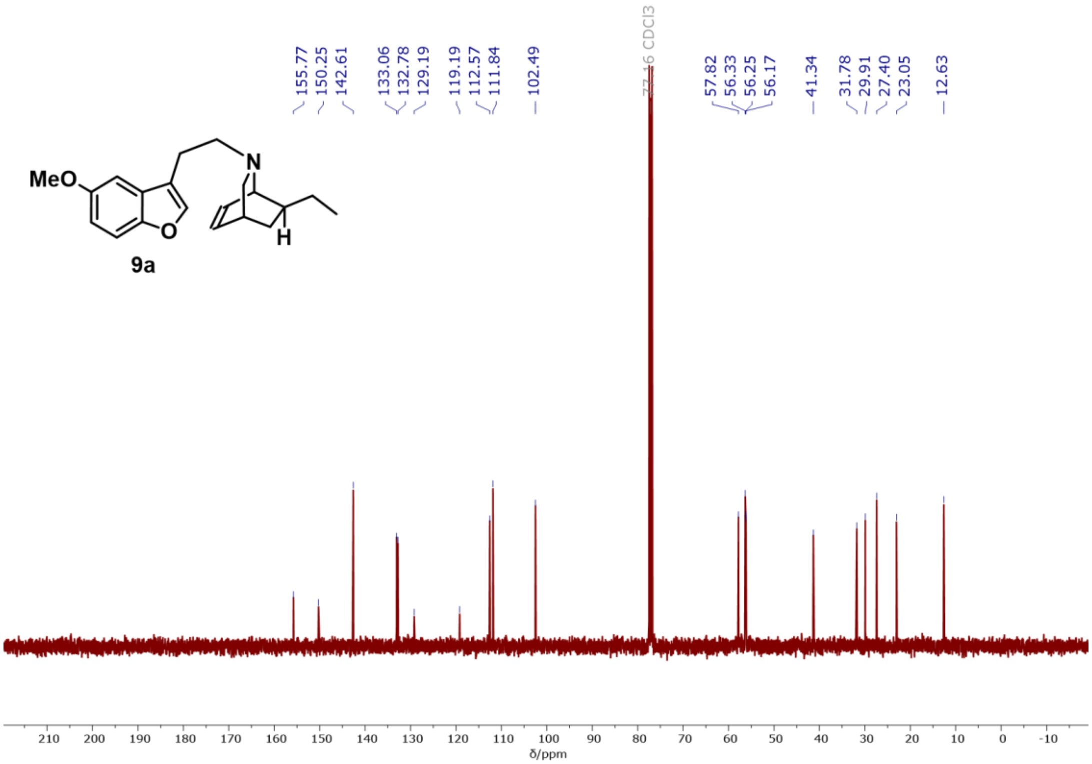


$^{13}C\{^{1}H\}$ NMR (101 MHz, $CDCl_3$) spectrum of compound **9a.**

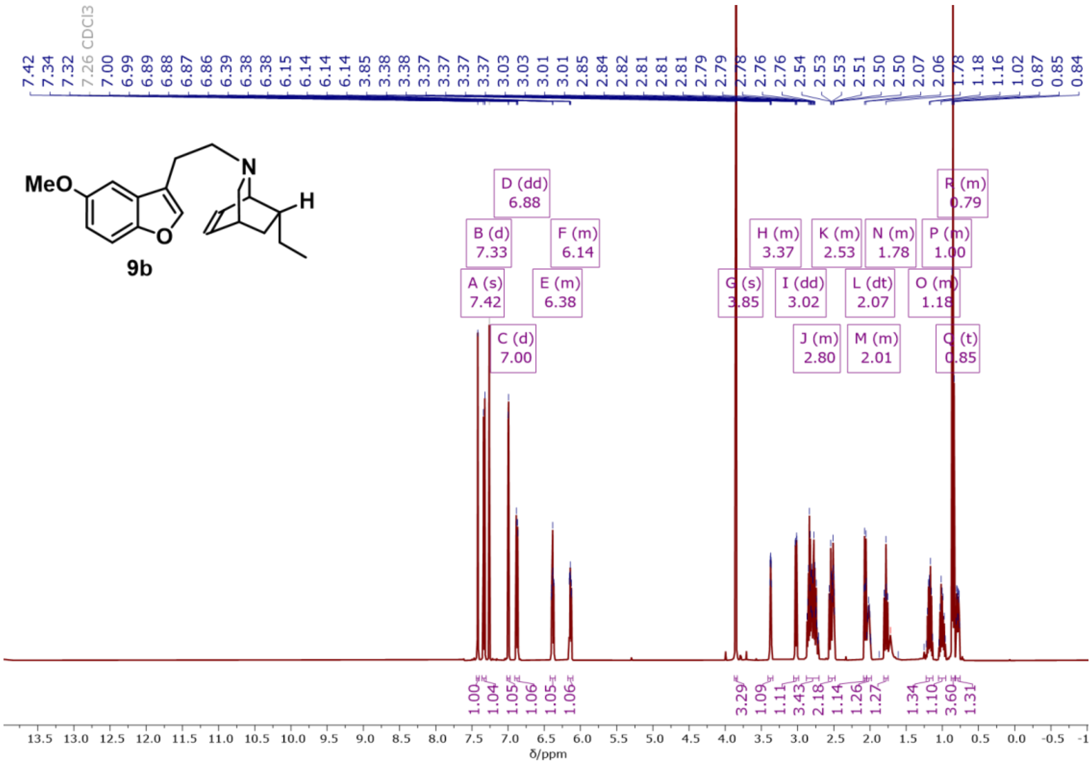


[1]H NMR (400 MHz, $CDCl_3$) spectrum of compound **9b.**

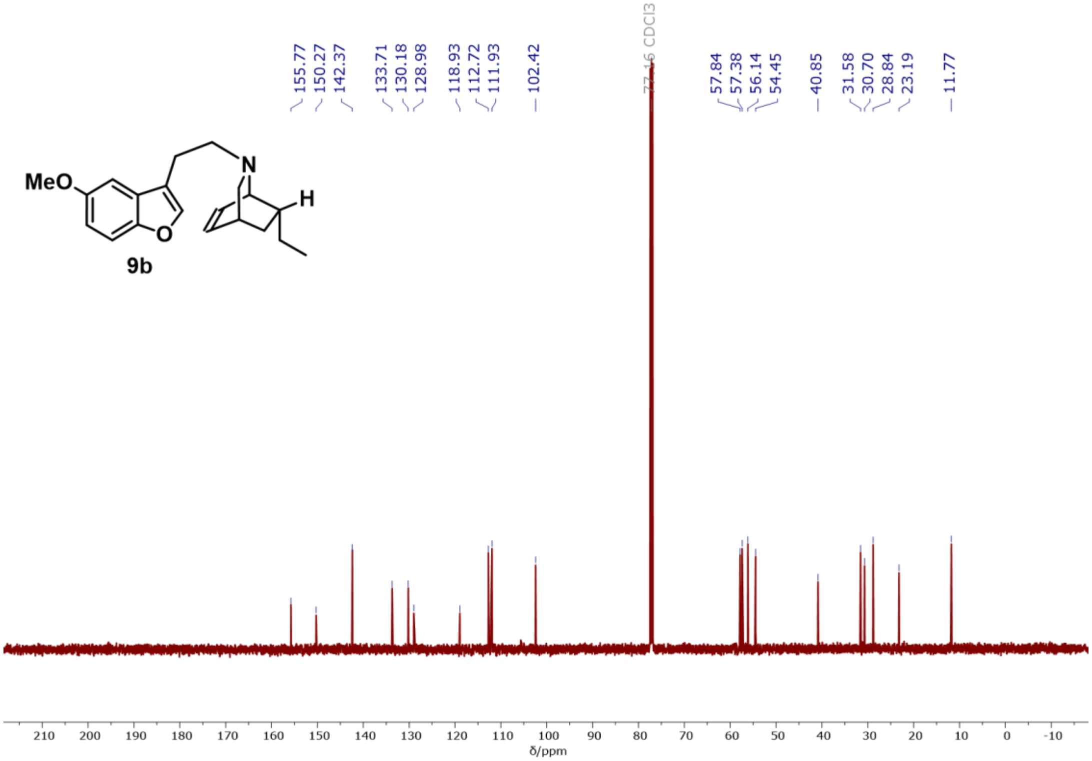


[13]C{[1]H} NMR (101 MHz, $CDCl_3$) spectrum of compound **9b.**

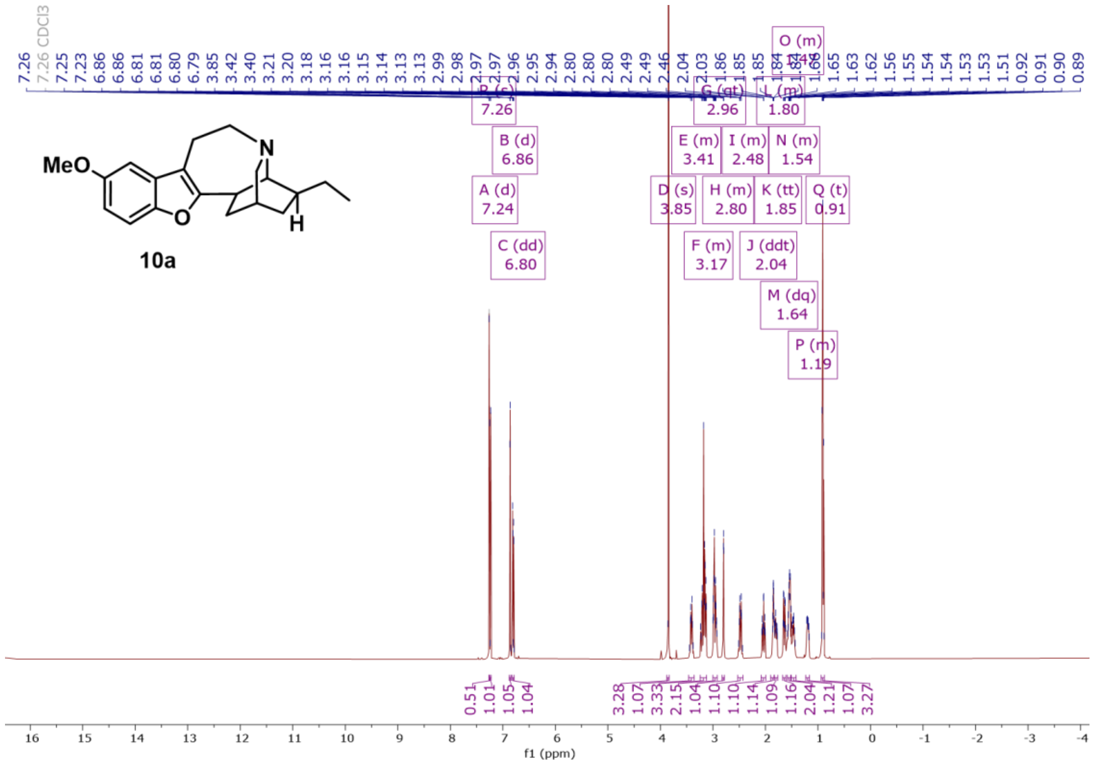

[1]H NMR (500 MHz, $CDCl_3$) spectrum of compound **10a.**

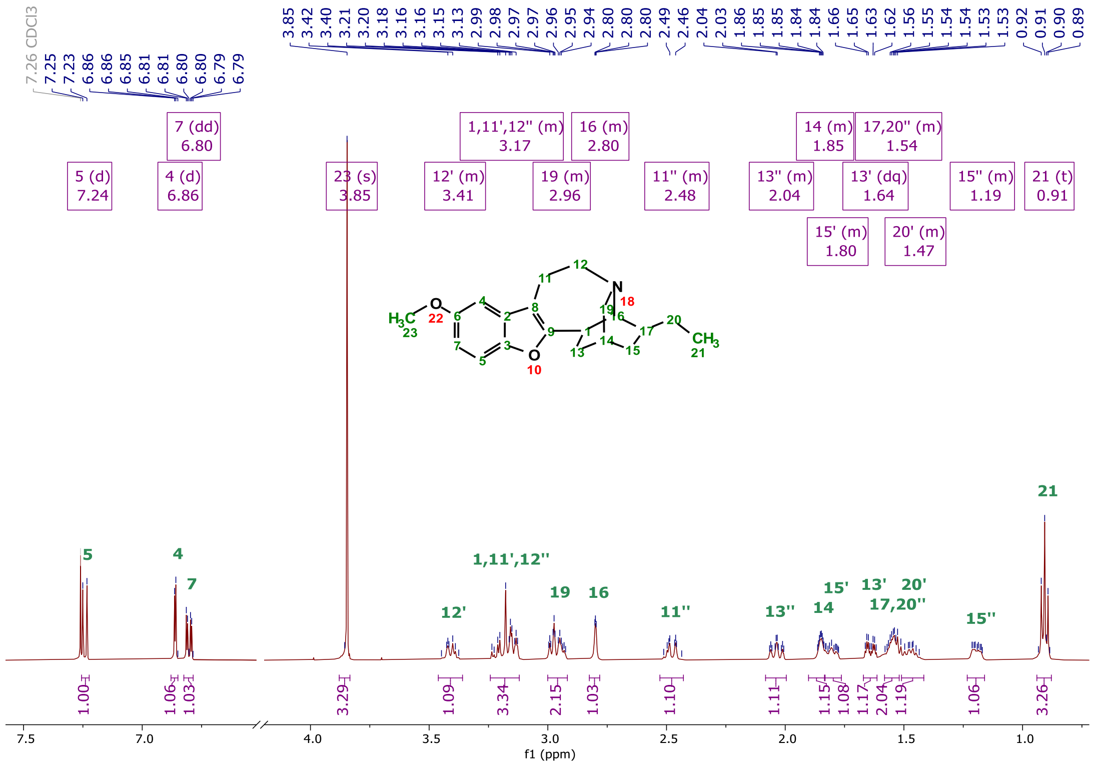

[1]H NMR (500 MHz, $CDCl_3$) magnified and assigned spectrum of compound **10a**. Blank regions of spectra were omitted for clarity.

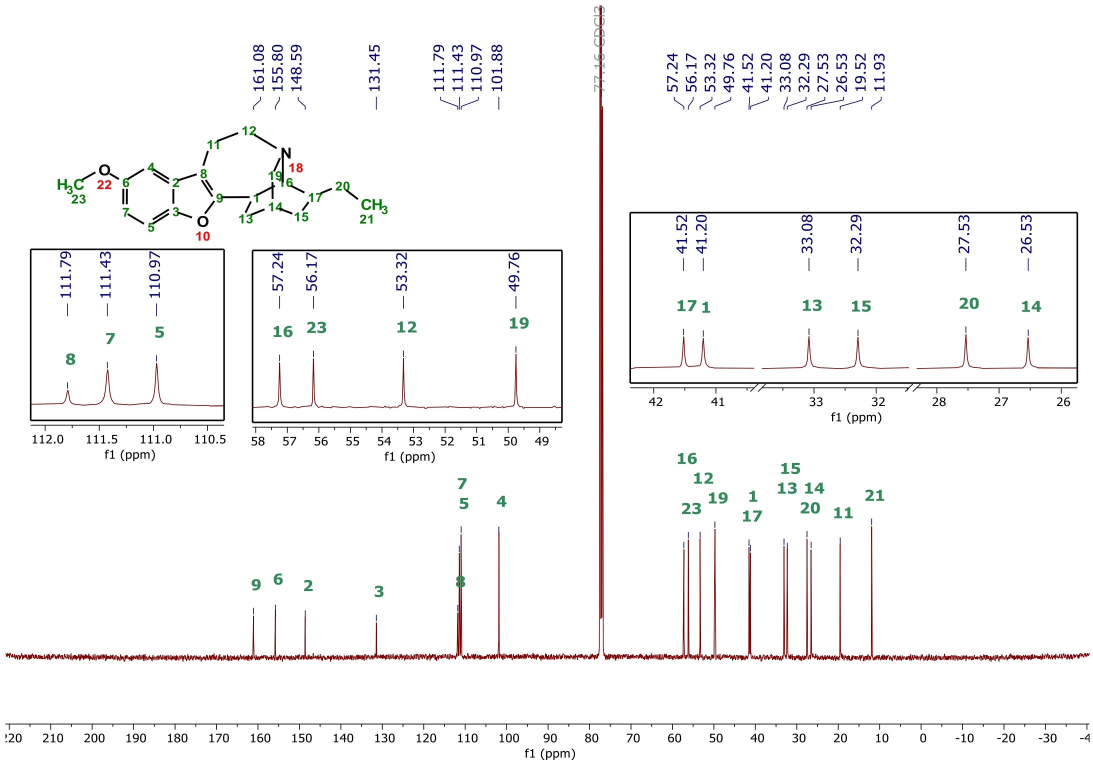


$^{13}C\{^{1}H\}$ NMR (126 MHz, $CDCl_3$) spectrum of compound **10a.** Expansions included for tightly clustered peaks.

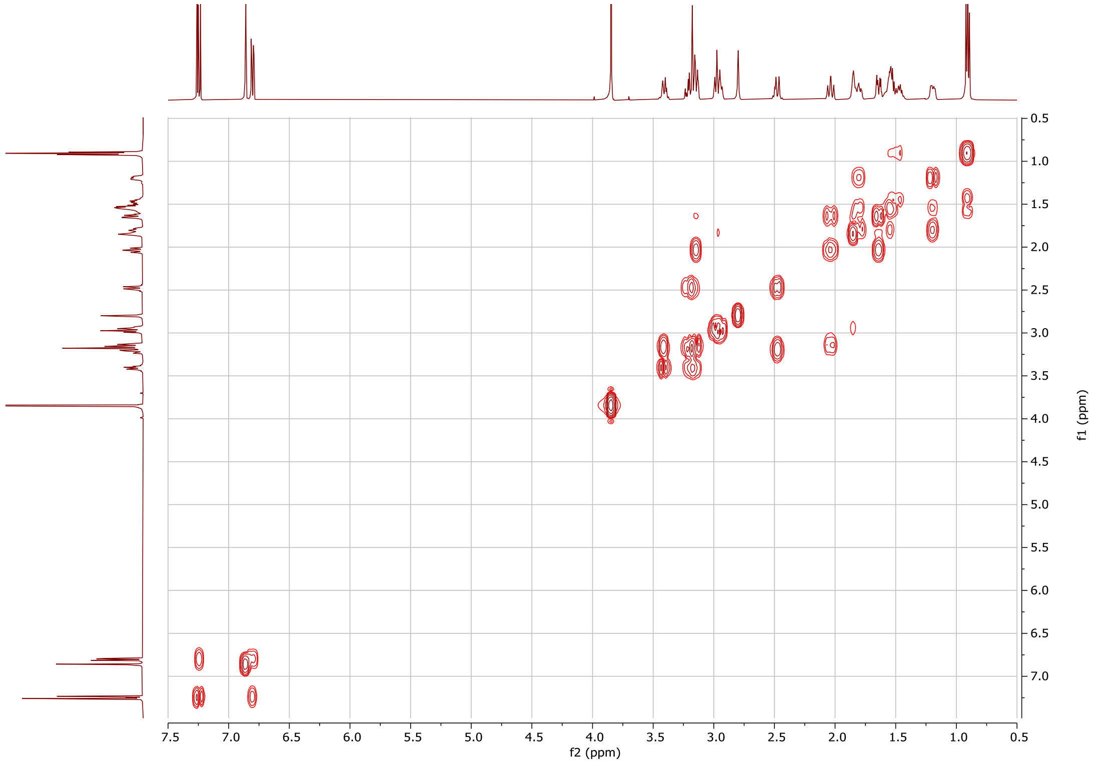


$^{1}H$-$^{1}H$ COSY NMR (500 MHz, $CDCl_3$) spectrum of compound **10a.**

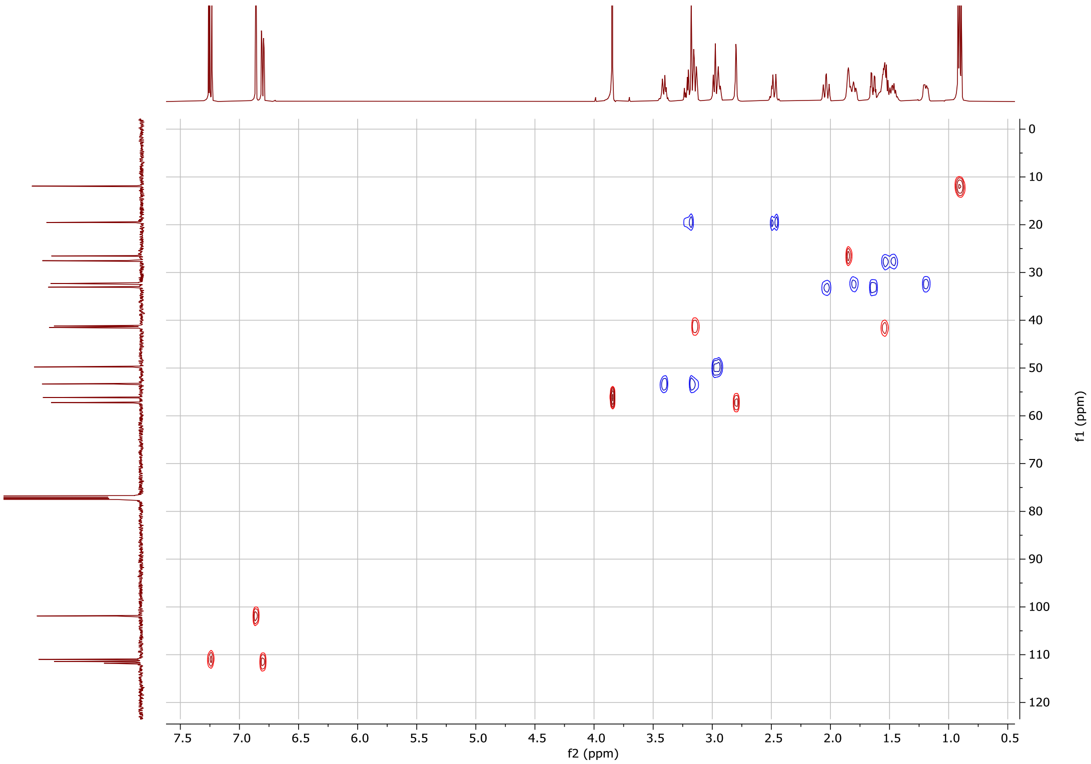


[1]H-[13]C HSQC multiplicity edited NMR (500/126 MHz, $CDCl_3$) spectrum of compound **10a.**

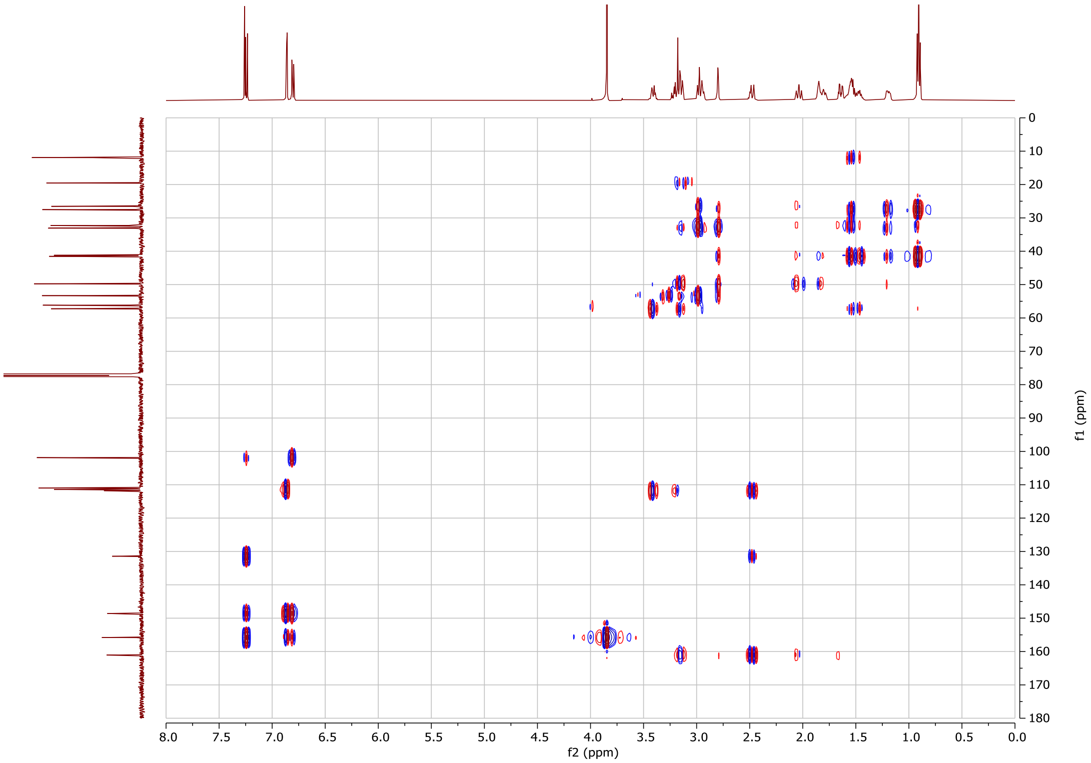


[1]H-[13]C HMBC NMR (500/126 MHz, $CDCl_3$) spectrum of compound **10a**, one-bond correlations are suppressed.

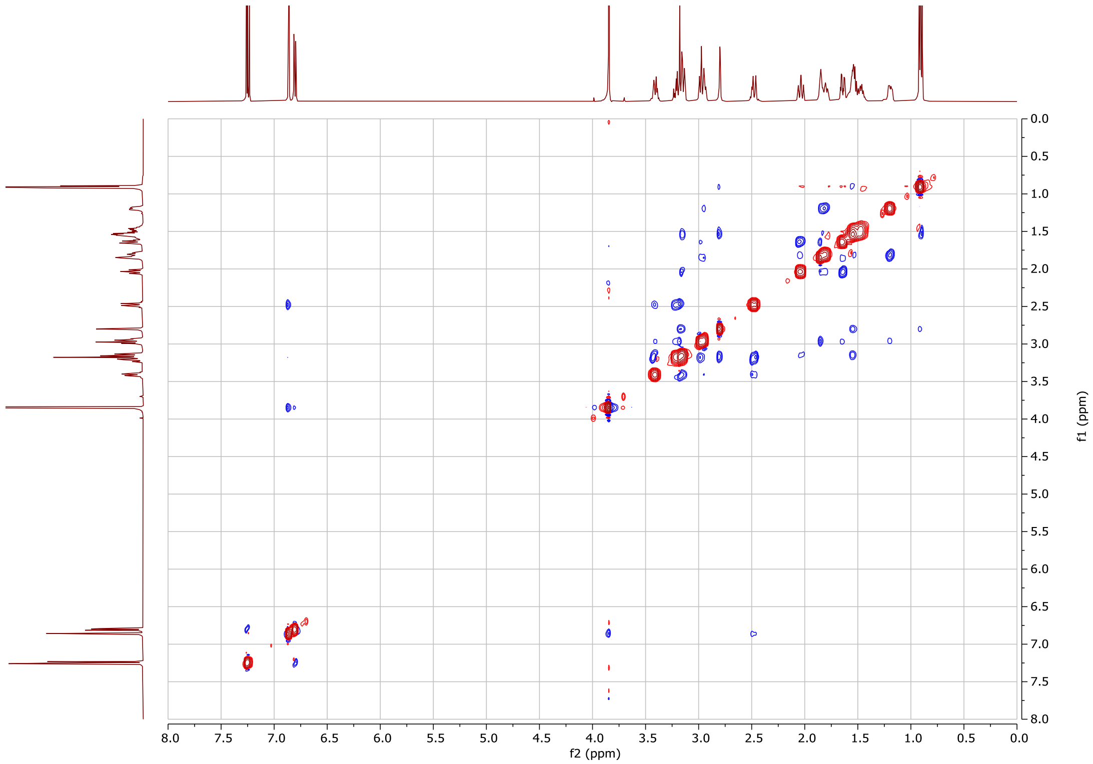

[1]H-[1]H NOESY NMR (500 MHz, $CDCl_3$) spectrum of compound **10a.**

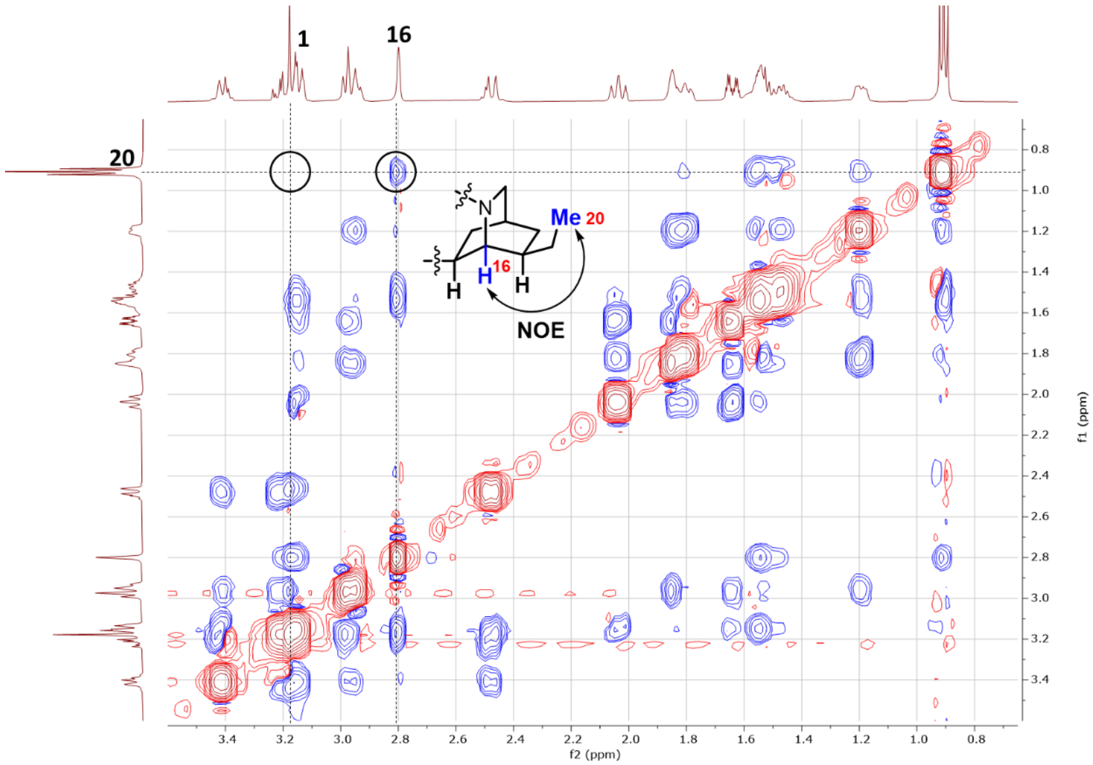

Isoquinuclidine portion of the NOESY spectrum of compound **10a.**

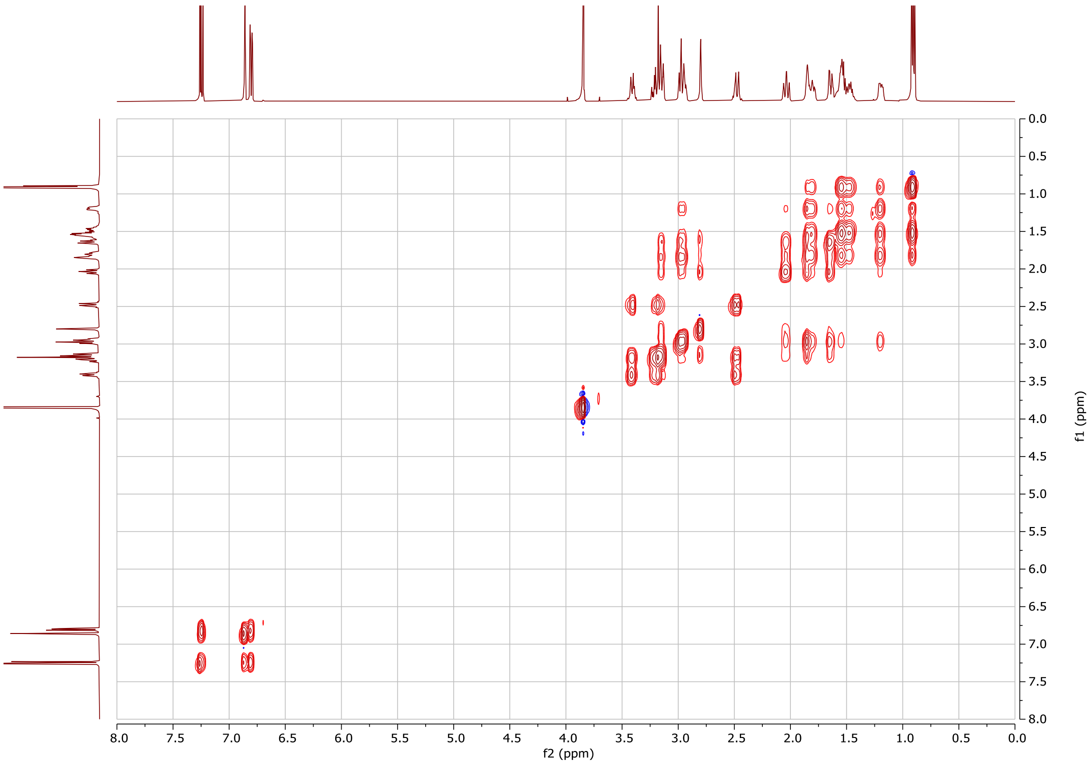


$^{1}H$-$^{1}H$ TOCSY NMR (500 MHz, $CDCl_3$) spectrum of compound **10a.**

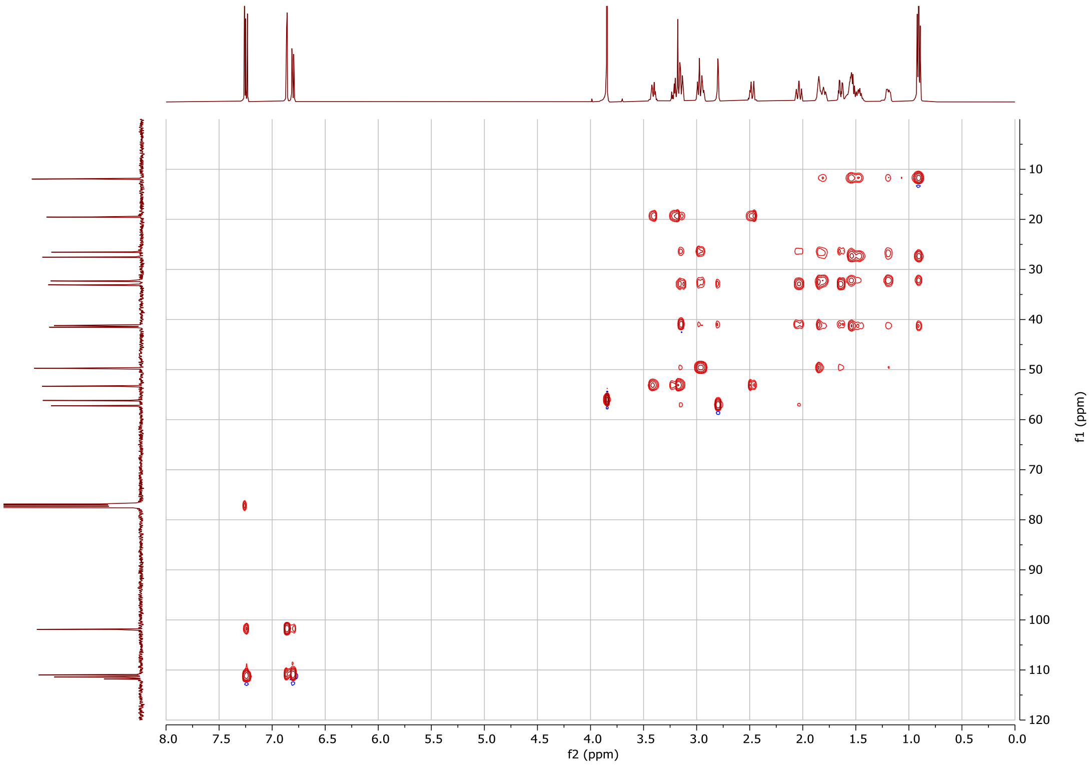


$^{1}H$-$^{13}C$ HSQC-TOCSY NMR (500/126 MHz, $CDCl_3$) spectrum of compound **10a**.

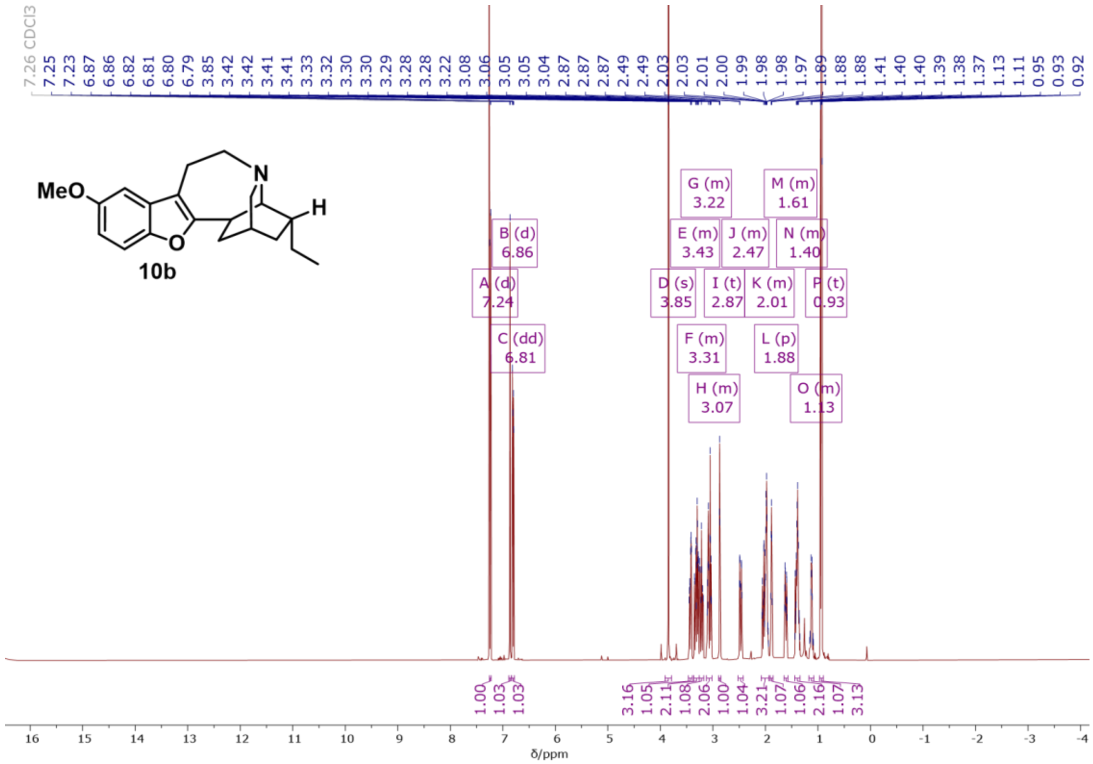


$^1$H NMR (500 MHz, $CDCl_3$) spectrum of compound **10b.**

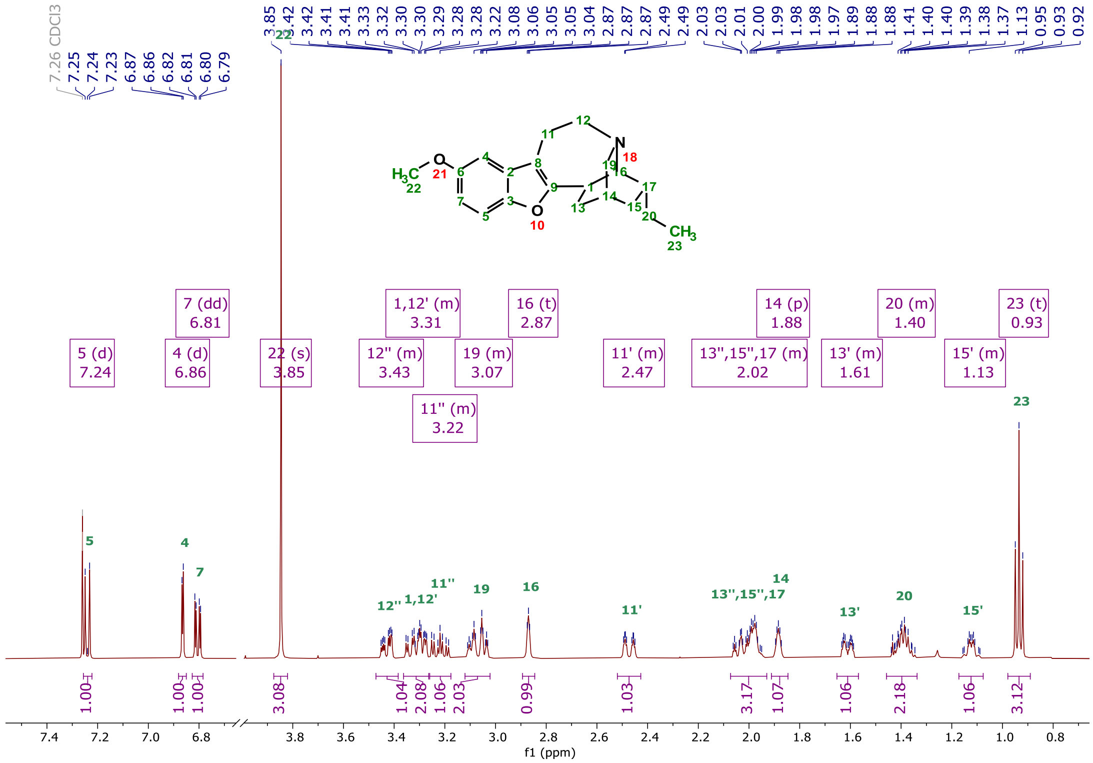


$^1$H NMR (500 MHz, $CDCl_3$) magnified and assigned spectrum of compound **10b**. Blank regions of spectra were omitted for clarity.

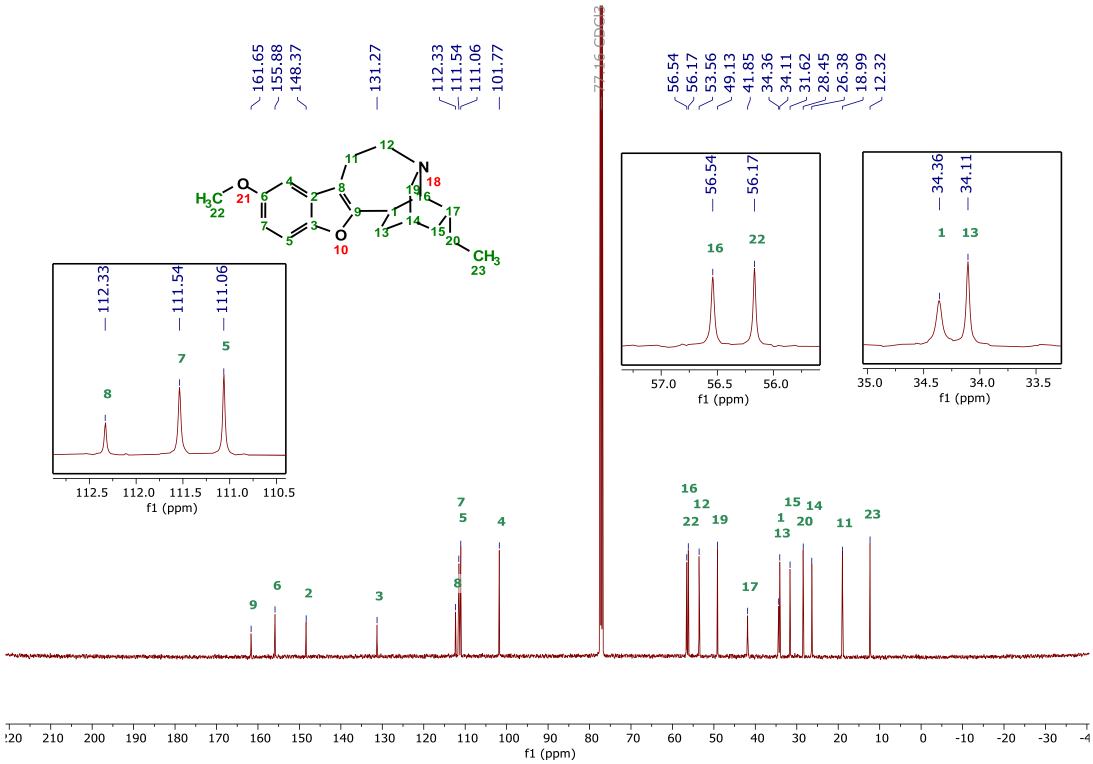

$^{13}C\{^{1}H\}$ NMR (126 MHz, $CDCl_3$) spectrum of compound **10b**. Expansions included for tightly clustered peaks.

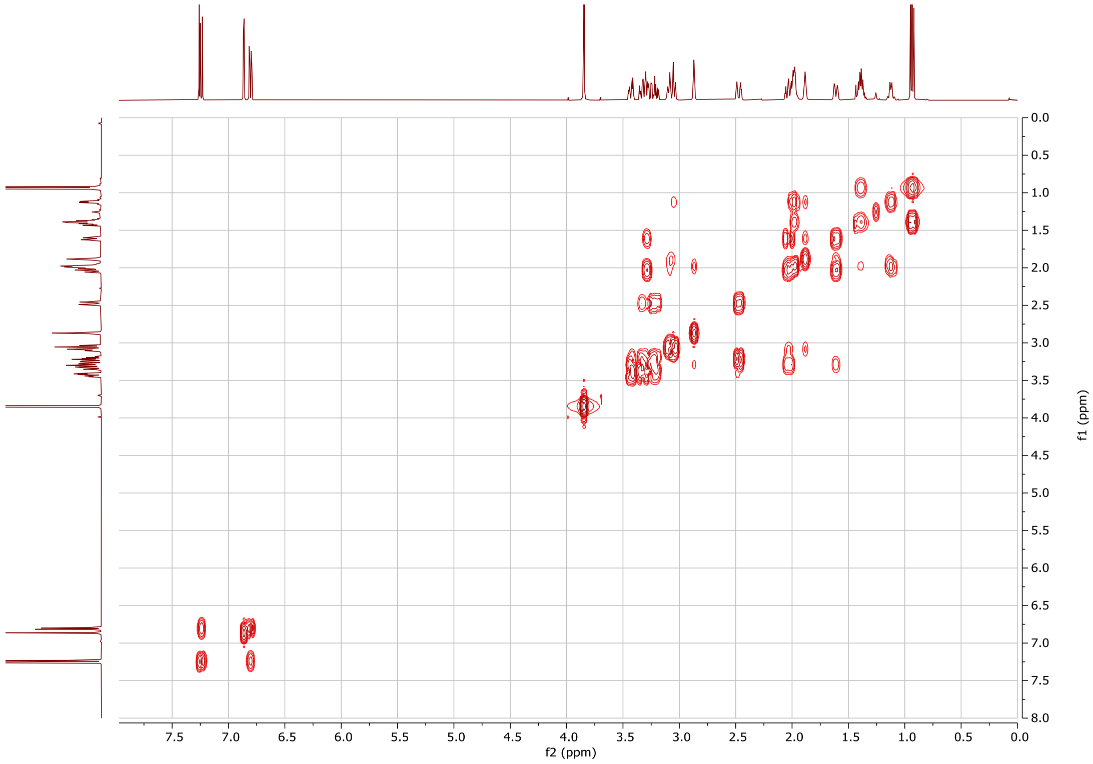

$^{1}H$-$^{1}H$ COSY NMR (500 MHz, $CDCl_3$) spectrum of compound **10b.**

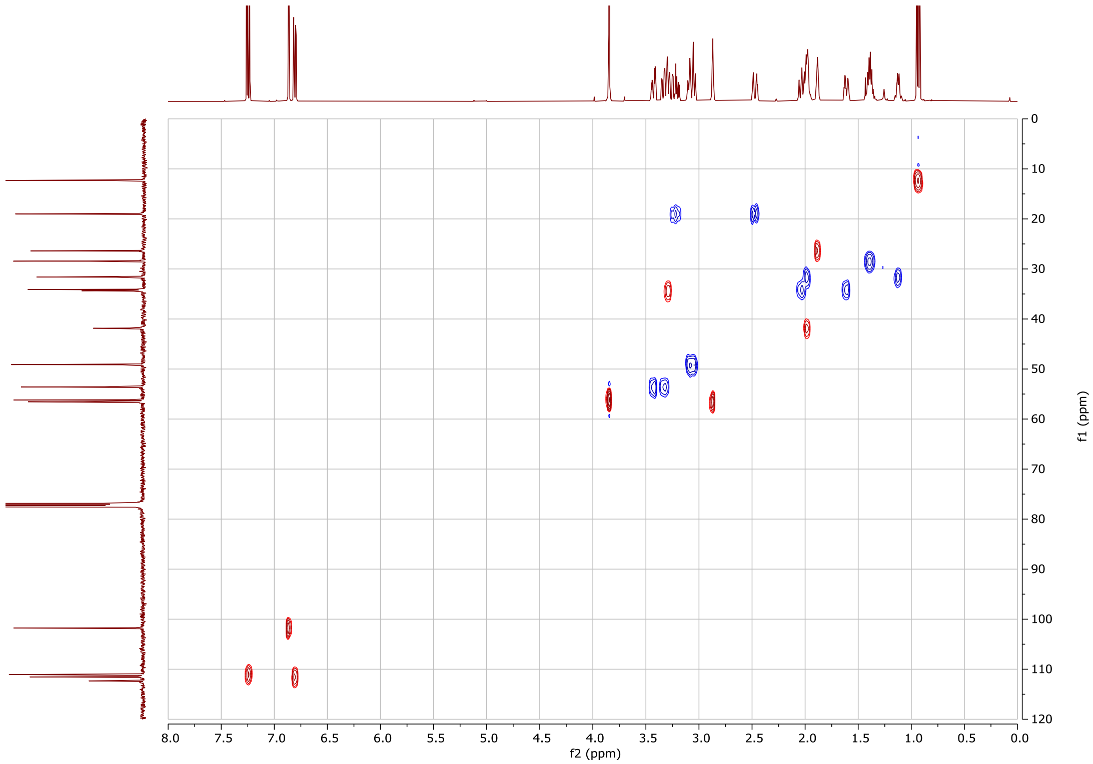


[1]H-[13]C HSQC multiplicity edited NMR (500/126 MHz, $CDCl_3$) spectrum of compound **10b.**

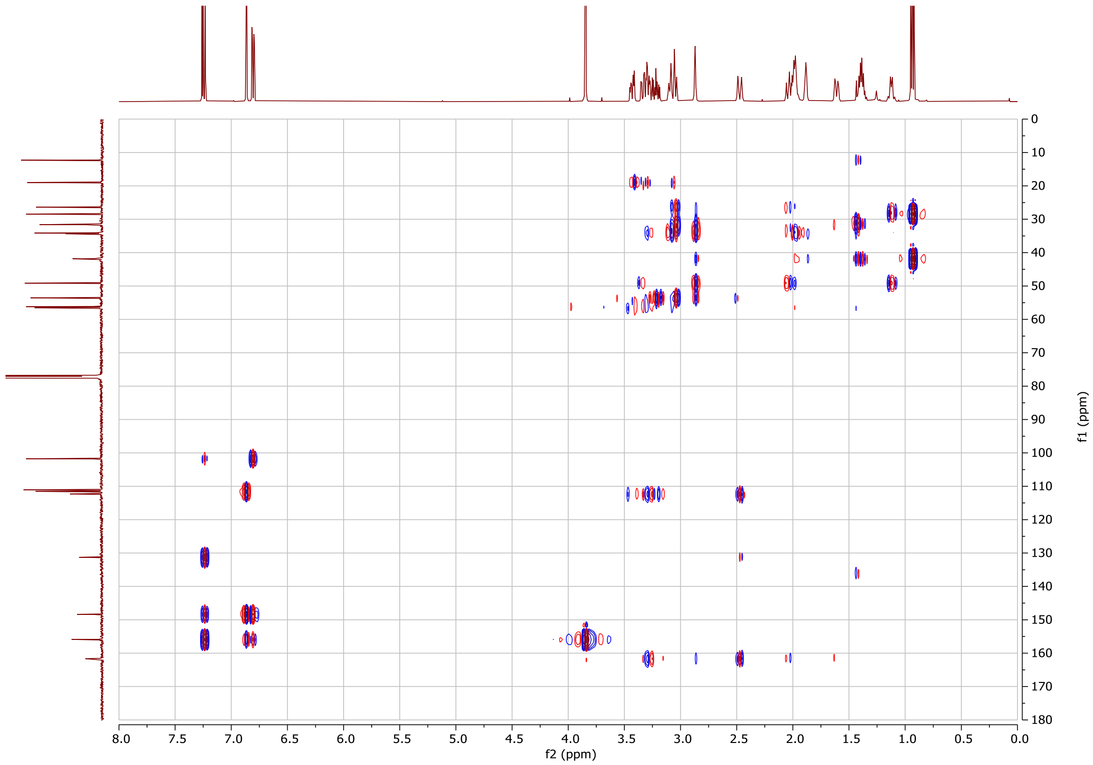


[1]H-[13]C HMBC NMR (500/126 MHz, $CDCl_3$) spectrum of compound **10b**, one-bond correlations are suppressed.

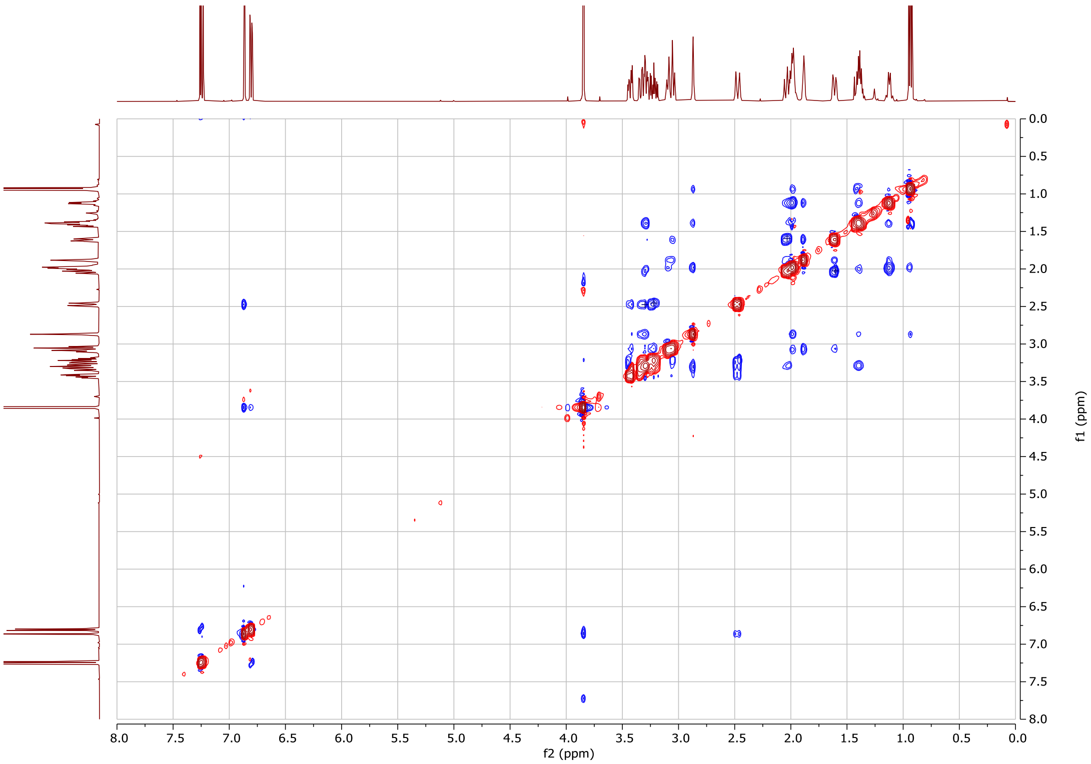


$^1H$-$^1H$ NOESY NMR (500 MHz, $CDCl_3$) spectrum of compound **10b.**

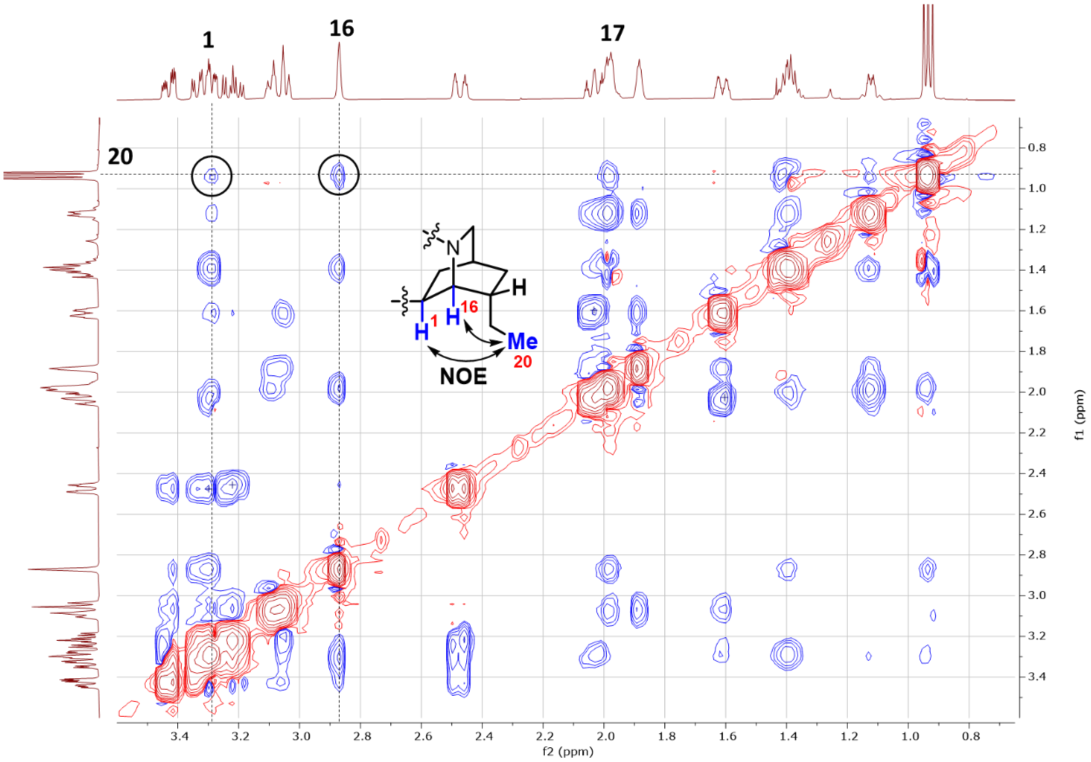


Isoquinuclidine portion of the NOESY spectrum of compound **10b.**

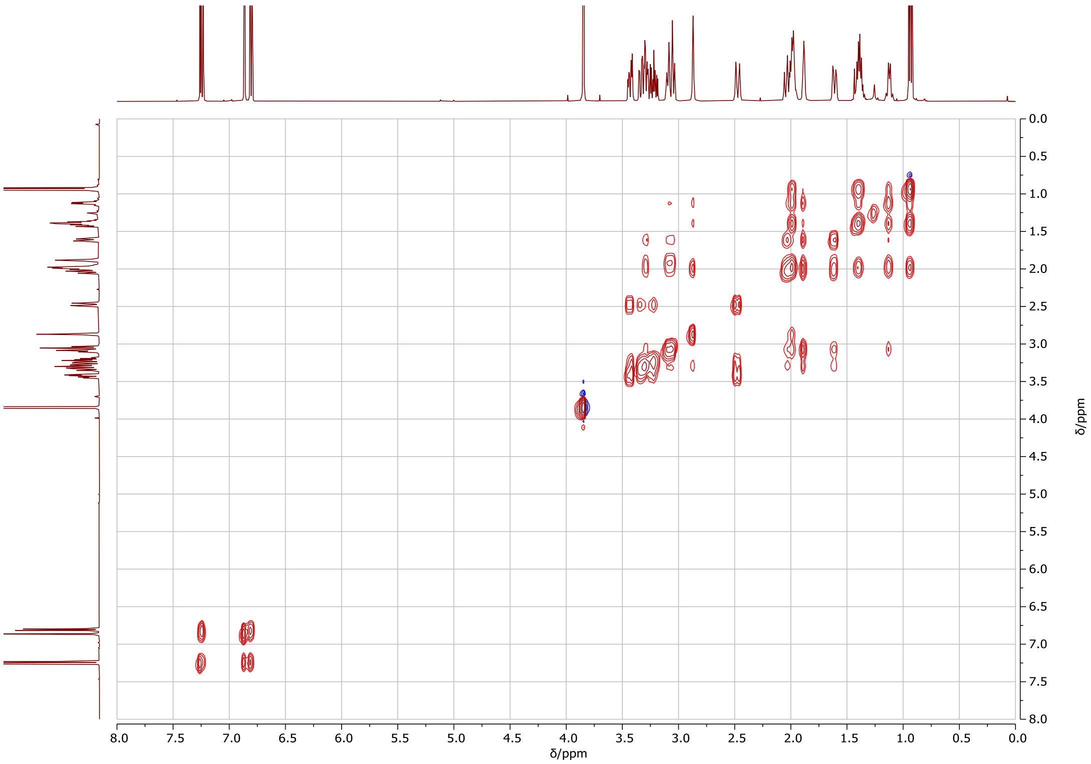


$^{1}H$-$^{1}H$ TOCSY NMR (500 MHz, $CDCl_3$) spectrum of compound **10b.**

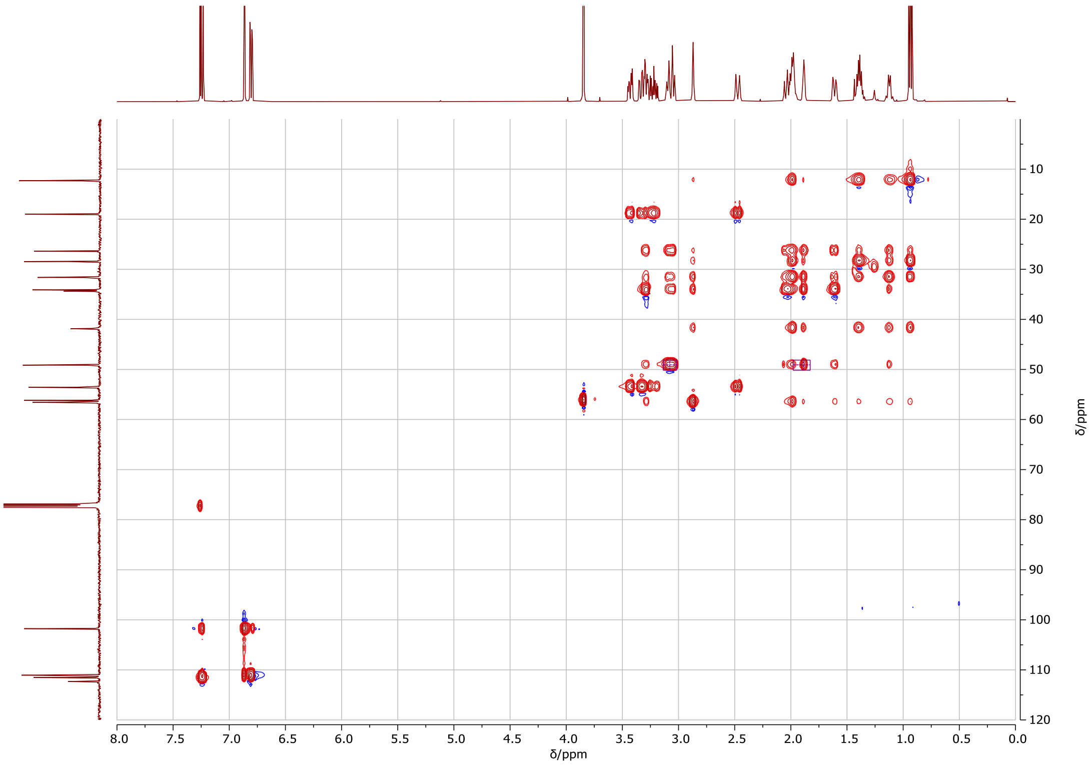


$^{1}H$-$^{13}C$ HSQC-TOCSY NMR (500/126 MHz, $CDCl_3$) spectrum of compound **10b**.

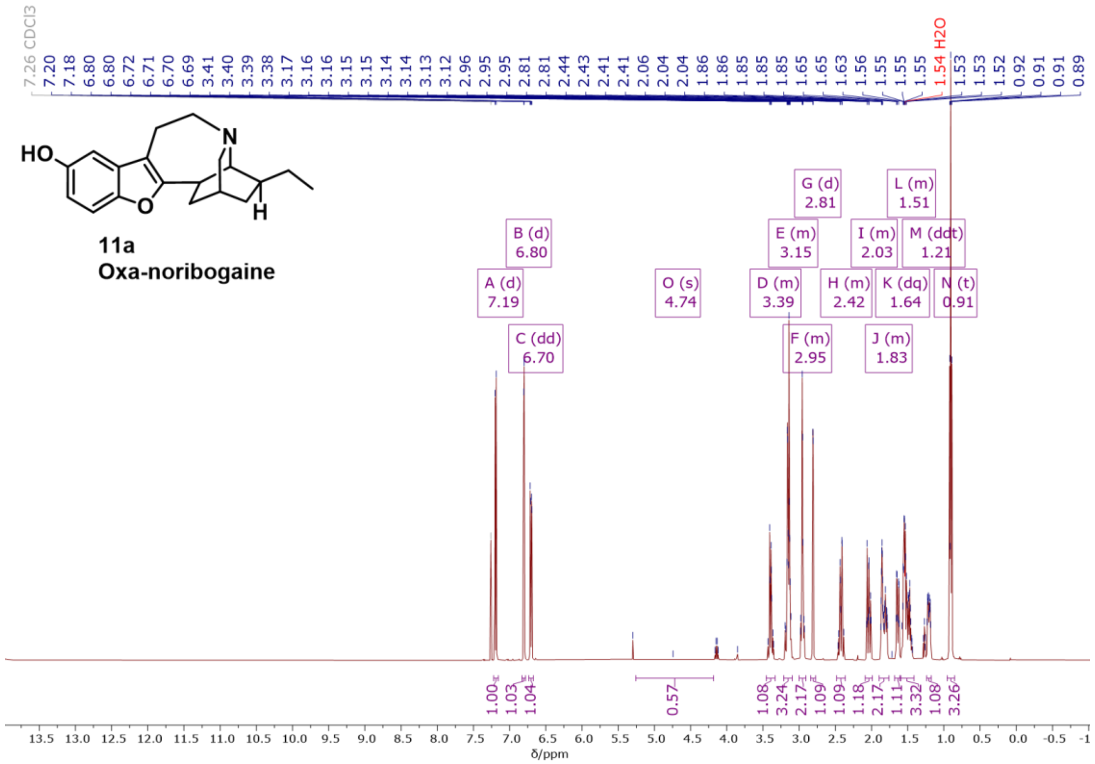

$^{1}H$ NMR (500 MHz, $CDCl_3$) spectrum of compound **11a.**

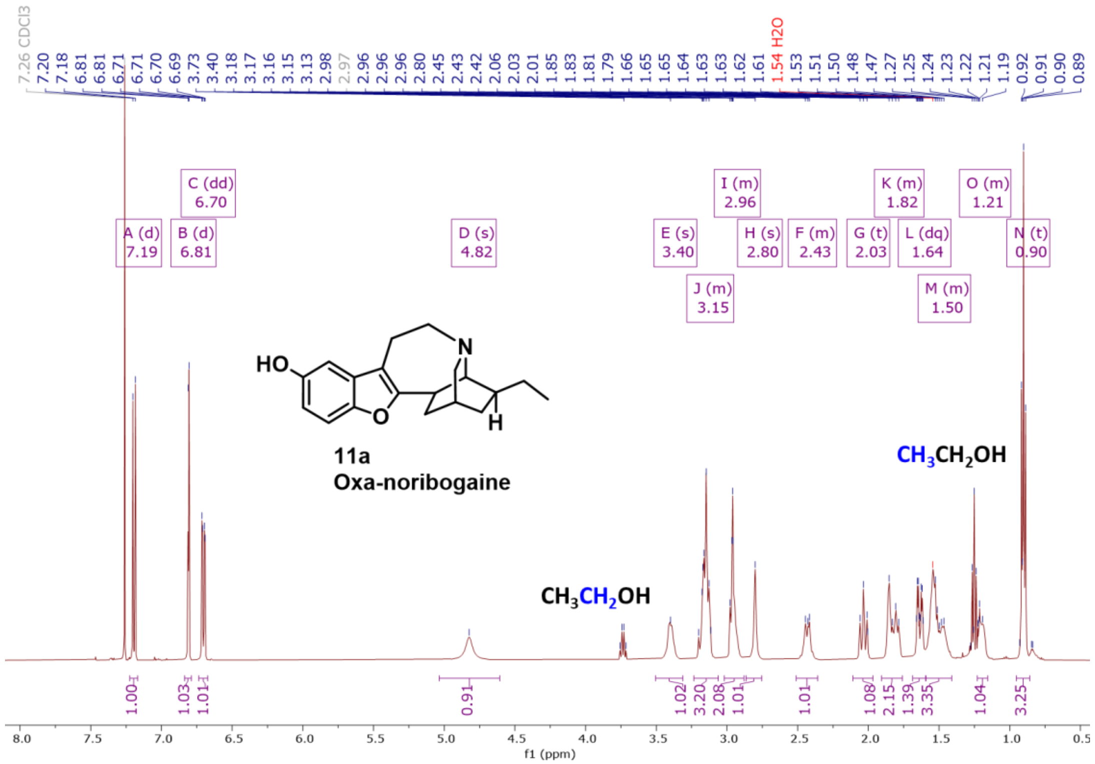

$^{1}H$ NMR (500 MHz, $CDCl_3$) spectrum of compound **11a**, multigram scale preparation. Residual solvent (EtOH) remains trapped in solid material even after extensive drying at elevated temperature (45 °C) in vacuum.

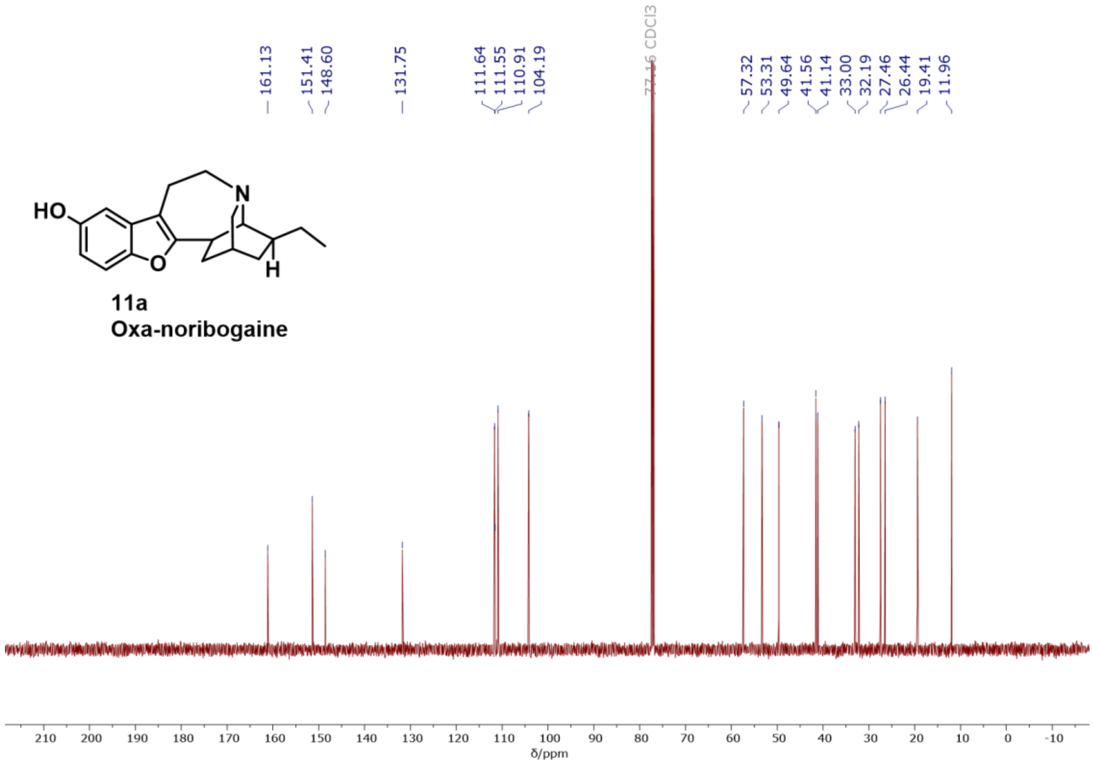

$^{13}C\{^{1}H\}$ NMR (126 MHz, $CDCl_3$) spectrum of compound **11a**.

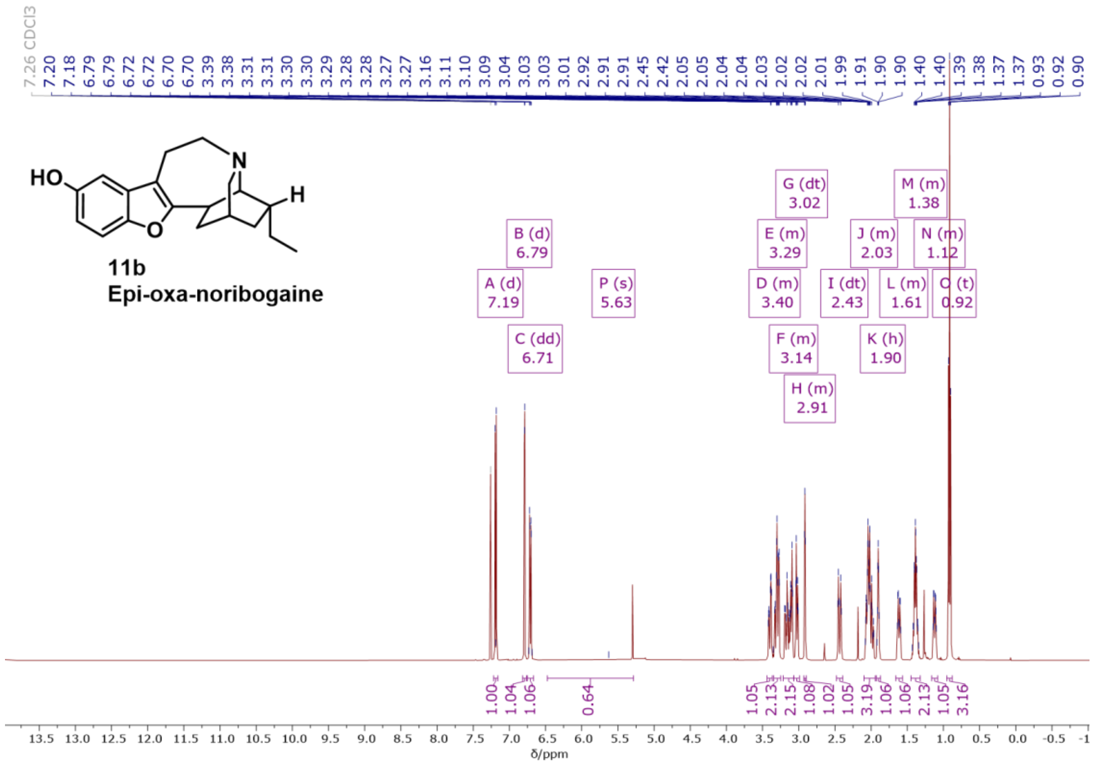

$^{1}H$ NMR (500 MHz, $CDCl_3$) spectrum of compound **11b.**

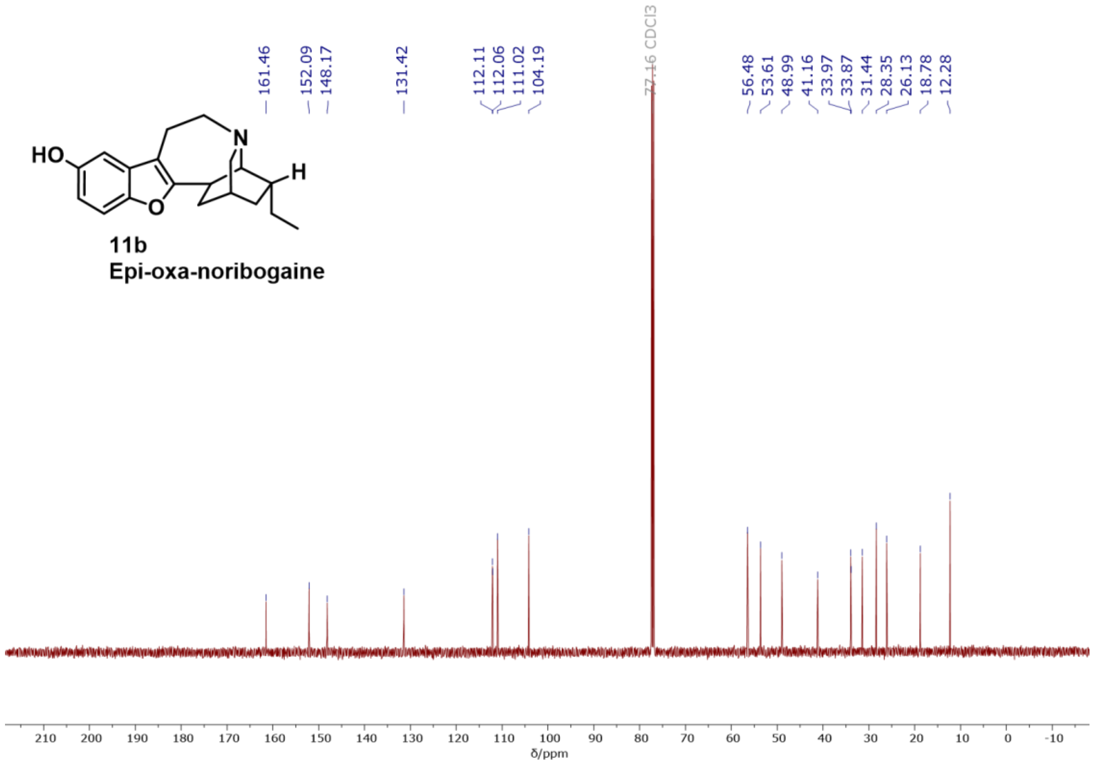

$^{13}C\{^{1}H\}$ NMR (126 MHz, $CDCl_3$) spectrum of compound **11b**.

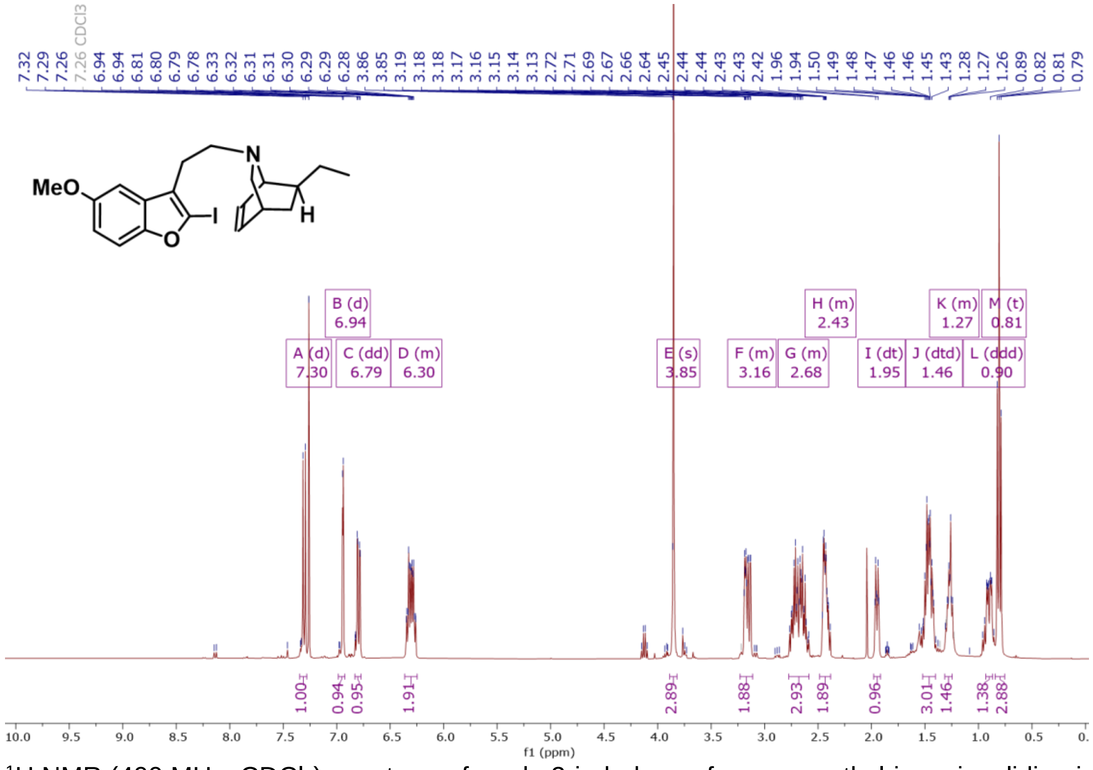

$^{1}H$ NMR (400 MHz, $CDCl_3$) spectrum of crude 2-iodo-benzofuran-*exo*-ethyl-isoquinuclidine intermediate**.**

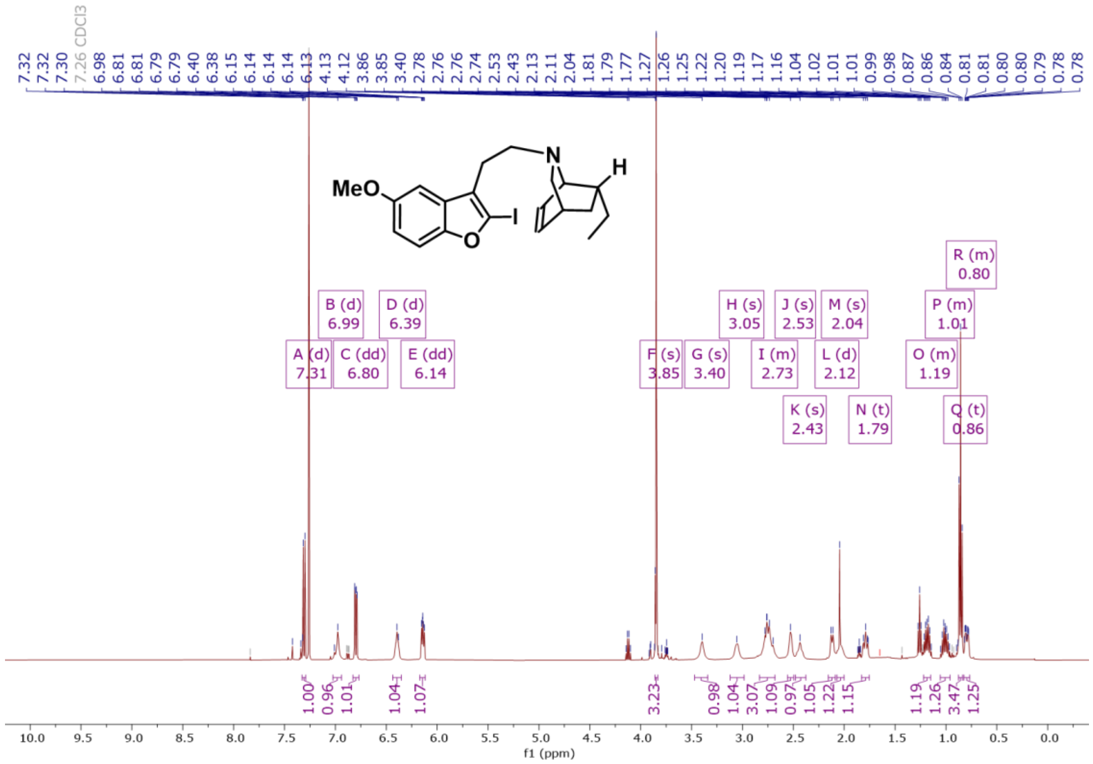


[1]H NMR (400 MHz, $CDCl_3$) spectrum of crude 2-iodo-benzofuran-*endo*-ethyl-isoquinuclidine intermediate**.**

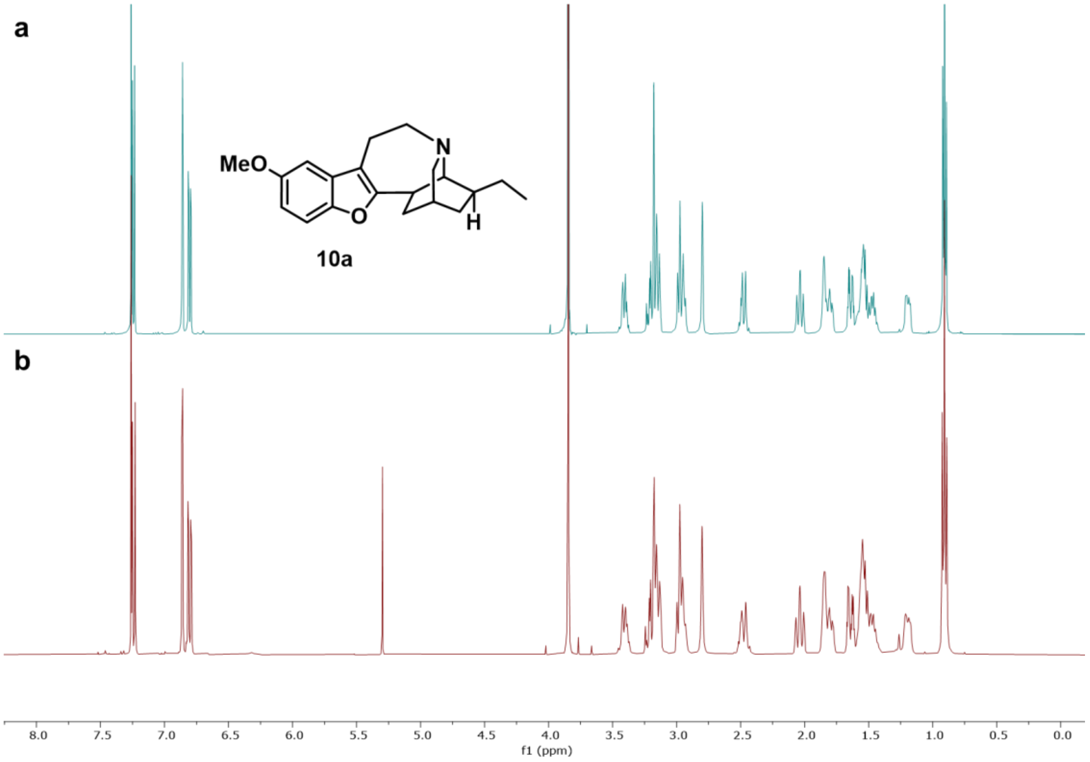


Stacked [1]H NMR spectra of oxa-ibogaine **10a** prepared using a) a Nickel mediated C-H coupling or b) reductive Heck procedure**.**

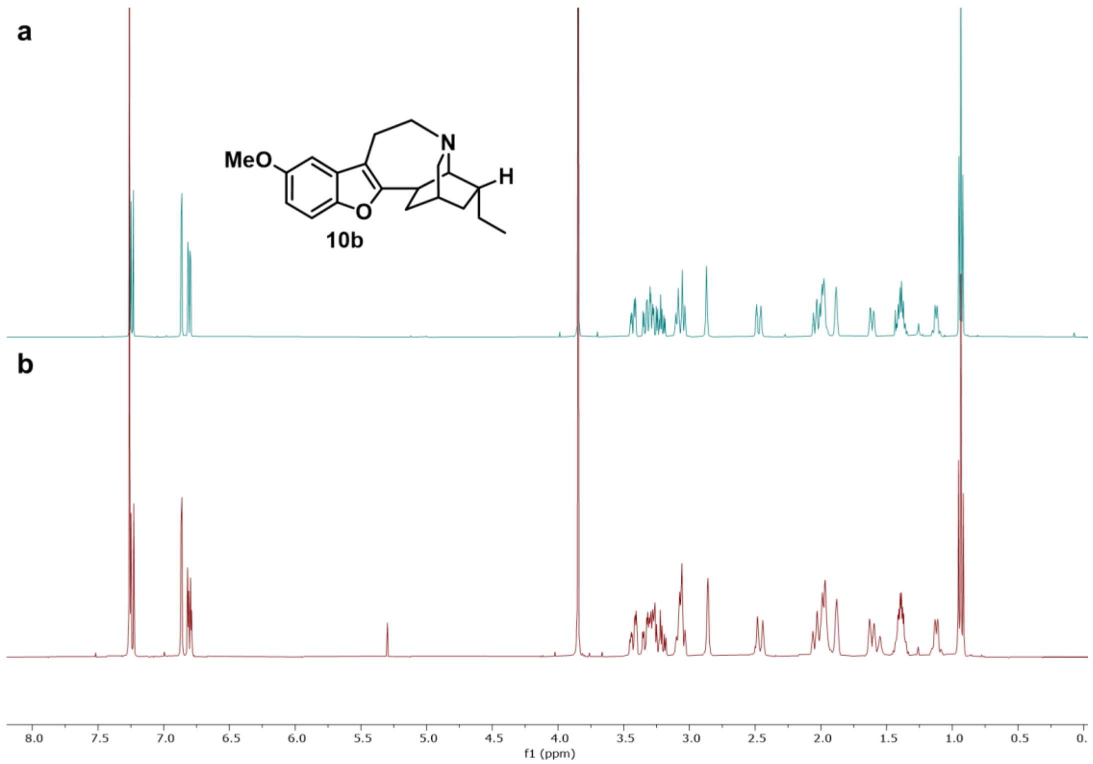


Stacked ¹H NMR spectra of epi-oxa-ibogaine **10b** prepared using a) a Nickel mediated C-H coupling or b) reductive Heck procedure**.**
